# Massively parallel pooled screening reveals genomic determinants of nanoparticle-cell interactions

**DOI:** 10.1101/2021.04.05.438521

**Authors:** Natalie Boehnke, Joelle P. Straehla, Hannah C. Safford, Mustafa Kocak, Matthew G. Rees, Melissa Ronan, Danny Rosenberg, Charles H. Adelmann, Raghu R. Chivukula, Namita Nabar, Adam G. Berger, Nicholas G. Lamson, Jaime H. Cheah, Hojun Li, Jennifer A. Roth, Angela N. Koehler, Paula T. Hammond

## Abstract

To accelerate the translation of cancer nanomedicine, we hypothesize that integrated genomic screens will improve understanding of the cellular processes governing nanoparticle trafficking. We developed a massively parallel high-throughput screening method leveraging barcoded, pooled cancer cell lines annotated with multi-omic data to investigate cell association patterns across a nanoparticle library spanning a range of formulations with clinical potential. This approach identified both the materials properties and cell-intrinsic features mediating nanoparticle-cell association. Coupling the data with machine learning algorithms, we constructed genomic nanoparticle trafficking networks and identified nanoparticle-specific biomarkers, including gene expression of SLC46A3. We engineered cell lines to validate SLC46A3 as a biomarker whose expression inversely predicts liposomal nanoparticle uptake both *in vitro* and *in vivo.* We further demonstrated the predictive capabilities extend beyond liposomal nanoparticles, regulating both uptake and transfection efficacy of solid lipid nanoparticles. Our work establishes the power of massively parallel pooled cell screens for nanoparticle delivery and enables the identification and utilization of biomarkers to rationally design nanoformulations for specific patient populations.

## Main Text

Nanoparticle (NP)-based therapeutics have enormous potential for personalized cancer therapy as they can encapsulate a range of therapeutic cargos including small molecules, biologics and, more recently, nucleic acids. Therapy-loaded NPs can be designed to prevent undesired degradation of the cargo, increase circulation time, and direct drugs specifically to target tumors.(*1–3*) There have been notable successes in clinical translation of nanomedicines, including liposomal formulations of doxorubicin (Doxil) and irinotecan (Onivyde®).(*4*) These formulations extend the half-life of the active agent and have the potential to lower toxicity, but do not efficiently accumulate in tumors.(*5, 6*)

Delivery challenges attributed to circulation, immune detection and clearance, as well as extravasation and diffusion through tissue all influence NP accumulation at target disease sites. Efforts to improve NP accumulation in tumors via active targeting motifs have been met with limited success, both in the laboratory and the clinic.(*1, 7*) Fewer efforts have focused on gaining a fundamental understanding of the biological features mediating successful NP-cell interaction and uptake. While progress has been made in understanding how specific physical and chemical NP properties affect trafficking and uptake, comprehensive evaluation of multiple NP parameters in combination has thus far been elusive. Additionally, the biologic diversity of cancer targets makes it prohibitively challenging to gain a holistic understanding of which NP properties dictate successful trafficking and drug delivery.(*8, 9*) Once NP parameters are considered in combination, the number of unique formulations to test increases exponentially, particularly as comparisons across several systems need to be drawn. A further barrier is the need to adapt the nanoparticle formulation of each encapsulated therapy for a given drug or target, as each formulation has its own unique biological fate.(*9*) As therapies continue to increase in molecular complexity, new nanocarrier formulations capable of delivering such entities will need to be developed and examined for their unique trafficking properties.

We and others have designed panels of NPs to elucidate the structure-function relationships to cellular targeting and uptake.(*10–13*) However, there is a need to equally consider the influence of biological heterogeneity on interactions at the NP-cell interface, for example by probing cells across cancer cell lineages with a range of genetic drivers and cell states. In the era of precision medicine, with the desire to deliver molecularly targeted and gene-based therapies to specific subcellular compartments within cancer cells, it is imperative to holistically probe the structure-function relationship of NPs as they relate to cellular interactions.

Inspired by recent advancements in cancer genomics,(*14*) we postulated that applying similar techniques to the study of cancer nanomedicine would uncover both the cell- and NP-specific features mediating efficient targeting and delivery. The combination of pooled screening with multi-omic annotation has accelerated target discovery and uncovered previously unrecognized mechanisms of action in small molecule screens. Specifically, in the Profiling Relative Inhibition in Mixtures (PRISM) method, DNA-barcoded mixtures of cells have recently been used for multiplexed viability screening. In cell line pools grouped by doubling time, 500 barcoded cell lines have been screened against tens of thousands of compounds to identify genotype-specific cancer vulnerabilities.(*15, 16*)

To comprehensively capture pan-cancer complexities and enable the statistical power to link NP association with cell intrinsic characteristics, we developed a competitive phenotypic screen to assess associations of a curated NP library across hundreds of cancer cell lines simultaneously. By pooling and plating 488 DNA barcoded cancer cell lines in a single well, we screened the interactions of a range of NP formulations with varied core compositions, surface chemistries, and diameters. We observed that NP core composition has a dominating influence on cell-specific interactions of the studied parameters. Coupling our biomarker findings with k-means clustering, we constructed genomic interaction networks associated with NP engagement, enabling the identification and connection of genes associated with the binding, recognition, and subcellular trafficking of distinct NP formulations. Moreover, through the use of univariate analyses and random forest algorithms, we identified that the gene *SLC46A3* holds significant value as a predictive, NP-specific biomarker. We further validated SLC46A3 as a negative regulator of liposomal NP uptake *in vitro* and *in vivo*. The strategy outlined herein identifies cellular features underlying nanoparticle engagement, adding a new dimension to the study of cancer nanomedicine.

### nanoPRISM: screening nanoparticle association with pooled cell lines

To screen hundreds of cancer cell lines simultaneously for NP-cancer cell line association patterns, we cultured pooled PRISM cells and incubated them with fluorescent NPs. We then implemented a fluorescence-activated cell sorting (FACS) adaptive gating strategy to sort cell populations into four bins (quartiles, A-D) based on fluorescence signal as a proxy for the extent of NP-cell association (Figure 1A). Experimental parameters were optimized to ensure sufficient cell number and barcode representation post-cell sorting (Figure S1) and NPs were incubated for 4 and 24 hours.

**Figure 1.**
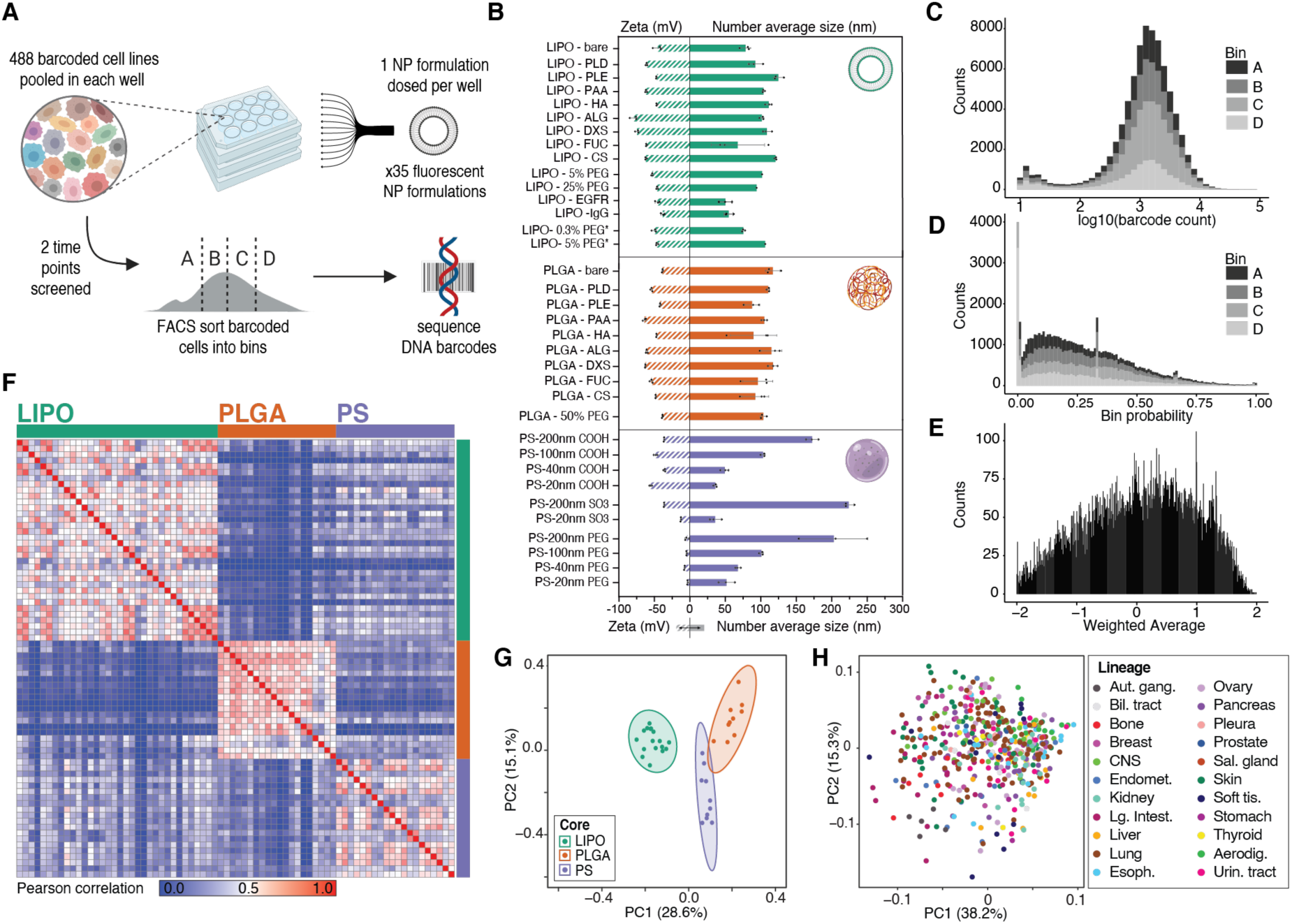
Assessing NP-cell interactions across hundreds of cancer cell lines simultaneously. (**A**) Schematic of the nanoPRISM assay: Fluorescently-labeled NPs are incubated with pooled cancer cells before fluorescence-activated cell sorting (FACS) by NP-association and sequencing of DNA barcodes for downstream analyses. (**B**) Characterization of the diameter and zeta potential of the NP library via dynamic light scattering. Data is represented as the mean and standard deviation of three technical repeats. Formulations marked with an asterisk represent drug-free analogs of clinical liposomal formulations as described in the text. (**C**) Raw data from the screen was obtained in the form of barcode counts, with similar numerical distribution of barcodes in each bin, represented as a stacked histogram. (**D**) Accounting for baseline differences in barcode representation yields the probability (P) that each cell line will be found in a particular bin. (**E**) Probabilities are collapsed into a single weighted average (WA) for each NP-cell line pair. (**F**) A similarity matrix collapsing WA values for 488 cell lines reveals clusters of NP formulations with the same core formulation. (**G-H**) Principal component analysis (PCA) of NP-cell line WA values at 24 h confirms distinct clustering of NP formulations based on core composition but cell lines do not form clusters, indicating lineage does not significantly influence NP-cancer cell interactions.

For this screen, we designed a modular NP library to capture the effects of NP core composition, surface chemistry, and size on cell interactions. This panel of 35 NPs encompassed both clinical and experimental formulations. Specifically, anionic liposomes were formulated and electrostatically coated with cationic poly-L-arginine (PLR) followed by a series of polyanions.(*17–21*) The polyanions were selected for their synthetic (polyacrylic acid, PAA), semisynthetic (poly-L-aspartate, PLD; poly-L-glutamate, PLE), or natural (hyaluronate, HA; dextran sulfate, DXS; fucoidan, FUC; alginate, ALG; chondroitin sulfate, CS) origin as well as the inclusion of both carboxylate and sulfate ions.(*22–24*) These same electrostatic coatings were used to modify polymeric NP cores (polylactide-*co*-glycolide, PLGA) to test the effects of core composition on NP-cell interactions. We optimized formulations to obtain a diameter of approximately 100 nm for the liposome and PLGA formulations as the similar sizes would enable cross-core comparisons. We also included commercially manufactured fluorescent carboxylate- and sulfate-modified polystyrene (PS) nanoparticles in a range of diameters from 20-200 nm, enabling study of particle size and surface chemistry. Because of the clinical importance of polyethylene glycol (PEG)-containing formulations,(*25*) PEGylated versions of liposome, PLGA, and PS particles were prepared, including the drug-free versions of two commercial formulations, liposomal doxorubicin (Doxil) and liposomal irinotecan (Onyvide®). The latter two formulations are denoted as LIPO-5% PEG* and LIPO-0.3% PEG*, respectively. All of the nanoparticles examined exhibited negative or neutral net charge, as the focus of this work is on systemic nanoparticle delivery systems. Positively charged nanoparticles have been shown to undergo nonspecific charge interactions with cells and proteins, leading to toxicity and premature clearance *in vivo*.(*26*) Dynamic light scattering (DLS) was used to characterize the diameter, zeta potential, and polydispersity index (Figure 1B, Tables S1-S2) of this NP library.

To ensure that our methods led to robust and meaningful data we selected an anti-epidermal growth factor receptor (EGFR) antibody as an active targeting control. We hypothesized that the design of our screen would allow us to identify features relevant to EGFR expression with a high level of confidence. A nonlethal EGFR antibody or IgG isotype control was covalently incorporated onto a liposome via a PEG tether.(*27*) We elected to focus on EGFR due to the wide range of native EGFR expression of the 488 cell lines included in our screen as well as prior evaluation of EGFR-targeting compounds via the PRISM assay (Figure S2).(*15*)

After incubation with the NP library, we utilized fluorescence-activated cell sorting to bin cells into quartiles according to fluorescence intensity (Figure S3). Cells were then lysed, and the DNA barcodes were amplified, sequenced, and deconvoluted according to previously detailed protocols.(*15, 28*) After quality control analysis of technical (n=2) and biologic (n=3) replicates, all 488 cell lines met quality control measures and were carried forward for downstream analyses (Figure S4, Supplementary Text). This dynamic gating strategy was used to enable comparison of cell line representation per bin (quartile) independent of fluorophore identity or amount incorporated into each tested formulation.

A probabilistic model was developed and applied to the data to infer the relative distribution of each cell line into the pre-determined bins (A-D) for each NP formulation. The probability of a cell from a given cell line falling into a given bin is used to represent those distributions, i.e., P_A_+ P_B_+ P_C_+ P_D_ = 1 (Figure 1C-D). The technical details and the model’s implementation are presented in the Supplementary Text section. Given the concordance of the inferred probabilities among the biologic replicates (Figure S5), we collapsed the replicates through their arithmetic average. Probabilities were then summarized using a weighting factor alpha (*α*) to calculate a weighted average (WA) for each NP-cell line pair: WA = -*α*P_A_-P_B_+P_C_+*α*P_D_ in which a higher WA implies higher NP-cell association and vice versa (Figure 1E). We trialed a range of weighting factors (*α* = 2, 10, 20 and 100) and found that downstream results were unchanged with the higher *α* values (Figure S6), and therefore, *α* = 2 was used for subsequent analyses.

### Cancer cells distinguish nanoparticles based on core composition

Pearson-based unsupervised hierarchical clustering of pairwise WAs identified NP core material as a strong determinant of cell association, with the three core materials tested (liposomal, PLGA and PS) forming distinct clusters (Figure 1F and S7A). This result was unexpected as we hypothesized surface chemistry to be a larger predictor of NP-cell interactions. Principal component analysis (PCA) similarly identified core specific trends at both the 4 and 24 hour time points (Figures 1G and S7B, D). Further analysis within each core material did reveal surface chemistry dependent trends, though they were more subtle than core-based clustering (Figure S8).

In contrast, no clusters were apparent when PCA was performed based on cell line, indicating that cancer cells of the same lineage did not have similar NP-association trends (Figure 1H, Figure S7C, E). Heterogeneity in NP-cell association in proliferating cells has been attributed to various aspects of cell growth and metabolism.(*29–32*) To ensure that differential cell proliferation did not confound our results, we performed a parallel growth experiment with the same pooled cells and found no correlation between estimated doubling time and WA (Figure S9).

### Cell-intrinsic features mediate nanoparticle trafficking

We applied data from the Cancer Cell Line Encyclopedia (CCLE)(*33, 34*) to identify genomic features that act as predictive biomarkers for NP-cell association. To do this, we employed both univariate analyses and a random forest algorithm to correlate the baseline molecular features of each cell line (cell lineage; gene copy number; messenger RNA, microRNA, protein or metabolite abundance; function-damaging, hotspot or missense mutations) with NP association.

#### EGFR-targeting compounds identify relevant biomarkers with high confidence

Using univariate analysis for all CCLE features, we identified EGFR gene expression and protein abundance as the two most significantly correlated hits (q = 4x 10^-100^ and q= 4×10^-76^, respectively) with anti-EGFR antibody, but much less significantly (q = 6 x 10^-9^ and q = 4 x 10^-10^, respectively) associated with the isotype control (Figure 2A, top panels). We also confirmed that fluorophore identity does not impact biomarker identification, demonstrating that both AlexaFluor 488 and Cy5 conjugated anti-EGFR antibodies perform similarly (new Figure S10).

**Figure 2.**
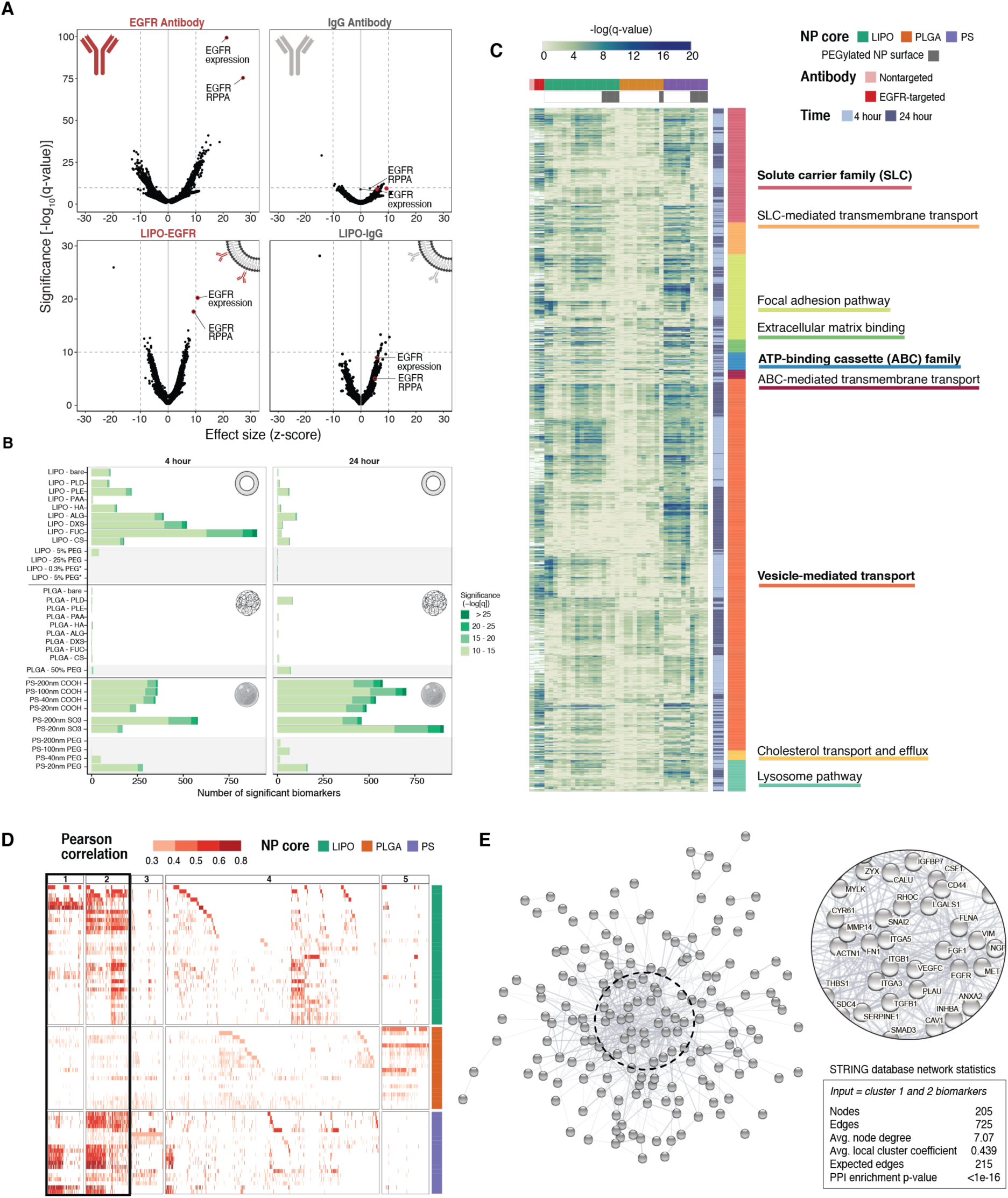
Correlative genomic analysis identifies expected validation biomarkers as well as hundreds of formulation- and time-dependent biomarkers. (**A**) Univariate analysis reveals EGFR gene expression and protein abundance (via reverse phase protein array; RPPA) to be strongly and positively correlated with high anti-EGFR association (top left). EGFR-related markers are much less significant in the isotype control (top right). The same EGFR-related hits, in addition to NP specific markers, are observed for antibody-conjugated liposomes (bottom row). (**B**) Univariate analysis identifies genomic features correlated with NP association. All biomarkers meeting a significance threshold of -log10(q-value) >10 are shown as stacked bar graphs separated by NP formulation and time point. PEGylated NP formulations are highlighted with a gray background. (**C**) A heatmap showing the significance of biomarkers associated with established transport, uptake, and adhesion gene sets. Targeting and non-targeting antibodies are included as references. Gene set headings are bolded and subsections are listed below respective headings. (**D**) A heatmap showing all gene- and protein-expression features with positive correlation identified by random forest algorithm in columns, and NP formulations in rows. Features are colored based on their Pearson correlation and clustered using k-means clustering, with clusters 1+2 highlighted as features present across multiple NP formulations. (**E**) Visual representation of the STRING network generated by inputting the 205 features from clusters 1+2, with network statistics. Each node represents a feature, and the edges represent predicted functional associations. The most interconnected nodes are labeled in the zoomed inset

In EGFR-conjugated liposomes, the same hits were also identified more significantly (q=6×10^-21^ and q=2×10^-18^, respectively) than the IgG control (q = 3 x 10^-9^ and q = 3 x 10^-6^, respectively) (Figure 2A, bottom panels).

The statistical significance of EGFR biomarkers was lower for the antibody-conjugated liposome than the free antibody, which may be due to steric blockage introduced by covalently linking an antibody to a NP surface that may interfere with binding to its target.(*35*) Thus, we demonstrated the ability to quantitatively compare expected biomarker targets of both free antibodies and antibody-conjugated NPs using our platform. This method of analysis will provide therapeutic insights in the design of antibody-drug conjugates, specifically in evaluating the effects of conjugation site or linker chemistry.

#### Biomarker number and significance are influenced by nanoparticle properties

We employed univariate analysis to correlate association and CCLE features for each NP formulation, both quantitatively and qualitatively using curated gene sets. First, we thresholded q-values at less than 1×10^-10^ to compare the absolute number of candidate biomarkers at varying degrees of significance (Figure 2B). Selection of this cutoff was guided by the IgG-conjugated antibody analysis, which returned few hits above this threshold. For liposomal NPs, we observed that the number of significant biomarkers was higher at 4 h than 24 h. We believe this may be indicative of active uptake processes, established to take place within the first few hours of NP-cell interactions, whereas at 24 hours, we may be capturing features associated with less specific interactions.(*36, 37*) We next investigated biomarkers associated with established uptake, transport, and adhesion gene sets (Figure 2C). (*38–40*) To examine the distribution of biomarker significance across curated gene sets and NP formulations, each gene was visualized using the -log(q-value) for gene expression. As expected, we identified highly significant biomarkers from gene sets important in drug import and export such as solute carrier (SLC) transporter family and ATP-binding cassette (ABC) family. Importantly, our screen provides data on both the significance and the relationship to NP delivery. For example, we found that ABCA1, which plays a role in cholesterol transport, has a positive relationship with liposomal NPs, while several members of the multidrug resistance subfamily (ABCB1/P-GP, ABCC1/MRP, ABCC4/MRP4) have a negative relationship with PLGA NPs (Figure S11).(*41*) We also identified biomarkers important for cell engagement (focal adhesion, extracellular matrix) as well as intracellular trafficking (vesicular transport, lysosome, and cholesterol transport). The significance of biomarkers in these curated gene sets were similarly between NP formulations and targeted antibodies. This highlights the ability of our screen to identify expected biomarkers and enable comparison between drug delivery modalities.

We also observed that liposome surface modification influences the number and significance of biomarkers. Specifically, liposomes electrostatically coated with polysaccharides (HA, ALG, DXS, FUC, CS) had the highest amount of associated biomarkers, which we hypothesize is due to the high degree of interactions between sugars and cell surface proteins as well as the potential for naturally occurring polysaccharides to interact with a wide range of cell surface elements.(*23, 42, 43*) In line with this hypothesis, the addition of PEG, a well-established antifouling polymer, reduces the number and significance of associated biomarkers almost to zero. In contrast to the highly specific hits generated from EGFR-conjugated liposomes (formulated using 25% PEG liposomes), this abrupt decrease in significant biomarkers further indicates the ability of our platform to identify specific NP binding and recognition elements. In contrast to the liposomal formulations, PLGA formulations, regardless of surface modification, resulted in few biomarkers at either time point. Lastly, a high number of significant biomarkers was associated with both carboxylated and sulfated PS NPs included in our screen, though there was no time dependence, in contrast to the liposomal formulation. While this result was initially surprising, as the PS formulations are made of synthetic polystyrene polymers, meaningful biological interactions with anionic polystyrenes, both in polymer and particle form, have been reported.(*44*) Specifically, it was described that nanoparticles bearing anionic polystyrene motifs have the appropriate mix of hydrophobicity and anionic charge character to interact favorably with trafficking proteins, including the caveolins.

#### NP biomarkers are connected and create trafficking networks

We then used an unbiased approach to identify predictive biomarkers using a random-forest algorithm, annotated by feature set: gene expression, gene copy number, and protein abundance (methods in Supplementary Text). Data from the 4 h time point was chosen for this analysis based on the EGFR-related hits for liposomes, which were more significant at 4 h than at 24 h. As we were interested in applying this approach to identify cellular features positively correlated with uptake (e.g., increased expression of trafficking proteins), hits negatively correlated with NP association were removed from this analysis. Next, we used K-means clustering to visualize biomarkers based on their relative importance and presence across formulations (Figure 2D). Clusters 1 and 2 contained 205 hits shared across NP formulations and were especially enriched for liposomal and PS NPs. These genes and proteins were input into the STRING database(*45–47*) to generate a protein-protein interaction (PPI) network that was found to be highly interconnected (PPI enrichment p-value <1×10^-16^) (Figure 2E). Notably, the network is enriched in proteins found in the plasma membrane, extracellular region, and extracellular matrix (false discovery rate [FDR] = 8×10^-12^, 3×10^-9^, and 3×10^-8^, respectively) based on enrichment analysis with gene ontology (GO) localization datasets (Figure S12).(*48–50*) The identification of overlapping biomarkers that are localized to the cell surface and have established protein-protein interactions led us to hypothesize that these proteins are important in early NP trafficking. Enrichment analyses using GO molecular functions datasets showed enrichment in numerous binding processes (Data S1, Figure S12), giving further credence to this theory. Our results serve as a framework for the comprehensive investigation of cellular processes important for NP engagement, which may prove useful for fundamental trafficking studies and target identification.

### SLC46A3 is a negative regulator of liposomal NP uptake

Evaluating univariate results across NP formulations, we identified one biomarker with a strong, inverse relationship with liposomal NP association: expression of solute carrier family 46 member 3 (*SLC46A3*). A member of the solute carrier (SLC) transporter family, SLC46A3, is a relatively unstudied transporter that has been localized to the lysosome.(*51, 52*) SLC46A3 was recently identified as a modulator of cytosolic copper homeostasis in hepatocytes, connecting hepatic copper levels with lipid catabolism and mitochondrial function.(*53*) This reported relationship between SLC46A3 and lipid catabolism may help to explain why SLC46A3 found to have a strong relationship with liposomal NP uptake and not uptake of polymeric NPs. In the context of cancer, SLC46A3 was recently shown to transport non-cleavable antibody-drug conjugate (ADC) catabolites from the lysosome to the cytosol, thereby being necessary for therapeutic efficacy. In this context, downregulation of SLC46A3 was identified as a resistance mechanism for antibody-drug conjugate delivery in cancer cells, including in patient samples of multiple myeloma.(*54–57*) While the biologic function of SLC46A3 in cancer is not yet clear, given the potential therapeutic implications and the unique, inverse relationship between SLC46A3 expression and NP delivery, we sought to validate the predictive power of *SLC46A3* as a biomarker for liposomal NP association.

*SLC46A3* expression was the most significant hit on univariate analysis and also the top ranked random forest feature for each liposomal NP tested at 24 h, regardless of surface modification (Figures 3A and S13). This inverse relationship between *SLC46A3* expression and NP association was found to be specific to liposomal NPs, and not observed with PLGA or PS NPs, and was maintained regardless of cancer cell lineage (Figures 3B and S8).

**Figure 3.**
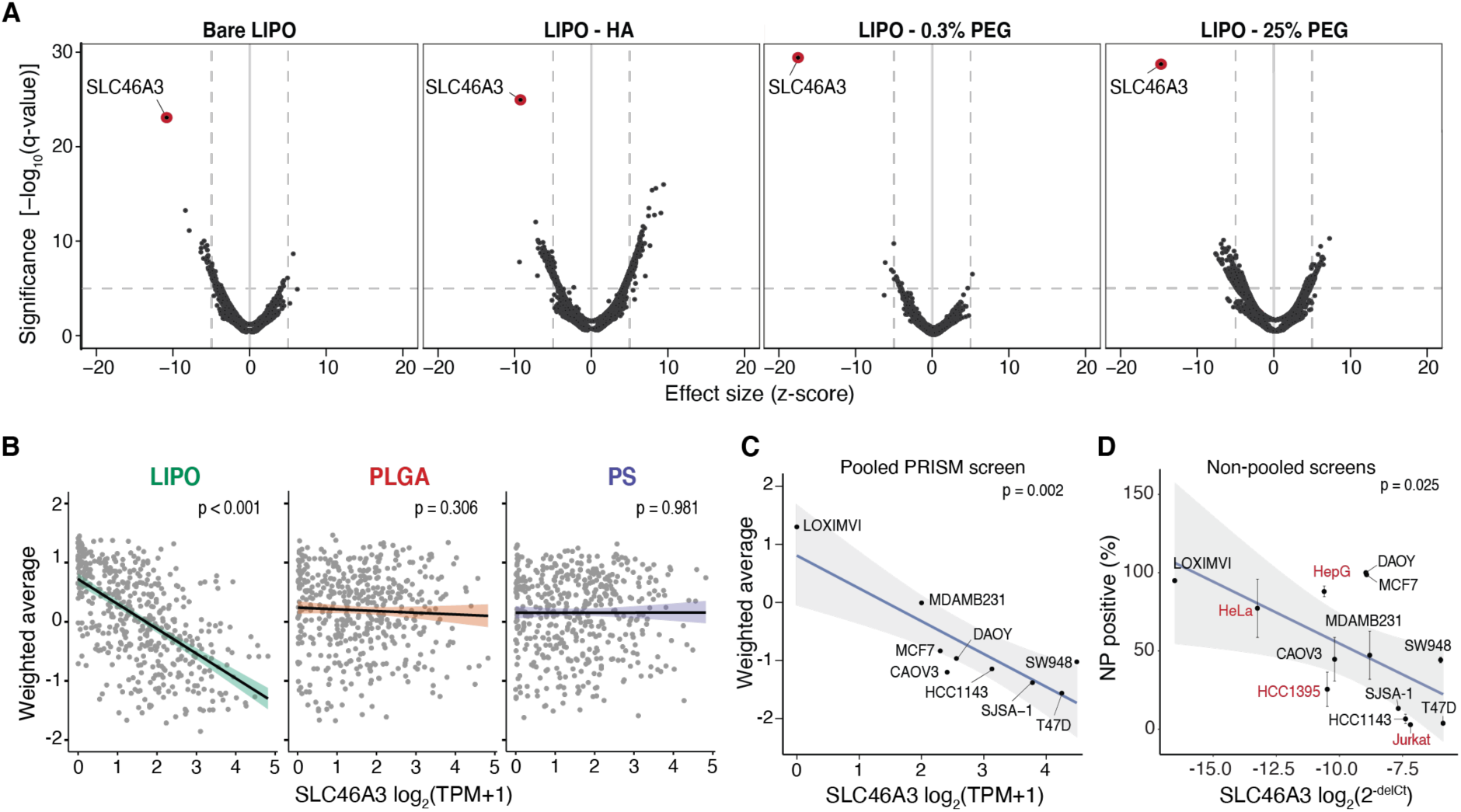
Native expression of the lysosomal transporter SLC46A3 is strongly predictive of NP-cell interaction for liposome formulations. (**A**) Univariate analysis identifies SLC46A3 expression as strongly yet inversely correlated with liposome association, regardless of liposomal surface modification. (**B**) Using linear regression to evaluate the biomarker relationship across core formulations reveals *SLC46A3* expression is inversely correlated with NP association in liposome-cell line pairs (*p* < 0.001) but not PLGA- and PS-cell line pairs (*p* > 0.05); n=488 for each plot. (**C**) Cell lines in the nanoPRISM pool exhibit a range of natural *SLC46A3* expression levels with a log linear correlation with uptake of liposomes. (**D**) This correlation is also exhibited when assessing liposome-cell associations via flow cytometry in a non-pooled fashion (*p* = 0.025). Data for Bare-Lipo is shown here. Cell lines in red were not part of the pooled PRISM screen. Data represented in D is shown as the mean and standard deviation of four biological replicates. Error bars are not shown when smaller than data points.

We selected nine cancer cell lines from the nanoPRISM pool and four additional cell lines, spanning multiple lineages, with a range of native *SLC46A3* expression levels for screening in a non-pooled fashion (Figures 3C-D, S3, S14-S15). Analogous to the pooled screen, individual cell lines were profiled using flow cytometry and NP-associated fluorescence was quantified after 24 h incubation; here SLC46A3 expression was concurrently quantified using quantitative polymerase chain reaction (qPCR) (Figures 3D and S9). In line with observations from pooled screening, the inverse relationship between liposome association and native *SLC46A3* expression was maintained, suggesting that *SLC46A3* may play a key role in regulating the degree of liposomal NP uptake.

To probe whether SLC46A3 expression level governs NP association, we selected the breast cancer cell line T47D, which exhibited low association with liposomal NP formulations and high SLC46A3 expression (Figure 4A). We knocked down SLC46A3 through the use of siRNA and evaluated the effect on liposomal NP association. We observed that T47D cells with reduced SLC46A3 levels had higher NP-cell association with both tested formulations, suggesting that modulating SLC46A3 expression alone can regulate NP-cell association levels (Figure 4B).

**Figure 4.**
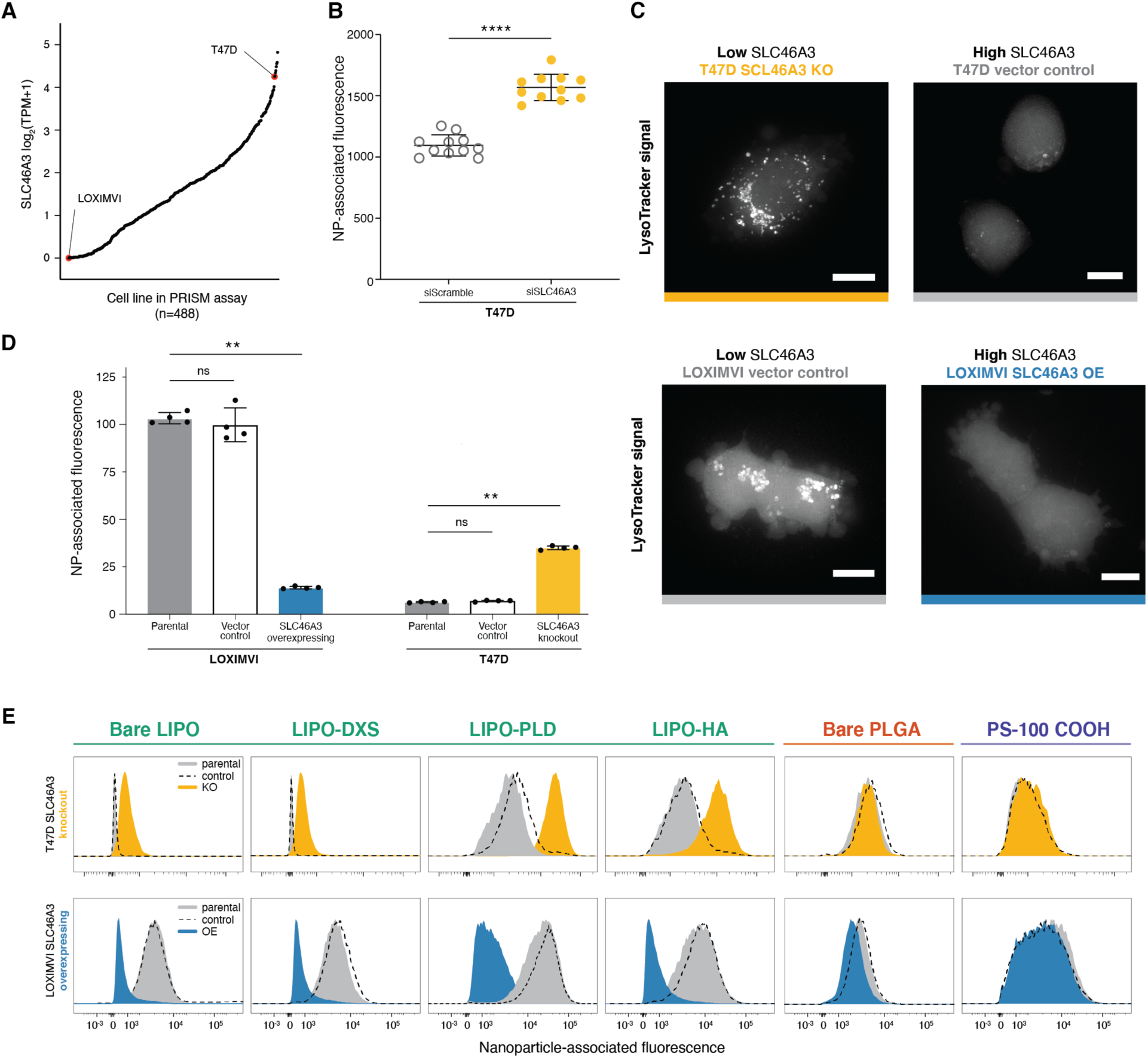
Modulating *SLC46A3* expression in cancer cell lines is sufficient to negatively regulate interaction with liposome NP formulations. (**A**) T47D and LOXIMVI cells have high and low *SLC46A3* expression, respectively, with respect to *SLC46A3* expression levels represented in the nanoPRISM cell line pool. (**B**) T47D cells treated with siRNA to knock down SLC46A3 have higher association with Lipo-PLD compared to T47D cells treated with a scrambled siRNA control (**** *p* < 0.0001, Mann-Whitney test). (**C**) Representative micrographs of Lysotracker signal in engineered cell lines showing endolysosomal compartments. Scale bars = 10 μm. (**D**) Using lentivirus to overexpress *SLC46A3* in LOXIMVI cells and CRISPR/Cas9 to knock out *SLC46A3* in T47D cells, we show that modulation results in significantly changed liposome association, as determined via flow cytometry (** *p* < 0.001, Kruskal-Wallis test), NP-associated fluorescence is defined as median fluorescence intensity normalized to untreated cells. Data is represented as the mean and standard deviation of four biological replicates. (**E**) Shifts in NP association were consistently observed across all tested liposomes, independent of surface modification. No shifts were observed with PLGA or PS formulations.

To further functionally evaluate the relationship of SLC46A3 expression and NP-cell association, we selected two cancer cell lines from the pooled screen that displayed strong phenotypes (Figure 4A): the T47D cell line and the melanoma cell line LOXIMVI, which exhibited high association with liposomal NP formulations. We developed a toolkit using these two cell lines by knocking out *SLC46A3* in T47D cells and inducing overexpression in LOXIMVIs (Figures S16A-G).

As SLC46A3 is a protein associated with lysosomal membranes(*54, 55, 58*), we utilized LysoTracker dye to evaluate the effect of SLC46A3 modulation on endolysosomal compartments in both T47D and LOXIMVI engineered cell lines (Figure 4C). We observed an SLC46A3-dependent change: cells with lower SLC46A3 expression (T47D-vector control, LOXIMVI-SLC46A3 OE) exhibited more brightly dyed endolysosomal compartments compared to their high SLC46A3 expression counterparts (T47D-*SLC46A3* knockout, LOXIMVI-vector control).

Overexpression of SLC46A3 in LOXIMVI cells significantly abrogated interaction with bare liposomes (*p* = 0.006) using flow cytometry profiling (Figure 4D). The T47D-*SLC46A3* knockout cell line demonstrated significantly increased association with bare liposomes compared to parental or vector control lines (*p =* 0.0017, Figure 4D). We further confirmed that these trends are generalizable across a range of surface functionalized liposomes (Figure 4E, S16H). Moreover, no significant changes in NP association were observed for PLGA and PS NPs (Figures 4E, S16I-J). We also confirmed that the presence of serum proteins in cell culture media does not abrogate this trend (Figure S16K). Taken together, these data indicate modulation of SLC46A3 alone in cancer cells is sufficient to negatively regulate association and uptake of liposomal NPs.

As flow cytometry does not provide spatial information with respect to NP-cell interactions, we employed imaging cytometry to characterize NP localization in a high throughput manner (Figure 5A-F). We selected four representative formulations: three liposomal NPs to probe the relationship of SLC46A3 expression with liposome trafficking; and one PLGA NP formulation with a common outer layer.

**Figure 5.**
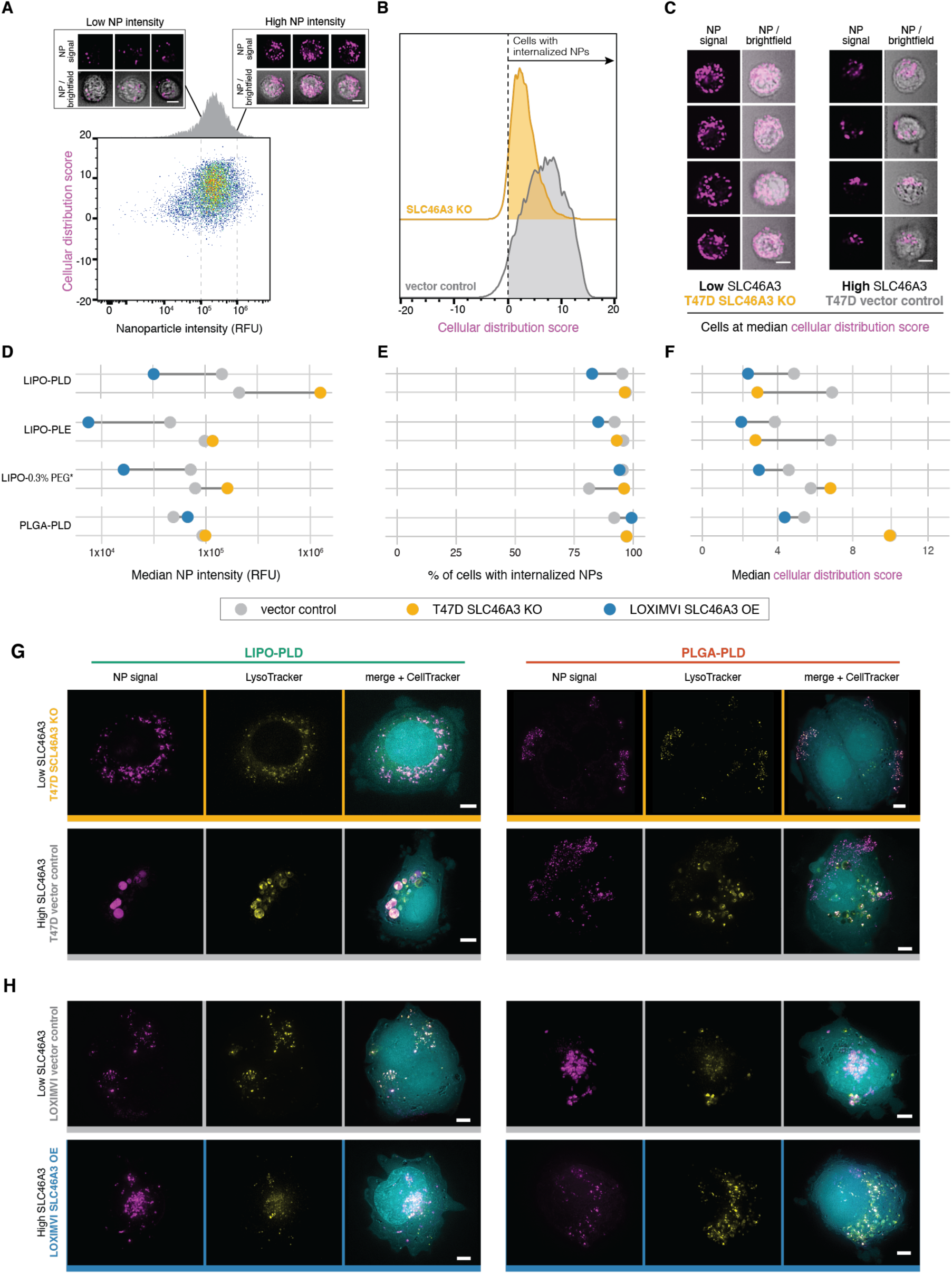
High throughput imaging cytometry confirms NP internalization and reveals SLC46A3 dependent changes to intracellular trafficking. (**A**) Imaging cytometry was used to investigate the intensity (x-axis) and distribution (y-axis) of NPs in a high-throughput manner. Bivariate density plot of n=10,000 cells (T47D-vector control) after 24 h incubation with LIPO-PLD NPs, with representative cell images at low and high NP signal. (**B**) Cellular distribution patterns of NPs were scored such that scores greater than 0 indicate cells with internalized NPs. Representative data from LIPO-PLD NPs in engineered T47D cells are shown. (**C**) Representative cell images at the median cellular distribution score for engineered T47D cells treated with LIPO-PLD NPs. (**D**) Quantification of median intensity of tested NP formulations in engineered T47D and LOXIMVI cell lines demonstrated SLC46A3-dependent changes. (**E**) NPs remained predominantly internalized independent of SLC46A3 expression levels. (**F**) Shifts in the median cellular distribution scores were observed in response to SLC46A3 modulation. Live cell micrographs of (**G**) T47D-vector control and T47D-*SLC46A3* knockout cells and (**H**) LOXIMVI-vector control and LOXIMVI-SLC46A3 OE cells incubated with LIPO-PLD and PLGA-PLD NPs for 24h. NP signal is pseudo-colored magenta, LysoTracker signal yellow, and CellTracker cyan. Scale bar = 5 μm.

Consistent with trends observed by flow cytometry, we observed an inverse relationship between NP intensity and SLC46A3 expression for liposomal, but not PLGA, NPs (Fig 5A, D, S11). Using brightfield images, we applied a mask to investigate cellular localization of NPs. All tested formulations were internalized, and this did not change with SLC46A3 modulation (Figure 5B, E).

We investigated localization of NPs by scoring NP signal based on distribution within each cell (Figure 5C, F, Figure S17D). We observed stark differences in median cellular distribution scores of liposomal NPs in relation to SLC46A3 expression levels in T47D cells. This was not observed for PLGA NPs, mimicking the previously observed core-specific relationship between NP-cell association and SLC46A3 expression. Changes in this score, though less pronounced, were also observed for liposomal NPs in LOXIMVI cells.

To confirm our findings with higher spatial resolution, we employed deconvolution microscopy of live cells and incorporated a lysosomal stain to observe changes in intracellular trafficking (Figure 5G-H). NPs appeared uniformly distributed within T47D-*SLC46A3* KO cells, co-localizing with endolysosomal vesicles. In contrast, LIPO-PLD NPs were localized to large endolysosomal clusters in T47D-vector control cells. This trend was also observed for LIPO-PLE and LIPO-0.3% PEG* NPs and at the earlier time point of 4 h (Figure S18). Changes in localization were not observed for the tested PLGA PLD NPs. This again indicates a NP core-dependent relationship with SLC46A3.

In the engineered LOXIMVI cell lines, we also observed co-localization of liposomal NPs with endolysomal signal. However, predictable changes in NP localization were not detected, in line with smaller changes in median cellular distribution scores.

#### Impact of SLC46A3 expression on endolysosomal maturation is minimal

To further probe the relationship between intracellular liposomal NP trafficking and SLC46A3 expression, we utilized imaging cytometry to spatially interrogate markers of endolysosomal transport. We elected to study markers of early (EEA1, Rab5A), late (Rab7), and recycling endosomes (Rab11) as well as lysosomes (LAMP1) in engineered LOXIMVI cells (Figures S19-S20). While no apparent differences in endolysosomal marker signal strength, size, and shape were observed when comparing LOXIMVI-SLC46A3 OE and LOXIMVI-vector control cells both in the absence and presence of liposomal NPs, modest changes in EEA1, Rab7, and LAMP1 texture were noted (Figure S19A-B). The significance of these morphologic changes is not clear, but our data supports a model in which SLC46A3 does not directly impact the number or localization of endosomes or lysosomes.

We then assigned colocalization values between each endolysosomal marker and NP signals. (Figure S19C-F). For both tested liposomal NP formulations we observed increasing colocalization from EEA1, Rab5, and Rab7, consistent with liposome trafficking from early to late endosomes. Colocalization between Rab7 and liposomal NPs was higher in LOXIMVI-SLC46A3 OE cells compared to vector control and the opposite relationship was observed for LAMP1 colocalization. Taken together, this may suggest that the effect of SLC46A3 expression on NP trafficking may be localized to late endosomes and lysosome compartments, potentially leading to retention of NPs in late endosomes or increased removal from lysosomes when SLC46A3 expression is elevated. Given prior reports of SLC46A3 localization in the lysosome,(*52*) pinpointing NP trafficking changes around endolysosomal vesicles is not surprising.

#### Liposome retention and accumulation remains SLC46A3-dependent in vivo

To evaluate the potential clinical utility of SLC46A3 as a negative regulator of liposomal NP delivery, we tested *in vivo* delivery of an FDA-approved nanoparticle analog, the drug-free version of liposomal irinotecan (LIPO-0.3% PEG*), in mice bearing subcutaneous LOXIMVI flank tumors. Fluorescently-labeled NPs were administered via a one-time intratumoral (IT) injection or repeat intravenous (IV) administration to evaluate tumor retention and accumulation, respectively (Figure 6A, Figure S21).

**Figure 6.**
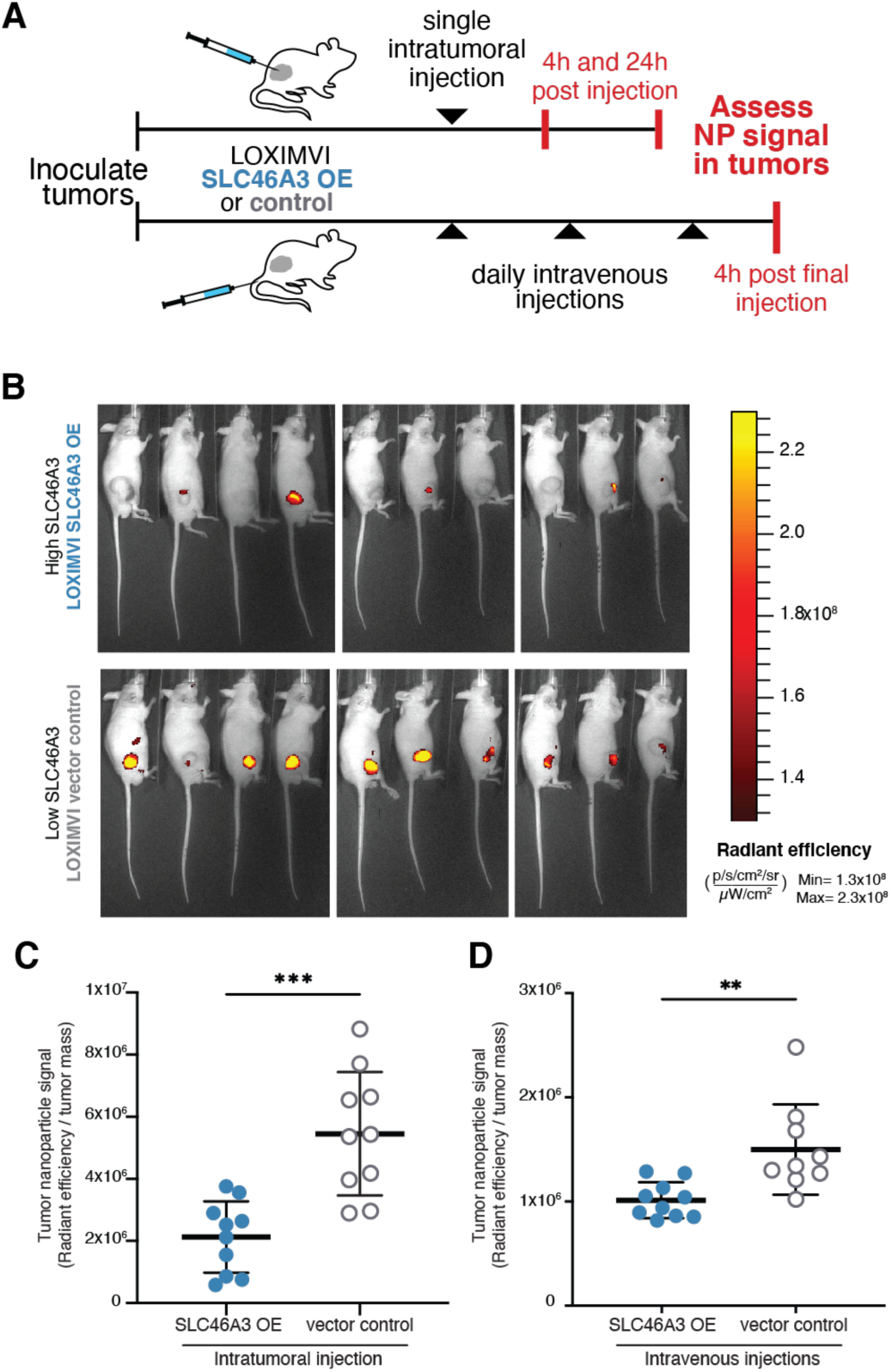
Retention and accumulation of PEGylated liposomes (LIPO-0.3% PEG*) in LOXIMVI tumors is dependent on SLC46A3 expression. (**A**) Fluorescently labeled LIPO-0.3% PEG* NPs were administered to mice bearing LOXIMVI flank tumors via a one-time intratumoral injection or repeat intravenous injections. (**B**) Whole animal fluorescence images of mice (4 males, 6 females per group) 24 h after being intratumorally injected with LIPO-0.3% PEG* NPs. (**C**) Quantification of LIPO-0.3% PEG* NP retention 24 h after intratumoral administration to LOXIMVI flank tumors. (**D**) Quantification of LIPO-0.3% PEG* NP accumulation after repeat IV injections. In panels C-D, nanoparticle signal is expressed on the y-axis as radiant efficiency divided by tumor mass (mg). The mean and standard deviation of n = 10 are shown with the exception of the LOXIMVI-vector control, repeat IV injection group, where n = 9 (** < 0.01, *** < 0.001, Mann-Whitney test).

NP signal was quantified both 4 and 24h following IT administration. In line with our hypothesis, as well as *in vitro* NP-associated fluorescence data (Figure S21A), we observed an inverse relationship between SLC46A3 expression and LIPO-0.3% PEG* NP retention that became more pronounced over time (*p* = 0.0115, 4 h; *p* = 0.0002, 24 h) (Figure 6C-D, Figure S21B-E). Moreover, these findings also align with our initial nanoPRISM findings, in which SLC46A3 expression was a more significant biomarker at 24 h (q-value=3.49×10^-30^, Data S2, Figure S13A) than at 4 h (q-value=1.47×10^-4^, Data S2, Figure S13A).

To determine if this newly identified biomarker can be used to predictably govern accumulation of nontargeted NPs, which bear no specific functional ligands on their surface, following systemic administration, we quantified NP signal following IV injections. Notably, we observed a significant relationship between SLC46A3 and NP accumulation (*p* = 0.0019) (Figure 6D, Figure S21F). This demonstrates the predictive power of SLC46A3 as a NP specific biomarker that holds true even in complex physiologic settings.

Together, these data highlight the real-world relevance of the nanoPRISM screening assay in general as well as the utility of SLC46A3 in particular a clinically actionable biomarker.

#### Solid lipid nanoparticle uptake and transfection are dependent on SLC46A3 levels

Given the recent translational success and promising potential of nucleic acid-carrying solid lipid nanoparticles (LNPs),(*59, 60*) we sought to determine if the relationship of SLC46A3 expression extends to LNP association as well as transfection efficiency. We generated fluorescently (Cy5) labeled LNPs containing messenger RNA (mRNA) encoding green fluorescent protein (GFP) (LNP 1) and incubated these particles with engineered LOXIMVI cell lines (Table S3, S5).

LNP association, as quantified by Cy5 signal, was significantly lower for LOXIMVI-SLC46A3 OE cells than LOXIMVI-vector control cells, showing the same relationship (lower SLC46A3 expression correlating with higher association) for LNPs that was shown for liposomal NPs (Figure 7A-B). Importantly, the same trend was seen for transfection, as quantified by GFP signal of formulation LNP 1 (Figure 7C). Taken together, these findings suggest that SLC46A3 regulates cytosolic delivery of mRNA cargo by way of LNP uptake. Expanding on this, we generated two additional LNPs, analogous to commercial formulations (Table S5).(*61–64*) The inverse trend between SLC46A3 expression and transfection was seen for all LNP formulations tested (Figure 7C). Confirmation of the inverse relationship between SLC46A3 expression and cell association in multiple LNP formulations validates the broad relevance of SLC46A3 as a predictive biomarker for lipid-based nanoparticle formulations.

**Figure 7.**
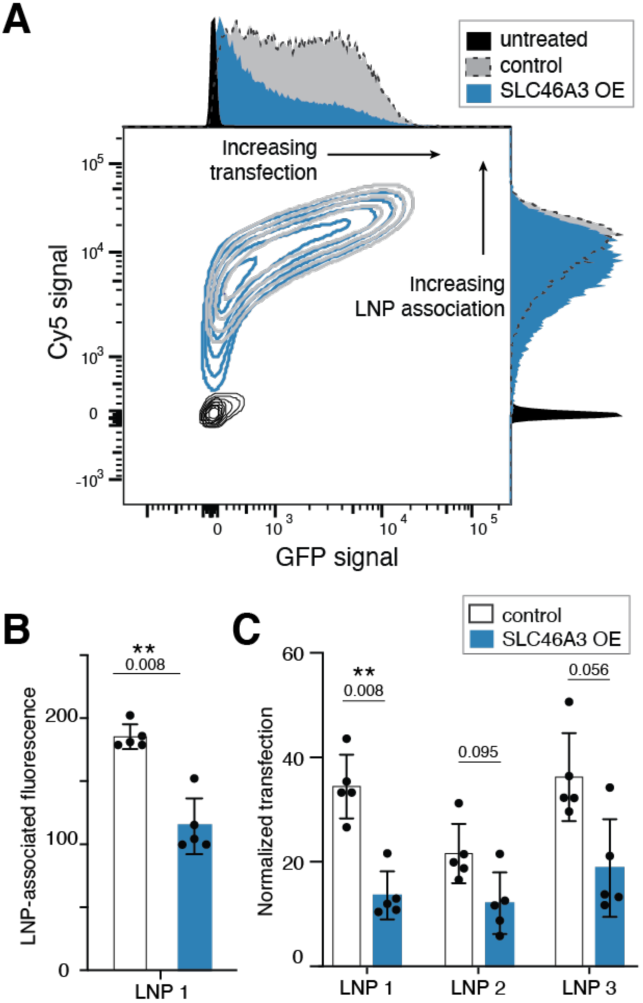
Solid lipid nanoparticle-cell association and transfection are *SLC46A3* dependent, as determined via flow cytometry. (A) Contour plot of Cy5 signal and GFP signal indicating decreased LNP-cell association and transfection efficacy in LOXIMVI cells overexpressing *SLC46A3.* **(B)** Quantification of LNP signal reveals a significant change in LNP-cell association across control and *SLC46A3* overexpressing LOXIMVI cells (** *p =* 0.008, Mann-Whitney). LNP-associated fluorescence is defined as median fluorescence intensity normalized to untreated cells. (**C**) Quantification of GFP signal reveals an *SLC46A3-*dependent trend in transfection efficacy for three tested LNP formulations (Mann Whitney). Normalized transfection is defined as median GFP intensity normalized to untreated cells.

## Discussion and Conclusions

This work represents the first high-throughput interrogation of NP-cancer cell interactions through the lens of multi-omics. Harnessing the power of pooled screening and high throughput sequencing, we developed and validated a platform to identify predictive biomarkers for NP interactions with cancer cells. We utilized this platform to screen a 35 member NP library against a panel of 488 cancer cell lines. This enabled the comprehensive study and identification of key parameters mediating NP-cell interactions, highlighting the importance of considering both nanomaterials and cellular features in concert.

While pooled screening is a powerful tool, we also note several important limitations. First, we primarily focused on lipid-based and polymeric NP formulations with translational drug delivery potential. We recognize that there are several additional categories of nanomaterials with wide ranging properties, such as inorganic systems, that can be useful for both therapeutic and diagnostic applications(*65, 66*) and believe additional biomarkers mediating the trafficking of inorganic NPs may be identified using similar screening approaches. Second, the results of *in vitro* screens are often met with limited success when translated *in vivo,* as NP-mediated delivery is dependent on many factors beyond the nano-cell interface.(*8*) However, the level of molecular characterization and statistical/computational power afforded by annotated biological datasets, such as the Cancer Cell Line Encyclopedia, is currently unrivaled. Therefore, existing *in vivo* screens cannot yet provide this breadth or statistical power. Keeping translational barriers in mind is key to successful validation of candidate biomarkers, and for this reason we employed multiple isogenic models and tested a range of lipid-based nanoparticles across *in vitro* and *in vivo* conditions. Third, an additional limitation of this screen is related to the availability of genomic datasets for each cell line tested, as dataset completeness contributes to the power of detection for both univariate and multivariate analyses. At the time of analysis, ten feature sets were available for the majority of cell lines in our pool (Figure S22). However, as datasets expand over time, it will be possible to re-analyze our data in the future. Especially for emerging fields such as proteomics and metabolomics, the opportunity to intersect nanoparticle delivery metrics with additional datasets could add a new dimension to our existing findings.

One strength of our screening approach is the use of robust analytical tools, such as univariate analyses and random forest algorithms, enabling us to identify biomarkers correlated with NP association. The robust and quantitative nature with which we detected EGFR hits for antibodies as well as antibody-targeted NPs shows the utility of this platform for the development and optimization of targeted drug delivery platforms, including antibody-targeted NPs and with potential to apply to other targeted therapeutics, including ADCs.

By clustering NP-specific biomarkers across formulations, we constructed interaction networks, identifying and connecting genes associated with NP binding, recognition, and subcellular trafficking. This provides the scientific community with a blueprint for the fundamental study of cellular processes mediating NP engagement, with applications for both basic and translational research.

We identified expression of SLC46A3, a lysosomal transporter, to be a negative regulator and predictive biomarker for lipid-based nanoparticle uptake and downstream functional efficacy. While SLC46A3 has recently been implicated in hepatic copper homeostasis as well as sensitivity to ADCs in cancer cells,(*53–55*) its role in NP delivery was previously unexplored. We first validated SLC46A3 as a negative regulator of lipid-based NP uptake in a panel of non-pooled cell lines, as well as engineered isogenic cell lines with modulated *SLC46A3* levels. Importantly, as all current FDA approved NPs for anticancer applications are liposomal formulations, there is significant potential for this biomarker to be quickly implemented in clinical studies with existing, approved formulations. To this end, we recapitulated our findings in an *in vivo* model using an analog of an FDA-approved liposomal NP formulation.

Moreover, we demonstrated that SLC46A3 has potential as a predictive biomarker beyond liposomal nanoparticles by investigating solid lipid nanoparticles. Both LNP cell-association *and mRNA transfection* were inversely correlated with SLC46A3 levels. These preliminary findings demonstrate that SLC46A3 expression may serve as a predictive biomarker for functional delivery of nucleic acid cargo via lipid nanoparticles. Our findings strongly support the continued exploration of SLC46A3 as a potential clinical biomarker for therapeutic nanoparticle delivery.

In summary, we present a powerful platform to study NP-cancer cell interactions simultaneously through the use of pooled screening, genomics, and machine learning algorithms. This provides a new dimension to the study of cancer nanomedicine. Application of this platform will serve useful not only for the rational design of nanocarriers, but also for the identification of specific phenotypes primed to benefit from targeted drug delivery and nanomedicine.

## Acknowledgments

This work was supported in part by SPARC funding at The Broad Institute.

This work was also supported by a grant from the Koch Institute’s Marble Center for Cancer Nanomedicine.

This work was supported in part by the Koch Institute Support (core) Grant P30-CA14051 from the National Cancer Institute.

We thank the Koch Institute’s Robert A. Swanson (1969) Biotechnology Center for technical support, specifically the Flow Cytometry, High Throughput Sciences, Genomics Core, Microscopy, and Preclinical Modeling, Imaging & Testing cores, the Hope Babette Tang (1983) Histology Facility, and the Peterson (1957) Nanotechnology Materials Core Facility.

NB was supported by a Department of Defense Congressionally Directed Medical Research Programs Peer Reviewed Cancer Research Program Horizon Award (W81XWH-19-1-0257) and the NIH-NCI (K99CA255844).

JPS was supported as a National Institutes of Health grant T32 trainee (CA136432-08) and by the Helen Gurley Brown Presidential Initiative of Dana-Farber Cancer Institute.

Fellowship support for CHA was from the NIH (NRSA F31 CA228241-01). RRC is a fellow of the Parker B. Francis Foundation.

NN was supported by a grant from the Gates Foundation.

Fellowship support for AGB was from the NIH (F30 DK130564) and a Termeer Fellowship of Medical Engineering and Science.

NGL was supported by Cancer Research UK and the Brain Tumour Charity (C42454/A28596) and a fellowship from the Ludwig Center at the Koch Institute for Integrative Cancer Research.

We would like to thank Todd Golub and Alex Burgin for formative feedback and helpful discussion.

We also gratefully acknowledge Thomas Diefenbach and the Ragon Institute Microscopy Core for assistance with imaging cytometry.

Figures 1A and 6A were created in part using Biorender.com

## Author contributions

Conceptualization: NB, JPS

Methodology: NB, JPS, MK

Formal Analysis: NB, JPS, MK, MGR, MR

Investigation: NB, JPS, HCS, MGR, DR, NN, AGB, NGL

Visualization: NB, JPS

Funding acquisition: NB, JPS, ANK, PTH

Project administration: NB, JPS, MR

Validation: NB, JPS, HCS, CHA, RRC, JHC, HL

Supervision: JAR, ANK, PTH

Writing – original draft: NB, JPS

Writing – review & editing: NB, JPS, HCS, MK, MGR, MR, CHA, RRC, NN, AGB, NGL, JHC, HL, JAR, ANK, PTH

## Data and materials availability

All data are available in the main text or the supplementary materials.

## Materials and Methods

### Materials

#### Reagents

1,2-distearoyl-sn-glycero-3-phospho-(1’-rac-glycerol) (sodium salt) (DSPG), 1,2-distearoyl-sn-glycero-3-phosphoethanolamine (DSPE), 1,2-distearoyl-sn-glycero-3-phosphocholine (DSPC), 1,2-dioleoyl-sn-glycero-3-ethylphosphocholine (chloride salt) (18:1 EPC), L-α-phosphatidylcholine (Soy-PC), 1,2-distearoyl-sn-glycero-3-phosphoethanolamine-N- [carboxy(polyethylene glycol)-2000] (sodium salt) (MPEG-2k-DSPE, for Ab liposomes), 1,2-distearoyl-sn-glycero-3-phosphoethanolamine-N-[methoxy (polyethylene glycol)-2000] (ammonium salt) (PEG_PE), and cholesterol were purchased from Avanti Polar Lipids. Sulfo-cyanine5 NHS ester and cyanine5 free acid were purchased from Lumiprobe. Methoxy PEG amine (HCl salt), MW 2000 Da (for PS PEGylation) was purchased from JenKem Technologies. Chloroform and methanol were purchased from TCI and Sigma, respectively. Poly(D,L-lactide-glycolide) (Resomer RG502H, 7 kDa:17 kDa) and Poly(D,L-lactide-glycolide) 50:50-b-PEG (10kDa PLGA, 2 kDa PEG) were purchased from Sigma.

Non-glycosylated monoclonal human IgG1 antibody against human EGFR (hegfr-mab12, lot: EG12-39-01) and isotype control -Human IgG1 (bgal-mab1, lot: BG1-41-01) were purchased from InvivoGen. Human EGFR (Research Grade Cetuximab Biosimilar) Alexa Fluor® 488-conjugated Antibody was purchased from R&D Biosystems. Poly-L-arginine hydrochloride (PLR200, 38.5 kDa), poly-L-aspartic acid sodium salt (PLD100, 14 kDa), and poly-L-glutamic acid sodium salt (PLE100, 15 kDa) were purchased from Alamanda Polymers. Sodium hyaluronate (40 kDa) was purchased from Lifecore Biomedical. Dextran sulfate (15 kDa), fucoidan (from fucus vesiculosus), and polyacrylic acid (8 kDa) were purchased from Sigma. Sodium alginate was purchased from NovaMatrix. Chondroitin sulfate A (10-30 kDa average MW) was purchased from Carbosynth Ltd. Yellow-green fluorescent polysytrene microspheres (Fluospheres), 5 M bioreagent grade NaCl solution, and 1 M bioreagent-grade HEPES were purchased from Fisher Scientific.

Whatman nuclepore polycarbonate hydrophilic membranes (400, 200, 100 and 50 nm sizes) were purchased from GE. All glassware was obtained from Chemglass. 5 mL Falcon brand round-bottom tubes with 35 µm cell strainer cap, 50/15 mL Falcon tubes and 50/5/2 mL DNA/Protein loBind Eppendorf tubes were purchased from VWR. D02-E100-05-N and C02-E100-05-N tangential flow filtration filters were purchased from Repligen. Polystyrene semi-micro cuvettes for the Malvern Zetasizer were purchased from VWR and DTS1070 folded capillary cells were purchased directly from Malvern. Black, glass bottom 364 well plates for the Wyatt DLS were purchased directly from the Peterson (1957) Nanotechnology Core Facility.

RPMI-1640 (Invitrogen/Corning), and PBS solution pH 7.4 (Gibco) were purchased from Fisher Scientific. Heat inactivated fetal bovine serum (FBS), Tween20, Igepal CA-630, Trizma hydrochloride, and Potassium chloride solution (BioUltra, ∼1 M in H2O) were purchased from Sigma. Proteinase K was purchased from Qiagen. Matrix Deepwell Storage Blocks (96 well, 1 mL) were purchased from ThermoFisher. Tissue culture plasticware, trypsin EDTA, Accutase, and penicillin streptomycin were purchased from Corning. LabTek 8-chamber coverslips (cat. no.155409), CellTracker Blue CMAC, CellTracker Orange CMRA, and LysoTracker Green were purchased from Fisher Scientific. 16% formaldehyde (methanol free) was purchased from Invitrogen. Normal goat serum was purchased from Cell Signaling Technologies. Bovine serum albumin and saponin were purchased from Sigma. Hoechst 33342 (10 mg/mL) was purchased from ThermoFisher.

Opti-mem media and Lipofectamine RNAiMAX Transfection reagent were purchased from ThermoFisher. siRNA (Silencer Select pre-designed siRNA) to silence SLC46A3 (siSLC46A3, s49280) and negative control no. 1 (siScramble) were purchased from Life Technologies.

Dilinoleylmethyl-4-dimethylaminobutyrate (DLin-MC3-DMA) was purchased from MedChemExpress. 1,2-dimyristoyl-*rac*-glycero-3-methoxypolyethylene glycol-2000 (DMG-PEG-2000) was purchased from Cayman Chemical Company. Cholesterol, 1,2-distearoyl-sn-glycero-3-phosphocholine (DSPC), 1,2-dioleyol-sn-glycero-3-phosphoethanolamine (18:1 (Δ9-Cis) PE) and 1,2-dioleyol-sn-glycero-3-phosphoethanolamine-N-(Cyanine-5) (18:1 Cy5 PE) were purchased from Avanti Polar Lipids. Absolute ethanol (200 proof), molecular-biology grade and UltraPure DNAse/RNAse-free Distilled Water were purchased from Fisher Scientific. VWR Spinbar Micro Stir Bars and 4-ml Amber Borosilicate Glass sample vials were purchased from VWR.

CleanCap Enhanced Green Fluorescent Protein mRNA (5-methoxyuridine) was purchased from TriLink Biotechnologies. The Quant-it™ RiboGreen RNA Assay Kit, Nunc F96 MicroWell Black polystyrene plates, and 3 M sodium acetate solution, pH 5.2, RNAse free, were purchased from Fisher Scientific. Triton X-100 was purchased from Sigma. 1x Tris-EDTA solution, pH 8.0 (IDTE) was purchased from Integrated DNA Technologies.

PCR primers and antibodies used for endolosysomal staining are detailed in their respective methods sections.

#### Cells

The generation and culture conditions of the stably barcoded and pooled PRISM cells (500 human cancer cell lines) are described in reference 15. The CAOV3, DAOY, HeLa, HepG2, HCC1395, HCC1143, MDA-MB-231, MCF7, SJSA-1, and SW948 cell lines were obtained from ATCC. The LOXIMVI and T47D cell lines were gifts from the Gertler Lab, and the Jurkat cell line was a gift from the Sabatini Lab. Cells were cultured at 37 °C with 5% CO_2_. Cell line-specific culture information is provided in the **SLC46A3 Validation Studies** section below.

### Methods

The nanoparticle-specific methods described below detail conditions utilized for synthesis and characterization and have been adapted in part from references 9, 10, 11, and 16.

#### Base Liposome Synthesis

Cholesterol and 1,2-distearoyl-sn-glycero-3-phosphocholine (DSPC) were dissolved in chloroform. 1,2-distearoyl-sn-glycero-3-phosphoethanolamine (DSPE) and 1,2-distearoyl-sn-glycero-3-phospho-(1’-rac-glycerol) (DSPG) were dissolved in a 65:35:8 mixture of chloroform, methanol and deionized water (milli-Q). A lipid mixture composed of 31 mol% DSPC, 31 mol% cholesterol, 31 mol% DSPG and 6 mol% DSPE was prepared in 50 mL round bottom flask and methanol was added dropwise until the solution cleared. The lipid solution was evaporated using a BUCHI rotovap system under heat (60 °C, water bath) until completely dry (<15 mBarr) to make a thin lipid film. A Branson sonicator bath was filled with milliQ water and heated until >65°C. The round bottom flask containing the lipid film was partially submerged in the water bath and milliQ water was added to re-suspend the lipid film to a concentration of 2 mg lipid/mL solution. The liposome solution was sonicated for 1 minute and then removed for 1 minute. This process was repeated three times and then transferred to an Avestin LiposoFast LF-50 liposome extruder. The extruder was connected to a Cole-Palmer Polystat Heated Recirculator Bath to maintain a temperature >65°C throughout the extruder. The liposome solution was extruded through sequentially smaller nucleopore membranes until a 50-100 nm liposome was obtained. This usually required two passes through a stack of one 400 and one 200 nm membrane followed by two passes through one 100 nm membrane and two passes through a 50 nm membrane. These liposomes were fluorescently labeled through NHS-coupling of sulfo-cyanine NHS ester dye to DSPE headgroups according to the dye manufacturer (Lumiprobe) instructions. Lipid film generation, rehydration, extrusion, and dye labeling steps were similarly applied to all liposome formulations unless noted otherwise.

#### Tangential Flow Filtration (TFF)

To remove excess dye, crude nanoparticle solution was connected to a Spectrum Labs KrosFlo II system using masterflex, Teflon-coated tubing. D02-E100-05-N membranes were used to purify the particles until dye was no longer seen in the permeate. This usually required 15- volume equivalent washes to be collected in the permeate. Dye levels in the permeate were monitored by running samples on a Tecan M1000 plate reader. Samples were run at flow rates of 80 mL/min with size 16 tubing. Once purified, the sample was concentrated and then recovered by reversing the direction of the peristaltic pump. To improve nanoparticle yield, 1-3 mLs of water were run backwards through the tubing to recover any remaining particles. 1x PBS was used as the exchange buffer for the first five washes followed by milliQ water for the rest of the purification steps. Following TFF, liposomes were characterized for size and zeta using dynamic light scattering (see **Characterization of Nanoparticles**). For LbL synthesis, TFF was used for purification between after deposition of each polyelectrolyte layer, following the above procedure. Instead of PBS, only milliQ water was passed through the TFF for LbL NP purification.

#### Synthesis of PEGylated Liposomes

To make the lipid stocks, cholesterol, DSPC, DSPE-PEG(2000) carboxylic acid, PEG-PE, and soy PC were dissolved in chloroform while DSPE was dissolved in a 65:35:8 mixture of chloroform, methanol and deionized water (milli-Q). For the 5% PEG formulation (LIPO-5% PEG), a lipid mixture composed of 55.67 mol% DSPC, 33.3 mol% cholesterol, 5 mol% DSPE-PEG(2000) carboxylic acid and 6 mol% DSPE was prepared in a 50 mL round bottom flask. For the 25% PEG formulation (LIPO-25% PEG), a lipid mixture composed of 35.67 mol% DSPC, 33.3 mol% cholesterol, 25 mol% DSPE-PEG(2000) carboxylic acid and 6 mol% DSPE was prepared in a 50 mL round bottom flask. For the drug-free formulation of liposomal irinotecan (LIPO-0.3% PEG*), a lipid mixture composed of 53.8 mol% DSPC, 39.9 mol% cholesterol, 0.3 mol% PEG-PE and 6 mol% DSPE was prepared in a 50 mL round bottom flask. Lastly, for the drug-free formulation of liposomal doxorubicin (LIPO-5% PEG*), a lipid mixture composed of 49 mol% soy PC, 40 mol% cholesterol, 5 mol% PEG-PE and 6 mol% DSPE was prepared in 50 mL round bottom flask. Methanol was added dropwise to all flasks until each mixture was clear. Lipid film generation, rehydration, extrusion, and dye labeling steps are described in the base liposome synthesis section. For the extrusion step, we note that the described PEGylated formulations were passed through one 400 and one 200 nm membrane followed by two passes through one 100 nm membrane.

#### PLGA Nanoparticle Synthesis

PLGA was dissolved at a concentration of 10 mg/mL in acetone and Cy5 free acid dye was dissolved at a concentration of 50 mg/mL in DMSO. 6 mL of milliQ water were added to a 20 mL scintillation vial and stirred gently on a plate. In a 2 mL Eppendorf tube, 2 ul of the dye was added to 1 mL of the PLGA solution, mixed and drawn into syringe with a 27-gauge needle attached. In the scintillation vial, the tip of the needle was submerged below the water line and the PLGA-Cy5 solution was slowly added to the water under constant stirring. The solution was left to stir 3 hours to allow for solvent evaporation. An additional 2 mL milliQ water were added the solution prior to purification using tangential flow filtration (as described previously).

#### Synthesis of Layer-by-Layer Nanoparticles

Liposomes and PLGA nanoparticles were layered by adding an equal volume of nanoparticle solution (not exceeding 1 mg/mL) to an equal volume of polyelectrolyte solution under sonication at room temperature. The mixture was sonicated for roughly 3 seconds. The optimal weight equivalent (wt. eq.) for each layer was determined through a polyelectrolyte titration using 50 uL samples of the nanoparticle for each tested wt. eq. Each test ratio was layered as described above and then characterized. If the resulting particle had a zeta potential greater than 30 mV or less than -30 mV, and an acceptable size, it was chosen as the optimal wt. eq. for each layer. The wt. eqs. of the cationic first layer, PLR, were 0.3 for the liposome core and 0.4 for the PLGA core. For the anionic second layer, the same weight equivalents of polyelectrolyte were used for both the liposome and PLGA core. The weight equivalent of each polyelectrolyte layer are as follows: 0.65 wt. eq. PLD, 0.65 wt. eq. PLE, 1.2 wt. eq. HA, 0.65 wt. eq. dextran sulfate, 1.2 wt. eq. fucoidan, 1.2 wt. eq. alginate, 0.2 wt. eq. PAA, and 1.2 wt. eq. chondroitin sulfate. Polyelectrolyte solutions for liposome layering, except for HA and alginate, were prepared in 50 mM HEPES and 40 mM NaCl (pH 7.4) which was diluted to 25 mM HEPES and 20 mM NaCl upon 1:1 mixing with the nanoparticle substrate in water. HA and alginate stocks were prepared in 10 mM HEPES which was diluted to 5 mM HEPES upon mixing with the nanoparticle substrate in water. All polyelectrolyte solutions for PLGA nanoparticle layering were prepared in water. Layered particles were incubated at room temperature for one hour before being purified via TFF and characterized.

#### Synthesis of PEG-PLGA Nanoparticles

PEG-PLGA (50:50, 10k PLGA, 2k PEG) was dissolved at a concentration of 10 mg/mL in a 1:1 ratio of acetone to DMSO and cyanine5 free acid dye was dissolved at a concentration of 50 mg/mL in DMSO. 6 mL of milliQ water was added to a 20 mL scintillation vial and stirred gently on a plate. In a 2 mL Eppendorf tube, 1 μL of the dye was added to 0.5 mL of the PEG-PLGA solution, mixed and drawn into syringe with a 27-gauge needle attached. In the scintillation vial, the tip of the needle was submerged below the water line and the PEG-PLGA-cyanine5 solution was slowly added to the water under constant stirring. The solution was left to stir 3 hours to allow for solvent evaporation. An additional 2 mL milliQ water were added the solution prior to purification via TFF and characterization.

#### PEGylation of Carboxylated Polystyrene Nanoparticles

20, 100 and 200 nm yellow green carboxylated fluospheres were prepared at stocks of 2% solids and 40 nm yellow green carboxylated fluospheres were prepared at a stock of 5% solids. A 5 mg/mL stock of 3k PEG-NH_3_Cl was prepared in DPBS and a 15 mg/mL stock of EDC was prepared in PBS. For the 20, 100 and 200 nm fluospheres, 200 μLs of 3k PEG-NH_3_Cl and 100 μLs of each fluospheres were combined and mixed in separate 1.5 mL Eppendorf tubes. For the 40 nm fluosphere, 200 uLs of 3k PEG-NH_3_Cl and 40 uLs of fluospheres were combined and mixed in a 1.5 mL Eppendorf tube. To each Eppendorf tube, 200 μL of the EDC solution was added. All reactions were carried out a room temperature and protected from light for 6-8 hours while mixing.

#### Fluorescently Tagging Antibodies

The cetuximab antibody was prepared in 0.1M sodium bicarbonate (pH 8.2) at a concentration of 0.5 mg/mL and the isotype IgG antibody was prepared in water at 2 mg/mL. SulfoCy5 NHS ester dye was prepared in DMSO at a concentration of 50 mg/mL. 200 μL of the cetuximab antibody was added to a protein lobind tube. In a separate protein lobind tube, 50 μL of the IgG antibody was added with 150 μL of 0.1M sodium bicarbonate (pH 8.4). To each solution, 1 μL of sulfoCy5 NHS ester dye was added. Both reactions were carried out a room temperature and protected from light for 6-8 hours while mixing. To purify the antibodies, a 3k MWCO spin column was used. Each antibody was washed 15 times to purify.

#### Conjugation of Antibodies to Nanoparticles

EDC and sulfoNHS stocks were prepared at 10 mg/mL in water. Isotype IgG and cetuximab antibodies were prepared at 0.2 mg/mL in PBS. 25% PEG liposomes (LIPO-25% PEG) were prepared at 1 mg/mL in water. 1 mL of nanoparticle was added to two separate protein lobind tubes. To each tube, 8 uL of EDC and 16 uL of sulfoNHS was added. Each tube was mixed at room temperature and protected from light for 30 minutes. 0.5 mL of isotype IgG was added to one lobind tube and 0.5 mL of cetuximab was added to the other lobind tube. Each reaction was mixed at room temperature and protected from light for 1 hour. After an hour, tangential flow filtration was used for purification and nanoparticles were characterized via DLS.

#### Characterization of Nanoparticles

Nanoparticle hydrodynamic size and polydispersity were measured using dynamic light scattering (Malvern ZS90 Particle Analyzer). Zeta measurements were also acquired with the Malvern ZS90 using laser doppler electrophoresis. Nanoparticle solutions were diluted in milliQ water in polystyrene, semi-micro cuvettes for size measurements and DTS1070 folded capillary cuvettes for zeta measurements. For LIPO-5% PEG, LIPO-25% PEG, LIPO-5% PEG*, LIPO-EGFR and LIPO-IgG, the hydrodynamic size for each nanoparticle was measured using high throughput dynamic light scattering (Wyatt Dyna Pro Plate Reader) with samples diluted in milliQ water and tested in a black, glass bottom 384 well plate.

#### Pooled PRISM Cell Dosing with NPs and Preparation for Flow Cytometry

Cells were seeded at 200,000 cells/well in 0.5 mL RPMI-1640 media supplemented with 10%FBS in a 12-well plate. Cells were allowed to grow for 24 hours prior to treatment with nanoparticles. Prior to dosing, all PLGA nanoparticle formulations were normalized to a concentration of 50 μg/mL and all other nanoparticle formulations were normalized to a concentration of 100 μg/mL. Cells were dosed with 50 μL of normalized PLGA nanoparticles and 25 μL of normalized nanoparticles for all other formulations (Table S3). Cells and nanoparticles were incubated for either 4 or 24 hours at 37°C and 5% CO_2_.

After incubation, cells were washed once with 500 μL of warm PBS and dissociated with 150 μL 0.25% Trypsin-EDTA. After 5 minutes at 37 °C, the trypsin was quenched with 200 μL of media and the cells were triturated vigorously to ensure that all cells had been dissociated from the plate. Cells were then transferred to a fluorescence-activated cell sorting (FACS) tube through a cell strainer cap and placed on ice until sorted.

#### Pooled PRISM Cell Dosing with Antibodies

10 wells of cells were washed twice with 200 μL of room temperature PBS and dissociated with 200 μL of accutase by incubating at 37°C for 5 mins. After incubation, 300 μL of cold FACS buffer (PBS + 2% FBS) was added to each well and the cells were triturated. Each well was transferred and combined in a 15 mL falcon tube and spun at 1000 rpm for 5 minutes at 4°C. After spinning, the supernatant was removed and the cells were counted and resuspended in FACS buffer at a concentration of 1e6 cells/mL. 200 μLs of the cell suspension was transferred to 12 separate FACS tubes. To three of the tubes, nothing was added to remain as an untreated control. To three tubes, 15 μL of 0.1 mg/mL Cy5-cetuximab was added. To three tubes, 15 μL of 0.1 mg/mL Cy5-IgG was added. To the final three tubes, 5 μL of EGFR-AF488 (used at undiluted stock concentration provided by manufacturer) were added. All tubes were vortexed gently and incubated in the dark at 4°C for one hour. After an hour, 200 μL of cold FACS buffer was added to each tube, the cells were triturated and then spun at 1000 rpm for 5 minutes at 4 °C. After centrifugation, the supernatant was removed and the cells were resuspended in 300 μL of cold FACS buffer. The cells were stored on ice until flow sorting.

#### Preparation of Untreated and Sorting Control Samples

Cells were washed once with warm PBS. For cells that were lysed in well, 150 μL of lysis buffer was added to each well and cells were incubated for 5 minutes at 37 °C. After incubation, 100 μL of PBS was added and the lysed cells were triturated and transferred to an Eppendorf tube. For trypsinized cells, after a PBS wash, 150 μL 0.25% Trypsin-EDTA was added to the wells and incubated for 5 minutes at 37°C. To quench the trypsin, 200 μL of media was added to the well and the cells were triturated and transferred to either a FACS tube (for sorted control) or Eppendorf tube (for unsorted control). The unsorted control cells were pelleted by centrifuging at 1000 rpm for 5 minutes and resuspended directly into 150 μL of lysis buffer.

#### Flow Cytometry and FACS Information

For FACS, samples were sorted using a BD FACSAria II Cell Sorter (BD Biosciences). Samples dosed with Cy5 nanoparticles or Cy5 antibodies were sorted on the APC channel (ex. 640, filters 660/20). Samples dosed with yellow green fluospheres or AF-488 antibodies were sorted on the GFP channel (ex. 488, filters 530/30).

For all flow analysis in validation studies, samples were analyzed using a BD LSR II Flow Cytometer with high throughput sampler (BD Biosciences). Samples dosed with Cy5 nanoparticles were analyzed on the APC channel (ex. 640, filters 670/30). Samples dosed with yellow green fluospheres were analyzed on the GFP channel (ex. 488, filters 515/20). Data was analyzed using FlowJo (version 10), and cells were gated for single cells based on an untreated, parental cell line for each condition using the side scatter and forward scatter plots; singlet gates were applied to all samples of the same parental cell line. Analysis of NP intensity was based on a single color (APC or GFP, as above) without compensation.

#### PRISMseq

Samples lysed in DNA Lysis Buffer (20mM Tris-HCl (pH 8.4), 50mM KCl, 0.45% NP-40, 0.45% Tween-20, 10% Proteinase K) were denatured at 95 °C and amplified with a 2X KAPA polymerase master mix. Custom primers (IDT) allowed samples to be dual-indexed for multiplexed Illumina sequencing by directly adding Illumina flow-cell binding sequences to the amplicon:

forward primer: 5’AATGATACGGCGACCACCGAGATCTACANNNNNNNNAAGGTGCTTCTCGATC TGCAT
reverse primer: 5’CAAGCAGAAGACGGCATACGAGATNNNNNNNNGTGACTGGAGTTCAGACGT GTGCT.

where N represents the indexing nucleotides. Resulting products were quality control checked for single-band amplification using gel electrophoresis and then pooled and purified for sequencing using the Zymo Select-a-Size DNA Clean & Concentrator kit. After pooling, the PCR product was quantified using the Qubit 3 Fluorometer. Samples were sequenced using Illumina HiSeq technology. Samples were loaded onto the HiSeq flow cell at a final concentration of 10pM with a 20% PhiX spike-in due to low diversity. Sequencing was run for 50 cycles, single-read.

#### SLC46A3 Validation Studies

##### Non-pooled screening

HCC1143 (RPMI-1640), HCC1395 (RPMI-1640), HeLa (RPMI-1640), SW948 (RPMI-1640), LOXIMVI (RPMI-1640), SJSA-1 (RPMI-1640), MCF7 (Eagle’s Minimum Essential Medium, EMEM), DAOY (EMEM), MDA-MB-231 (Dulbecco’s Modified Eagle’s Medium, DMEM), CAOV3 (DMEM), T47D (RPMI-1640), and HepG2 (DMEM) cells were seeded individually at 10,000 cells/well in 100 μL of media, supplemented with 10% FBS and 1X Penicillin-Streptomycin. Base media are provided in parentheses following cell line names. Cells were allowed to grow overnight prior to treatment with nanoparticles. Prior to dosing, all nanoparticle formulations were normalized to a concentration of 50 μg/mL. Cells were dosed with 10 μL of normalized nanoparticles. Cells and nanoparticles were incubated for either 4 or 24 hours at 37°C and 5% CO_2_.

After incubation, cells were washed once with 100 μL of warm PBS and dissociated with 20 μL 0.25% Trypsin-EDTA. After 5 minutes at 37°C, the trypsin was quenched with 180 μL of media and the cells were triturated vigorously to ensure that all cells had been dissociated from the plate. Cells were placed on ice until analyzed using high throughput sampler.

##### Transient silencing of SLC46A3

For flow cytometry assessment, T47D cells were seeded at 8,000 cells/well in 100 μL of RPMI-1640 media, supplemented with 10% FBS and 1X Penicillin-Streptomycin for 24 hours at 37 °C and 5% CO_2_. Lipofectamine RNAiMax Transfection (ThermoFisher) was used according to manufacturer instructions to formulate siRNA for 96 well plate dosing (1 pmol siRNA/well). Cells were treated with siRNA for 24 hours prior to NP addition (0.1 mg/mL, 5% well volume). 24 hours after NP treatment, cells were prepared for flow cytometry and analyzed as described above.

For PCR profiling, T47D cells were seeded at 400,000 cells/well in a 6 well plate. Lipofectamine RNAiMax Transfection Reagent (ThermoFisher) was used according to manufacturer instructions to formulate siRNA for 6 well plate dosing (25 pmol siRNA/well). Cells were treated with siRNA for 48 hours prior to washing, trypsin treatment, and pelleting.

#### *SLC46A3* Overexpression: Viral transfection of LOXIMVI cells

Lentiviral vectors were purchased from the Broad Institute’s Genetic Perturbation Platform (GPP), specifically ccsbBroad304_09945 (SLC46A3) and ccsbBroad304_99991 (Luciferase, vector control).

LOXIMVI cells were grown and passaged in RPMI-1640 supplemented with 10% FBS and 1X Pen/Strep until ready for infection. LOXIMVI cells were trypsinized, counted and resuspended to a concentration of 1.36×10^6^ cells/mL. A solution of 2X polybrene was added to the cell suspension such that the final concentration of polybrene was 8 μg/mL. Cells were seeded into two six-well plates at 750,000 cells/well. Lentiviral vectors were separately added to plates at six different doses: 0, 25, 50, 100, 200, and 400 μLs. After, 1 mL of media was added to each well and the cells were incubated overnight at 37 °C and 5% CO_2_ and media changed at 17 hours post-seeding. At 48 hours after seeding, the cells were re-seeded at 375,000 cells/well in 2 mLs of blasticidin containing media (final blasticidin concentration was 1 μg/mL). The selection progress was monitored via flow cytometry (Figure S12).

#### *SLC46A3* Knock-out via CRISPR-Cas9 in T47D cells

*SLC46A3* knock-out T47D cell lines were generated by infection with lentiCRISPRv2-Opti (Addgene # 163126) vectors encoding Cas9 and single guide RNAs (sgRNAs).(*67*) The following oligonucleotides were used for sgRNA cloning and include cloning overhangs for ligation after BsmBI digest of lentiCRISPRv2-Opti vector:

sgGFP_F: caccGGGCGAGGAGCTGTTCACCG
sgGFP_R: aaacCGGTGAACAGCTCCTCGCCC
sgSLC46A3_F: caccgAAAGCAAGCTCCCCAAAATG
sgSLC46A3_R: aaacCATTTTGGGGAGCTTGCTTTc

Clonal knock-out cell lines were isolated through fluorescence-activated cell sorting, and biallelic frame-shifts were confirmed by deep-sequencing (allele 1: -32 bp frameshift 501 reads; allele 2: -10 bp frameshift; 477 reads). The T47D *SLC46A3* knock-out line described has the mutant alleles c.442_453del and c.440_449del.

Quantification of SLC46A3 protein expression via western blot was not possible due to the lack of commercially available antibodies with proper specificity. In addition to our own experimental conclusions, this point has also been referenced in the literature.^48^

#### Quantitative PCR for *SLC46A3* transcript levels

RNeasy Plus Mini Kits for RNA extraction and QuantiTect Reverse Transcriptase Kits were purchased from Qiagen. β-mercaptoethanol was purchased from Sigma-Aldrich. Roche Light Cycler-DNA Master SYBR Green I mastermix and Corning Axygen 384-well PCR microplates were purchased through the MIT BioMicro Center / KI Genomics Core. IDTE buffer, nuclease free water, and PrimeTime PCR Primers were purchased from Integrated DNA Technologies (IDT). The primers have the following assay ID numbers and sequences:

HS.PT.39a.2214836 (GAPDH)

R: TGTAGTTGAGGTCAATGAAGGG
F: ACATCGCTCAGACACCATG
HS.PT.58.22528687 (SLC46A3)

R: GAACAGAGAATGGCACAATAGTG
F: ACGATGACAGGAATGGCTATG

Cells were pelleted and stored at -80 °C prior to RNA extraction. Total RNA was extracted according to the instructions provided with the Qiagen RNeasy kit. Briefly, lysis buffer was prepared with the recommended amount of β-mercaptoethanol to protect RNA from degradation. 1-3 million cells worth of lysate were added to spin columns. Total RNA was eluted from the columns using 30 µL nuclease-free water. Nanodrop (Thermo Fisher) spectrophotometry was used to assess RNA concentration and quality, and all 260/280 values were greater than 1.8. cDNA was synthesized according to manufacturer’s instructions using 1 µg of template RNA. cDNA was stored at -20° C or placed on ice for immediate use.

For qPCR reactions, cDNA was diluted 1:50 with nuclease-free water, and primers were diluted to 20x (10 µM) in IDTE buffer according to the manufacturer’s specifications. RT-qPCR was set up in a 384-well plate with 8 µL diluted cDNA, 10 µL 2x SYBR Green master mix, 0.8 µL nuclease-free water, and 1.2 µL 20x primer. Each condition was performed in technical triplicate. No primer (IDTE buffer instead of primer) and no cDNA (water instead of cDNA) controls were also used to ensure there was not contamination. RT-qPCR was run on a Lightcycler 480 (Roche) and Ct values obtained using the second derivative. The ΔCt method was used to compare expression between cell lines, normalizing to GAPDH. The ΔΔCt method was used to measure the effects of siRNA, knockout, and overexpression, normalizing to GAPDH. For siRNA treatment, expression was further normalized to that with scrambled siRNA treatment. For knockout and overexpression models, expression was further normalized to that of the parental cell lines.

#### Imaging Cytometry Sample Preparation

##### Figures 5A-F, S17

Engineered T47D and LOXIMVI cells were plated in T25 flasks in 5 mL of media at the following densities: 2.3-2.6 x 10^6^ cells/flask for LOXIMVI-vector control, LOXIMVI-SLC46A3 OE, and T47D-vector control cells; 3.8 x 10^6^ cells/flask for T47D-*SLC46A3* knockout cells. Cells were allowed to adhere for 24 h (37 °C, 5% CO_2_) prior to treatment with 250 μL NP solutions, ranging from 0.05-0.1 mg/mL core concentration, for 24 h.

For CellTracker/LysoTracker staining, the following staining solutions were prepared: 50 nM LysoTracker Green and/or 50 μM CellTracker Orange CMRA in serum-free RPMI-1640. CellTracker and LysoTracker concentrations were selected based on manufacturer recommendations as well as literature protocol.(*68*)

Cells were trypsinized and transferred to 15 mL falcon tubes prior to washing 2x with warmed PBS. Between washes, cells were pelleted at 300 rcf for 5 min. For CellTracker and LysoTracker staining, cells were re-suspended in 0.5 mL respective staining solution, and incubated in the dark at 37 °C for 60 min. Cells were then pelleted and washed 2x with warmed PBS.

For fixation, cell pellets were re-suspended in 2% formaldehyde in PBS, incubated on ice for 20 min, then washed 2x with PBS. Cell pellets were then re-suspended in Hoechst-3342 (1 μg/mL in PBS) for 2 min prior to washing 2x with PBS. Cells were re-suspended and stored in 2% FBS in PBS overnight prior to running samples on ImageStream.

Samples were run on an ImageStreamX Mark II(Luminex). Single color controls were prepared using the above protocol for compensation. For analysis, only the Cy5 (NP) and brightfield channels were used.

##### Figure S19-S20

Engineered LOXIMVI cells were plated in T175 flasks in 15 mL of media at 6 x 10^6^ cells/flask. Cells were allowed to adhere for 24 h (37 °C, 5% CO_2_) prior to treatment with 750 μL NP solutions (0.1 mg/mL core concentration) for 24 h.

Cells were trypsinized and transferred to 15 mL falcon tubes prior to washing 2x with warmed PBS. Between washes, cells were pelleted at 400 rcf for 5 min. For fixation, cell pellets were re-suspended in 2% formaldehyde in PBS (2 mL), incubated on ice for 20 min, then washed 2x with PBS. Cell pellets were then re-suspended in 5% normal goat serum + 0.025% saponin in PBS (1 mL) and incubated at room temperature for 1 hour. During this time, antibodies were prepared at the dilutions noted in table S4 in ice cold 1% BSA + 0.025% saponin in PBS. Cell solutions were divided up into respective antibody staining groups (SLC46A3 OE = 6 grouos, vector control = 5 groups) and transferred to a V bottom 96 well plate. Cells were pelleted at 400 rcf for 5 min prior to aspirating blocking solution and resuspending cell pellets in antibody solutions (0.1 mL) and incubating overnight at 4 °C.

Cells were washed 2x with PBS prior to resuspending cell pellets with secondary antibody solution and incubating at room temperature for 1 hour. Cells were then washed 2x with PBS, fixed with 1% formaldehyde in PBS (0.1 mL) for 5 minutes at room temperature, washed again with PBS 2x prior to re-suspending and incubating with Hoechst 33342 (1 μg/mL in PBS, 0.3 mL) for 2 minutes at room temperature.

Samples were run on an ImageStreamX Mark II(Luminex). Single color controls were prepared using the above protocol for compensation.

#### Imaging Cytometry Analysis Workflow

Data analysis was carried out using AMNIS IDEAS software (version 6.2). First, singlet cells were gated based upon scatter plots using brightfield images (Ch01), with aspect ratio intensity on the y-axis and cell area on the x-axis, as shown in Figure S11; the same gate was used for all samples. For cells treated with nanoparticles only, the built-in ‘Internalization; function in IDEAS software was used to generate a cellular distribution score with default settings (a brightfield mask with a 5% erosion applied to all singlet cells). For cells treated with nanoparticles and subsequently stained with endolysosomal antibodies, five built-in analyses were run using IDEAS software: colocalization between nanoparticle signal and antibody signal; internalization of nanoparticle signal; internalization of antibody signal; ‘spot counter’ for nanoparticle signal; ‘spot counter’ for antibody signal. In addition, the ‘feature finder’ tool was used to generate quantitative measures of texture, size, and signal strength (Figure S13). Select raw data was exported from IDEAS as ‘.fcs’ files, and FlowJo software was used to visualize the data.

#### Deconvolution Optical Microscopy

Chambered cover glass was coated with rat tail collagen (Corning, 300 μL of 50 μg/mL in 0.02N acetic acid). After 5 minutes, the wells were washed with room temperature PBS and allowed to dry in a sterile environment. Wells were stored at 4 °C up to one week prior to seeding cells in 300 μL media at the following densities: 5,000 cells/well for T47D-vector control cells, LOXIMVI-vector control and SLC46A3 OE cells; 5,500 cells/well for T47D-*SLC46A3* KO cells. The cells were allowed to adhere for 24 h (37 °C, 5% CO_2_) prior to treatment with 15 μL of a 0.1 mg/mL NP solution (Cy5 channel) for either 4 or 24 h. Then, cells were washed 3x with warm PBS before adding LysoTracker Green (130 nM final concentration) + CellTracker Blue CMAC (13 μM final concentration) solution, which was prepared right before use in phenol red-free RPMI 1640. Cells were incubated in the dark at 37 °C, 5% CO_2_ for 45 min prior to aspirating dye solution, washing 3x with warm PBS, and adding 300 μL phenol red-free RPMI 1640 to each well. The cells were imaged with the Applied Precision DeltaVision Ultimate Focus Microscope with TIRF Module (Inverted Olympus X71 microscope) equipped with 405, 488, 512, and 568 nm lasers. Images were acquired with a either a 60x (with enhanced magnification) or 100x objective. All images were acquired with OMX softWoRx software (Applied Precison/GE). Image LUTs were linearly adjusted to improve contrast using FIJI. Z slices were merged into Z projections as shown in Figures 5G-H and S12. For CellTracker signal, a single (bottom most) slice was interleaved with the Z projection of the NP and LysoTracker signal.

#### Animal Studies

All animal experiments were approved by the Massachusetts Institute of Technology Committee on Animal Care (CAC) and were conducted under the oversight of the Division of Comparative Medicine (DCM). Flank tumors of LOXIMVI-vector control and LOXIMVI-SLC46A3 OE cells were established with a subcutaneous injection of 0.5-1.0 x 10^6^ cells as a 1:1 mixture with MatriGel (Corning) and PBS to the right flank of NCr nude mice.

##### Nanoparticle Formulation

Cy5 labeled NPs (1 mg/mL, LIPO-0.3% PEG*) were formulated in 5% dextrose (sterilized by filtering through a 0.2 μM filter).

##### Intratumoral Injection Studies

For intratumoral (IT) studies, within genders, mice with established flank tumors were randomly assigned to either the 4 or 24 h dosing cohort (n = 10; for 4 hour time point n = 5 female + 5 male mice, for 24 hour time point, n = 6 female + 4 male mice/cohort). Four or 24 hours after injection, mice were imaged using the In Vivo Imaging System (IVIS) Spectrum whole animal imaging device (PerkinElmer) using ex = 640/em = 700 nm to capture Cy5 signal. Immediately following imaging, mice were humanely euthanized and tumors were excised and imaged again via IVIS. Tumors were then placed into pre-weighed tubes containing 1 mL PBS. Tumors were weighed and their weights recorded for normalization of tumor fluorescence by tumor mass.

Tumor tissue was embedded in OCT compound (Tissue-Tek) and frozen over dry ice prior to sectioning. Sectioning and hematoxylin and eosin (H&E) staining was performed by Koch Institute’s histology core facility.

To confirm animal gender did not confound our findings, using data obtained from the IT study, we compared the total radiant efficiency divided by tumor mass (mg) of male and female mice at both the 4 and 24 h time points, and we did not find a statistically significant difference in NP tumor accumulation (*p* > 0.05).

##### Intravenous Injection Studies

For intravenous (IV) studies, n = 10 for the SLC46A3 overexpressing group (n = 5 female + 5 male mice) and n= 9 for the vector control group (n = 5 female + 4 male mice). Nanoparticles were administered to mice using tail vein injections (3 total, spaced 24 hour apart). Four hours after the third and final injection, mice were humanely euthanized and tumors were excised and imaged using the In Vivo Imaging System Lumina whole animal imaging device (PerkinElmer) to capture Cy5 signal (ex = 620/em = 670 nm). Tumors were then placed into pre-weighed tubes containing 1 mL PBS. Tumors were weighed and their weights recorded for normalization of tumor fluorescence by tumor mass.

#### Solid Lipid Nanoparticle (LNP) Formulation

Cholesterol, 1,2-distearoyl-sn-glycero-3-phosphocholine (DSPC), 1,2-dioleyol-sn-glycero-3-phosphoethanolamine (18:1 (Δ9-Cis) PE), and 1,2-dioleyol-sn-glycero-3-phosphoethanolamine-N-(Cyanine-5) (18:1 Cy5 PE) were dried from chloroform stocks under vacuum, then dissolved in 100% ethanol. Dilinoleylmethyl-4-dimethylaminobutyrate (DLin-MC3-DMA) and 1,2-dimyristoyl-*rac*-glycero-3-methoxypolyethylene glycol-2000 (DMG-PEG-2000) were dissolved in 100% ethanol. Lipid mixtures of the following molar compositions were prepared in Eppendorf tubes:

1. 50 mol % DLin-MC3-DMA, 38.5% cholesterol, 1.5% DMG-PEG-2000, 9.5% 18:1 (Δ9-Cis) PE, 0.5% 18:1 Cy5-PE; (4) 23 mol % DLin-MC3-DMA, 71% cholesterol, 1% DMG-PEG-2000, 5% DSPC;
2. 23 mol % DLin-MC3-DMA, 71% cholesterol, 1% DMG-PEG-2000, 5% DSPC;
3. 50 mol % DLin-MC3-DMA, 38.5% cholesterol, 1.5% DMG-PEG-2000, 10% DSPC.

mRNA was dissolved in 25 mM sodium acetate at 25 μg/ml. 4 volumes of mRNA were added to a 4-ml scintillation vial and stirred gently on a plate at room temperature. While stirring, 1 volume of lipid mixture in 100% ethanol was pipetted in rapidly. The solution was removed from stirring for 5 min. Then, the solution was re-stirred, and 5 volumes of DNAse/RNAse free water were pipetted in rapidly. The solution was then removed from stirring. Ethanol was allowed to evaporate overnight, and LNPs were resuspended to 10 ng/μL mRNA. LNP solutions were stored at 4C and dosed within 24 hours of preparation.

For characterization of LNPs, see **Characterization of Nanoparticles** and Table S5.

#### Lipid nanoparticle encapsulation efficiency

mRNA encapsulation efficiency was determined using the Quant-it RiboGreen RNA assay kit. Briefly, in a Nunc F96 MicroWell Black polystyrene plate, 5 μL mRNA-LNP samples were incubated in 45 μL either 1x TE or 0.5 (v/v)% Triton X-100 solution in 1x TE. Samples were incubated for 10 min, shaking at 100 rpm, at 37 °C. RiboGreen reagent was diluted 200-fold into 1x TE in 2-ml Eppendorf tubes, protected from light. Samples were then mixed with 50 μL diluted RiboGreen reagent. Then, samples were shaken at 300 rpm at room temperature, protected from light, for 5 min. Fluorescence intensities were read immediately on a Tecan M1000 plate reader, at an excitation of 485 nm and emission of 525 nm. Encapsulation was calculated as: (Fluorescence of Triton X-100 LNPs – Fluorescence of TE LNPs) / (Fluorescence of Triton X-100 LNPs).

#### Analysis of LNP-Cell Association and Transfection Efficacy

Cells were seeded individually in 96 well plates at 10,000 cells/well in 100 μL of media, supplemented with 10% FBS and 1X Penicillin-Streptomycin. Cells were allowed to grow 48 h prior to treatment with LNPs. Cells were dosed with 10 μL of normalized LNPs (100 ng mRNA).

After incubation for 24 h, cells were washed once with 100 μL of warm PBS and dissociated with 20 μL 0.25% Trypsin-EDTA. After 5 minutes at 37°C, the trypsin was quenched with 180 μL of media and the cells were triturated vigorously to ensure that all cells had been dissociated from the plate. 5 μL propidium iodide (PI, 50 μg/mL in PBS) were added to each well and incubated on ice for 15 minutes prior to flow analysis using a high throughput sampler.

Samples were analyzed in the GFP channel (ex. 488, filters 515/20), PE-Texas Red channel (ex. 561, filters 610/20), and APC channel (ex. 640, filters 670/30). Data was analyzed using FlowJo (version 10), and cells were gated first for PI negative populations based on untreated and heat killed controls, then for single cells based on an untreated cells from respective cell lines for each condition using the side scatter and forward scatter plots; singlet gates were applied to all samples of the same parental cell line.

#### Statistical Analysis

Methods pertaining to nanoPRISM analysis are detailed in **Supplementary Text**, below. All statistical analysis for non-pooled validation studies was performed using GraphPad PRISM 9. For single comparisons (non-parametric), the Mann-Whitney test was used. For multiple comparison testing, the Kruskall-Wallis test was used to compare treatment groups to the parental control.

## Supplementary Text

### nanoPRISM Probabilistic Model Development

In the statistical analysis of the sequencing data, a simple probabilistic model is employed to infer each cell’s probability from a given cell line to fall into the predefined bins after treating with a given NP formulation. In particular, for each NP treatment, the observed count for cell line *i* and bin *j* and technical replicate *k* is denoted with x_i,j,k_, and standard Poisson model for sequencing noise is assumed, i.e. *x_i,j,k_* ∼ *Pois*(*μ_i,j,k_*).

Furthermore, the expected value of this random variable, *μ_i,j,k_*, is factored into three operational quantities *μ_i,j,k_* = *λ_j,k_S_i_P_i,j_*, where *λ_j,k_* is a sample-specific scaling factor to take into the sequencing and PCR efficiency into account, *S_i_* is the initial abundance of cell line *i* before the treatment, and *P_i,j_* is the probability of each cell to fall into bin *j*.

In this formulation, we denote the control samples (not sorted into the bins) with a dummy bin- = 0 and *P_i,_*_0_ = 1for all *i*, and maximize the likelihood function

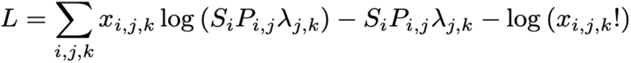

subject to the constraints 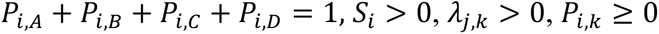, for all *i, j, k,* using a standard projected gradient descent algorithm to infer the binning probabilities *P_i,k_* independently for each biological sample and their concordance across replicates are used as a QC check. The R scripts used to process the data and infer the binning probabilities are available upon request.

### Univariate Analysis Description

For each given WA profile and each dataset in the cancer cell line encyclopedia (CCLE), we regressed the WA profile of the NP formulation on each column of the feature dataset and calculated the regression coefficient along with its corresponding standard error under the homogeneity assumption. Next, we applied the adaptive shrinkage method(*69*) to obtain moderated effect sizes, standard deviations and corresponding q-values. In the figures, the ratio of the posterior effect sizes and the standard deviations are presented as z-scores. This analysis was conducted in R(*70*) and figures were produced using the package ggplot2(*71*).

The methodology described above is available in the public github repo along with the documentation for how to use it:

#### https://github.com/broadinstitute/cdsr_models/blob/master/R/linear_association.R

### Random Forest Description

For the multivariate biomarker analysis, the weighted average of the binning probabilities scores, *W*’s, are used as the response variable and standard random forest regression models (RF) fitted proceeding a correlation-based feature selection. In particular, we fit two RF models for each NP formulation where the first one, *CCLE model*, uses a concatenation of the core cell line characteristics (mRNA expression, mutation status, copy number changes, and lineage annotations) as published in https://depmap.org, while the second one, *ALL*, includes more features (proteomics, CRISPR knock-outs, micro RNA, metabolomics) by limiting the analysis on the overlapping cell lines across all the datasets. For each model, 10-fold cross-validation is employed while in each fold an RF model fit after choosing the most correlated 500 features to the response variable). The cross-validated predictions then used to calculate Pearson scores (the correlation between the observed and predicted responses), and R^2^ values to assess the model performance. As the final step, these values are reported along with the default feature importances. The code used to fit the RF models is available at https://github.com/broadinstitute/cdsr_models.

### Principal Component Analysis and K-means Clustering

Principal component analysis (PCA) was performed using the weighted average of each nanoparticle-cell line pair from the by collapsing data by nanoparticle or cell-line. K-means clustering was performed on a subset of biomarkers generated by RF method that met the following criteria: CCLE features including gene expression, gene copy number, or protein abundance; univariate analysis z-score greater than 0. The Pearson correlation for biomarkers was then input for k-means clustering to generate 5 clusters. These analyses were conducted in R using ggfortify(*72, 73*) to perform PCA or k-means clustering and figures were produced using the package ggplot2(*71*).

## Supplemental Figures

**Figure S1.**
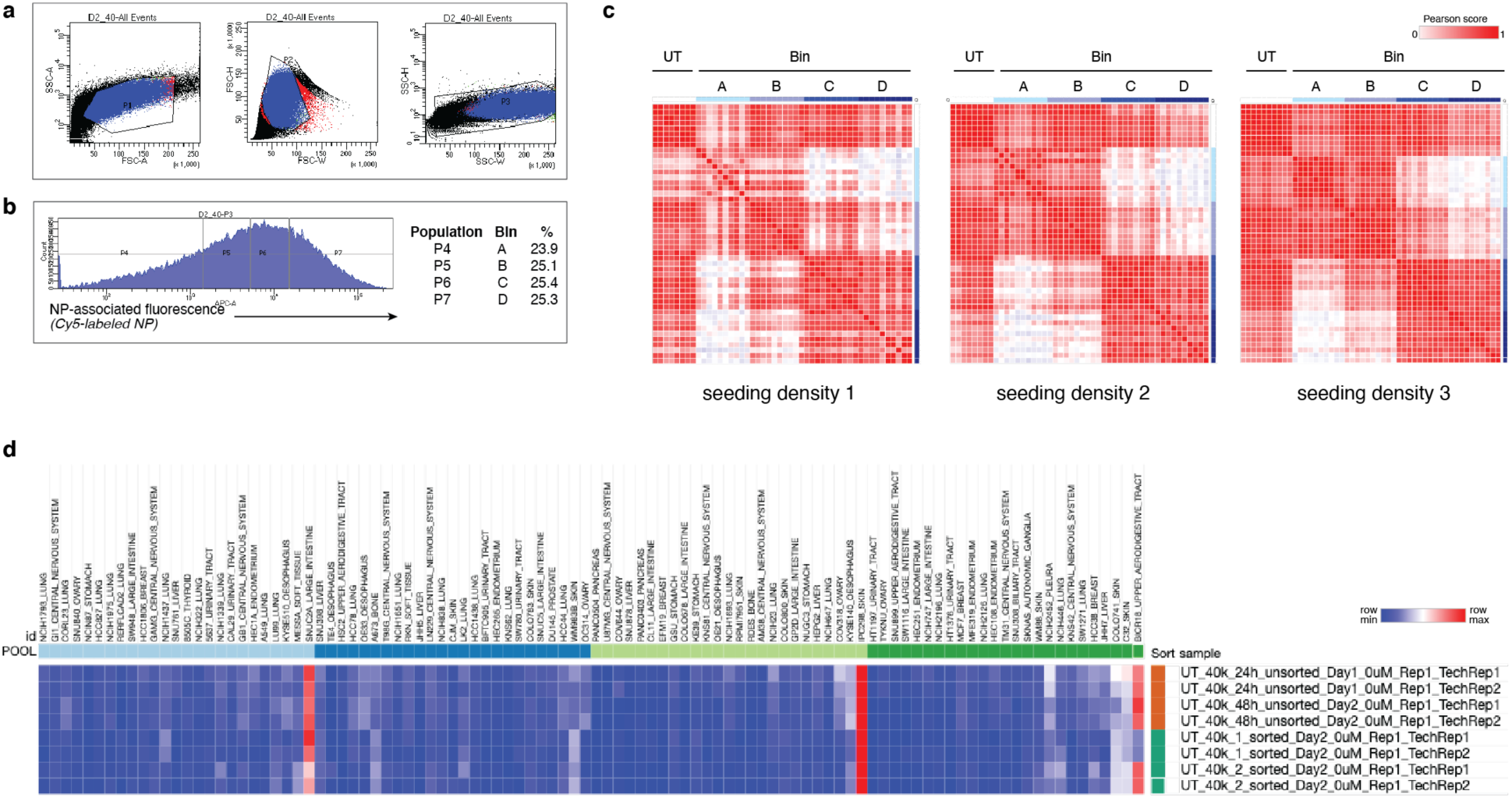
Optimization of nanoPRISM screening parameters using 100 pooled, barcoded cancer cell lines. (**A**) Representative data from pilot study showing gating strategy for pooled cells in order to remove debris and doublets. Black dots represent all events, P1 events are red, P2 green, P3 blue. Gate P1 removes debris by plotting events on a forward versus side scatter plot, and gates P2 and P3 remove doublets/aggregates using forward scatter and side scatter plots, respectively. Cells were pipetted through a cell strainer immediately prior to running so minimal aggregates are seen. (**B**) Events from P3 (non-debris, non-doublet cells) were either collected at that point as untreated (UT) controls or sorted into 4 bins based on nanoparticle (NP)-associated fluorescence, with an APC filter used to identify Cy5-specific fluorescence. Dynamic gating was used to obtain approximate quartiles. (**C**) Four sets of 25 cell lines were pooled together in a single well and lysed prior to sorting (‘unsorted’) or after sorting through the P1-P3 gates (‘sorted’), and the resulting populations were sequenced for barcode abundance. Each cell line was identified after sorting, and barcode abundance matched the unsorted controls, indicating that cell sorting did not induce a bottleneck. (**D**) In order to determine the effect of seeding density on cell sorting, three different seeding densities were tested and results were viewed using a similarity matrix with the hypothesis that untreated (UT) cells should be nearly identical and the low NP-association bins (A and B) would cluster separately from the high NP-association bins (C and D) such that A+B contain relatively distinct cell populations from C+D. Seeding densities 2 and 3 met this criteria, and seeding density 2 was most feasible for scale up to a larger screen.

**Figure S2.**
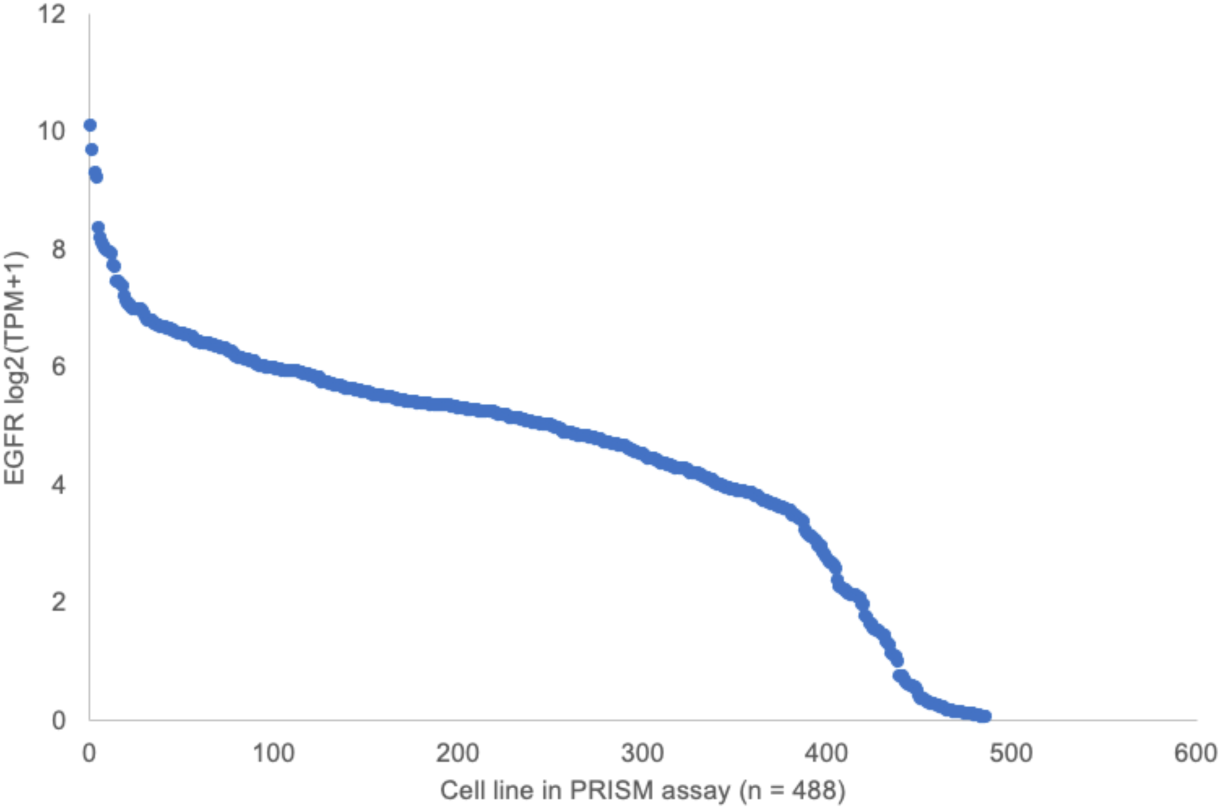
EGFR expression varies across the assayed cancer cell lines. EGFR data was obtained from the Dependency Map (DepMap).

**Figure S3.**
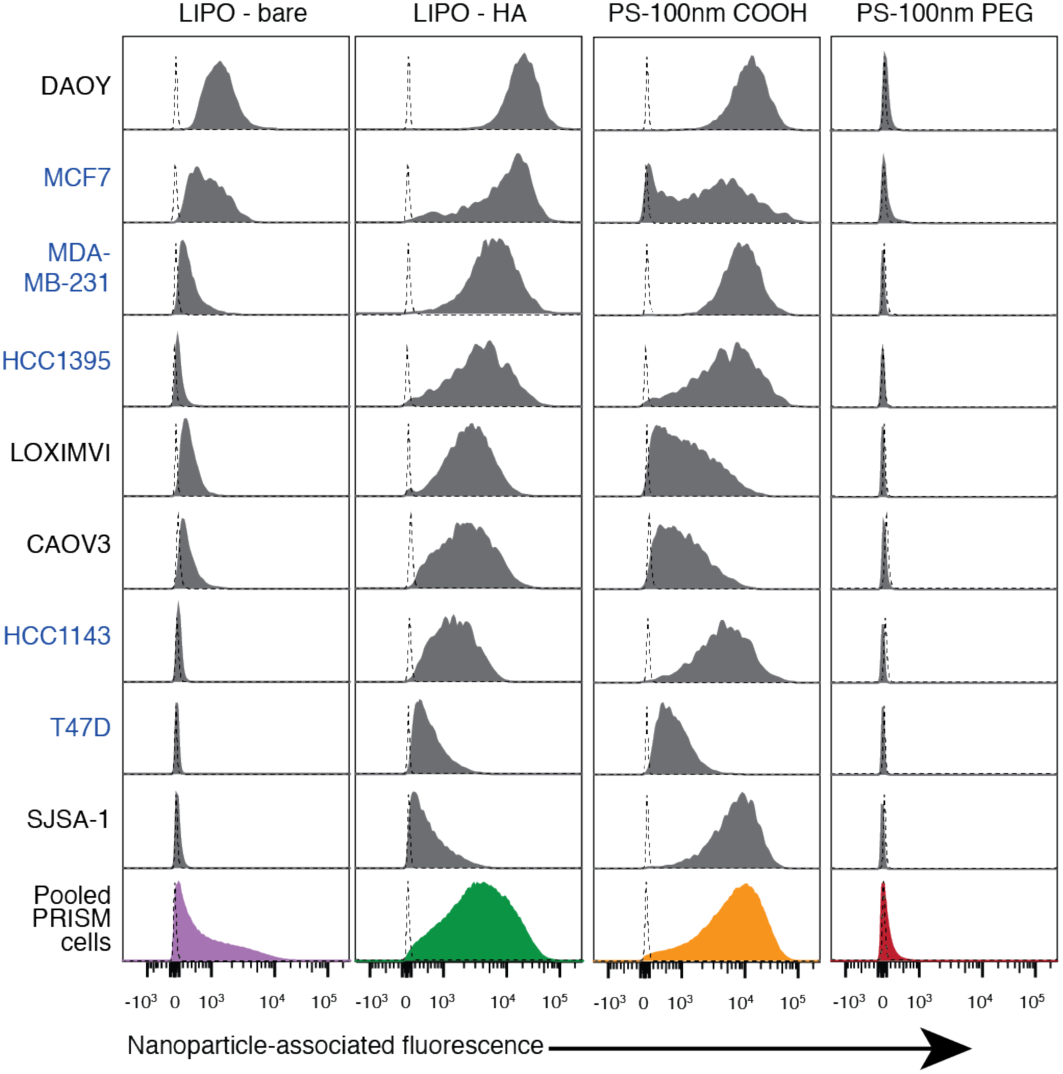
Flow histograms of PRISM cells and nine cancer cell lines treated with liposomal and polystyrene NP formulations. Blue text labels indicate breast cancer cell lineage.

**Figure S4.**
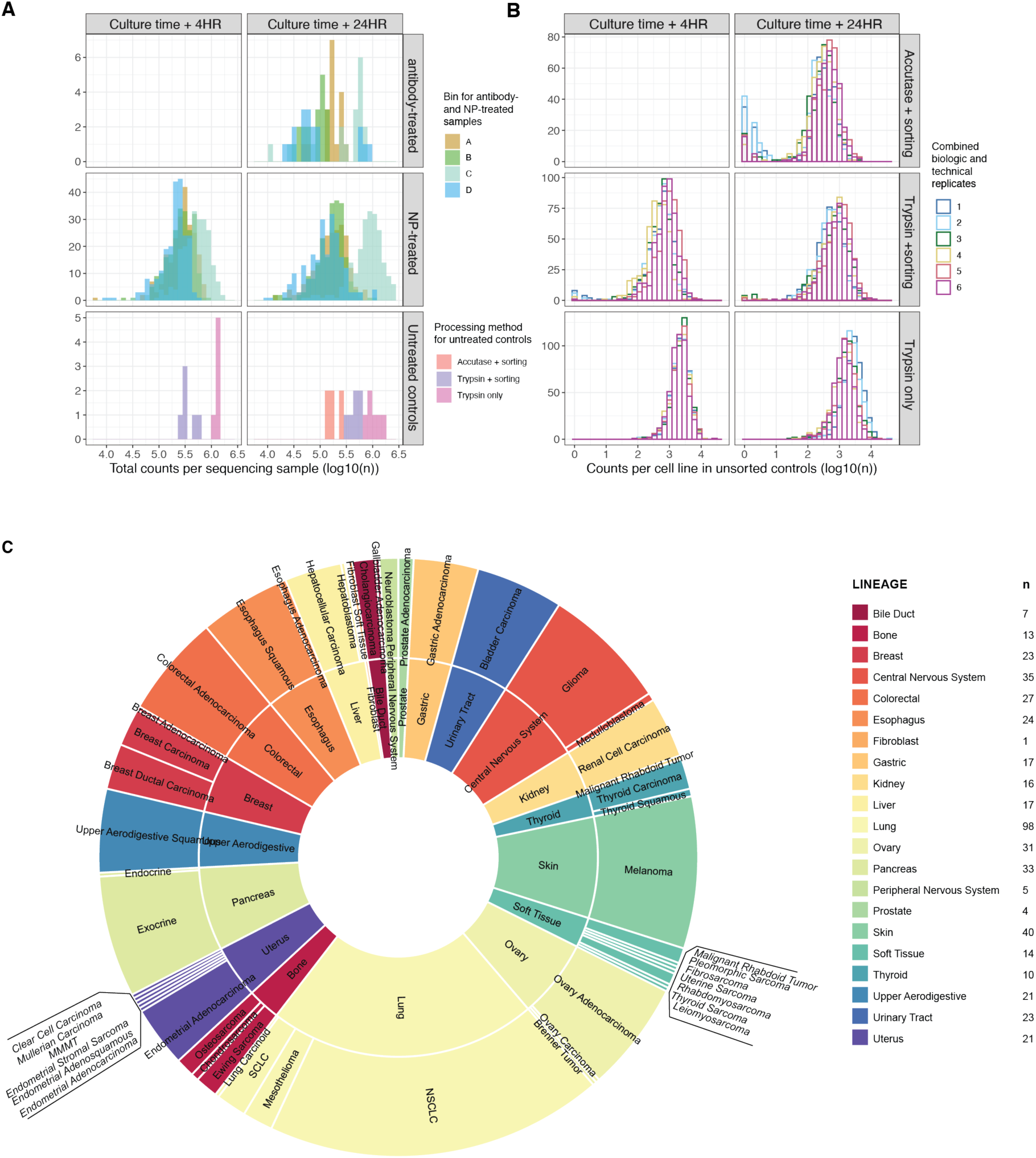
Quality control analysis of 488 cell lines prior to downstream analysis and breakdown of cell lines by lineage and sub-lineage. (A) Breakdown of Figure 1C detailing barcode counts for antibody- and NP-treated samples per bin as well as untreated controls (unbinned). For the untreated controls, see *Preparation of Untreated and Sorting Control Samples* for experimental details. For the accutase + sorting group, see *Pooled PRISM Cell Dosing with Antibodies* for cell preparation (without antibody treatment). **(B)** For the untreated controls, the distribution of counts across cell lines were examined for each biological (n = 3) and technical replicate (n = 2). **(C)** Sunburst plot showing the breakdown of the 488 cell lines by primary lineage (inner circle) and secondary lineage (outer circle) as defined by the DepMap portal.

**Figure S5.**
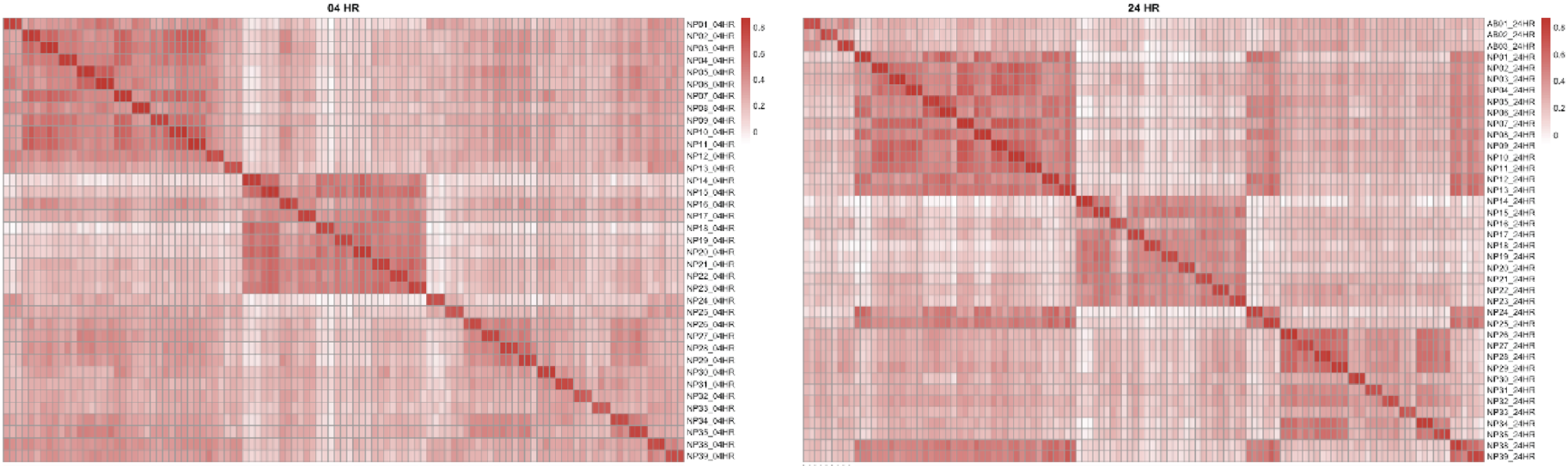
Visualization of biological replicates included in nanoPRISM. Rows represent individual nanoparticle formulations and columns indicate biological replicates (n = 3). Colors indicate Pearson correlation.

**Figure S6.**
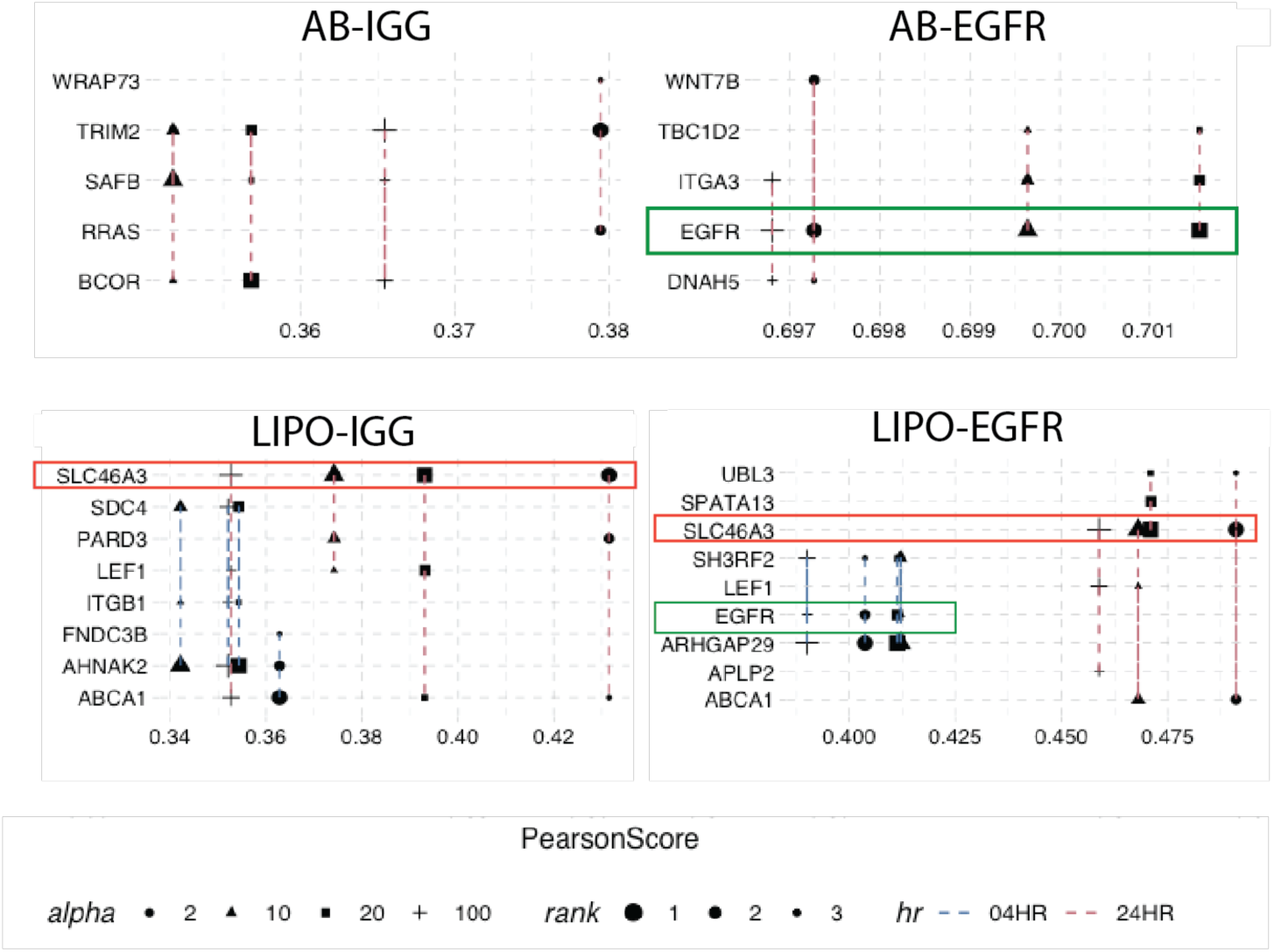
Visualization of biomarker rank (Pearson score) following weighted average alpha value trial. Random forest top ranked biomarkers for validation compounds are compared visually here using a range of alpha values. Green boxes highlight that the EGFR gene expression hit remains the top ranked random forest hit for the EGFR antibody (AB-EGFR) condition and the second ranked hit for the EGFR-conjugated liposome (LIPO-EGFR) over the range of alpha values tested, with minimal change in the Pearson score. The red boxes highlight gene expression of *SLC46A3*, which was found to be the top ranked random forest hit for all liposome formulations. Neither the rank or the Pearson score of this hit varied substantially across the range of alpha values tested.

**Figure S7.**
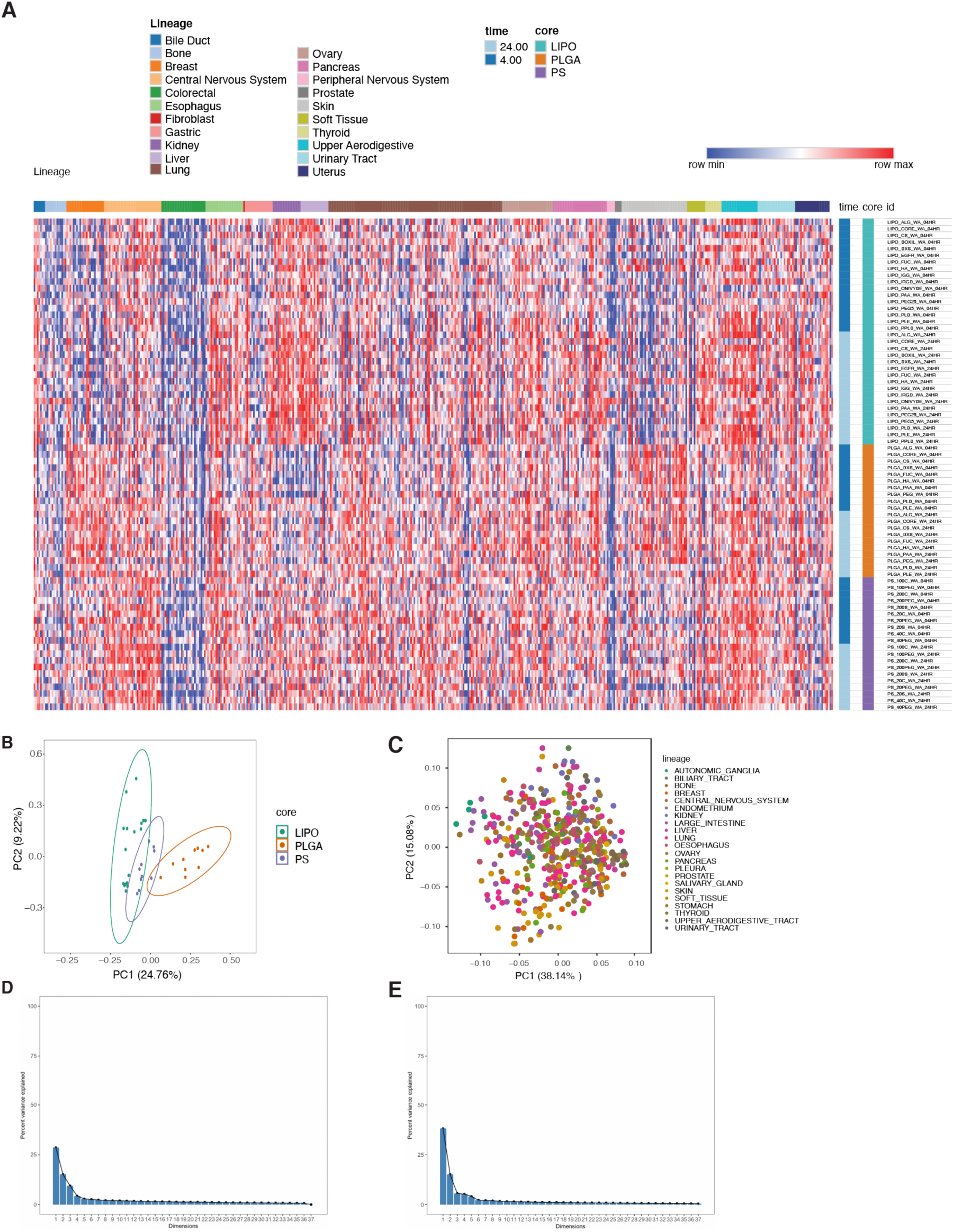
Heatmap of weighted average for all NP-cell line pairs at 4 and 24 h. Principal component analyses of NP-cell line weighted averages at 4 h, collapsed by NP core material. or by cell line lineage. (**A**) Raw data used to generate Figures 1F and S5B-C, displayed in heatmap form with each row representing a NP formulation and each column representing a cell line. (**B**) Principal component analysis with 4 hour data, collapsing weighted average data for all cell lines and (**C**) nanoparticle formulations, respectively. Scree plots showing the percentage of explained variance by dimension. D) 24hr data for weighted average by NP core and E) 24hr data for weight average by cell

**Figure S8.**
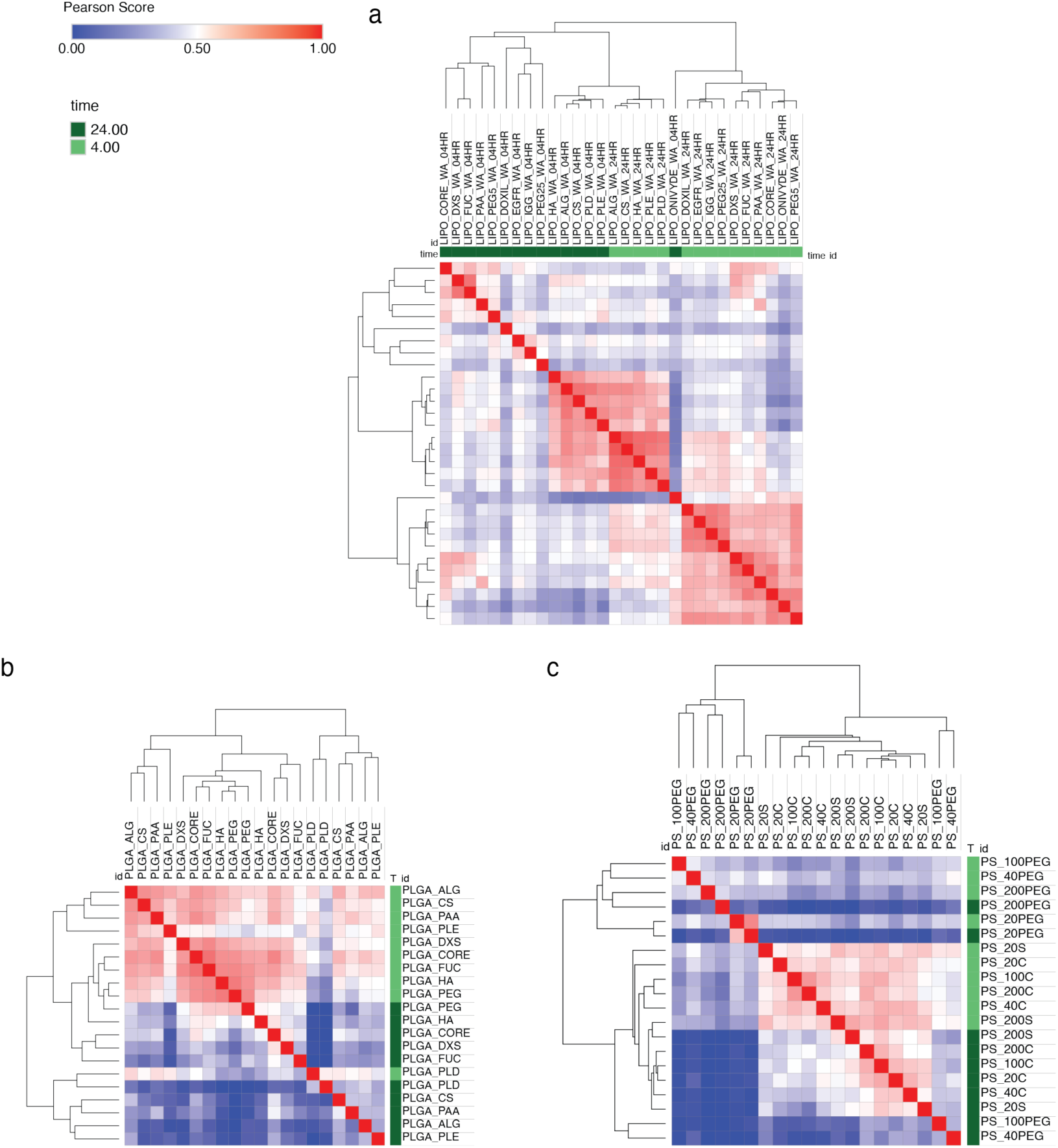
Similarity matrices separated by nanoparticle core material show time- and surface chemistry-dependent clusters. Each formulation is compared pairwise by Pearson score, with unsupervised hierarchical clustering shown for (**A**) liposome formulations (**B**) poly(lactic-co-glycolic acid) (PLGA), and (**C**) polystyrene (PS).

**Figure S9.**
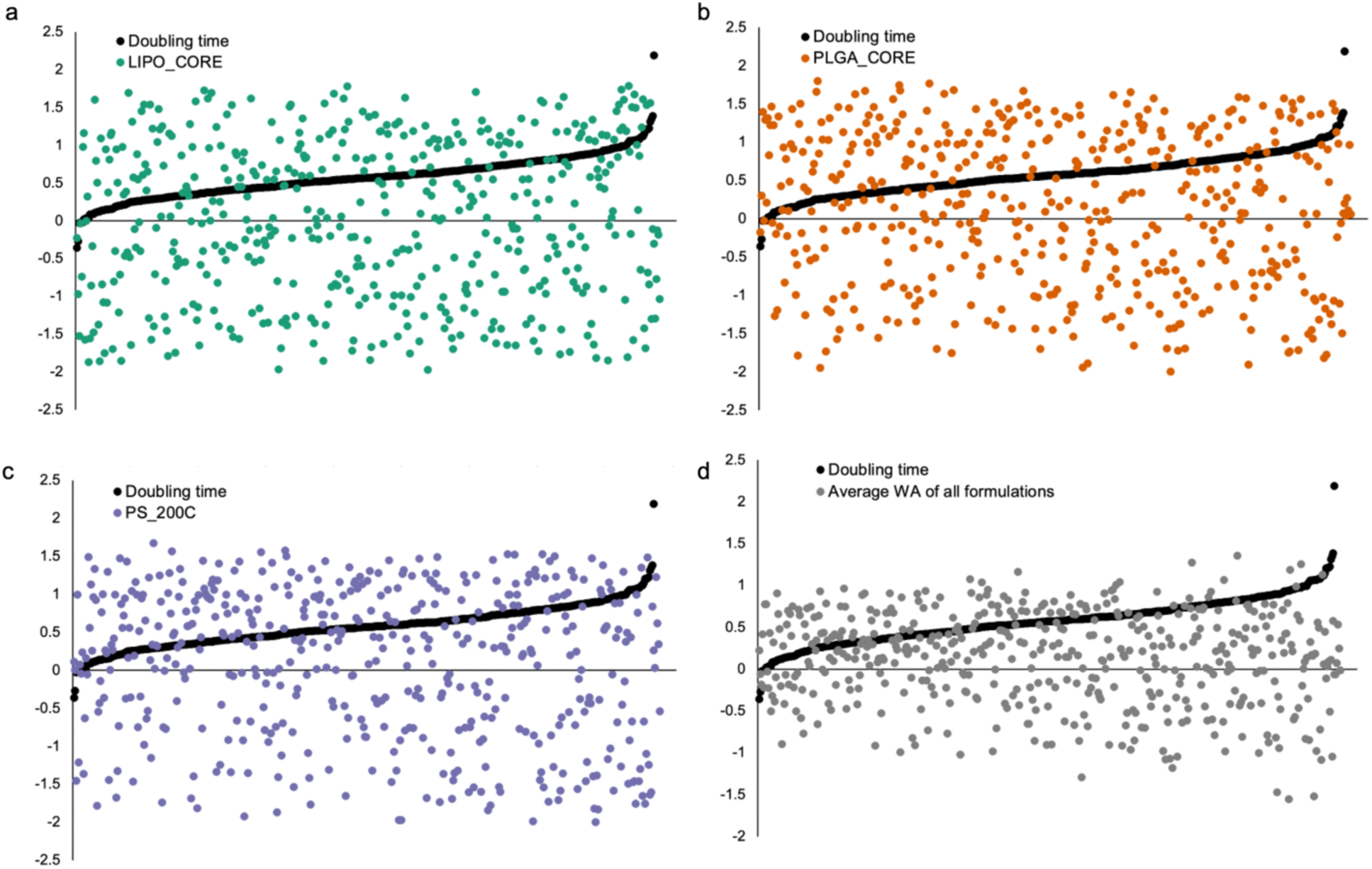
Comparison of weighted average (WA) values (24 h) and cell line doubling time. WA values and the Z scores of cell line doubling times (both represented on the y-axis) for each cell line were plotted. No visual trends between the two parameters were observed. Linear regression further revealed no meaningful correlation between WA and a cell line’s doubling time (r^2^ = 0.0054, 0.015, 0.020, and 0.012 for plots A-D, respectively).

**Figure S10.**
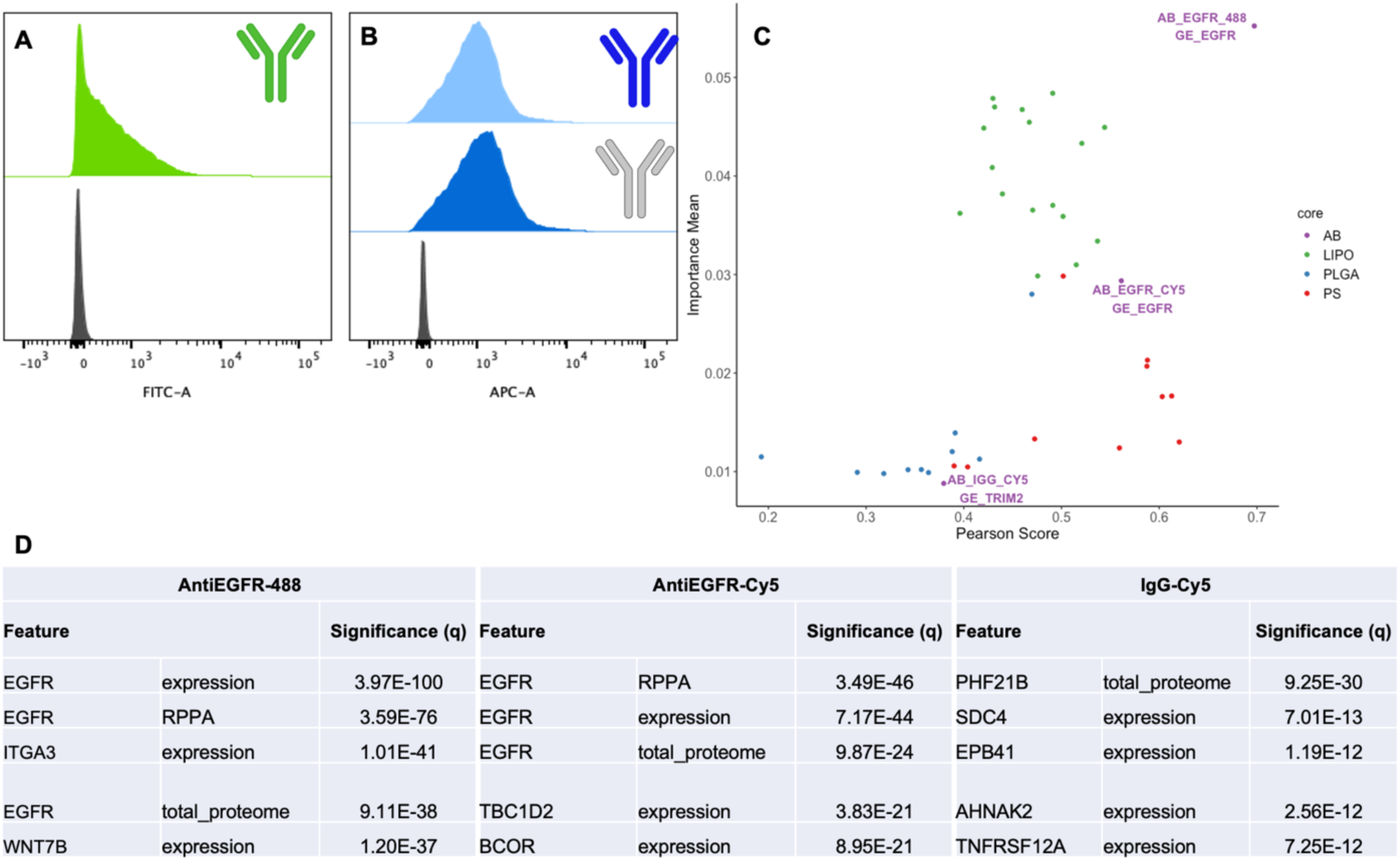
Biomarker identification is not dependent on fluorophore. (**A**) Flow histograms of PRISM cells treated with anti-EGFR conjugated to AF488 as well as untreated counterparts. (**B**) Flow histograms of PRISM cells treated with (top) anti-EGFR conjugated to Cy5, (middle) IgG isotype control conjugated to Cy5, and (bottom) untreated cells. (**C**) Plot illustrating the top ranked Random Forest feature associated with anti-EGFR and IgG treatments. (**D**) Summary table of top five most significant features for anti-EGFR and IgG and associated significance values.

**Figure S11.**
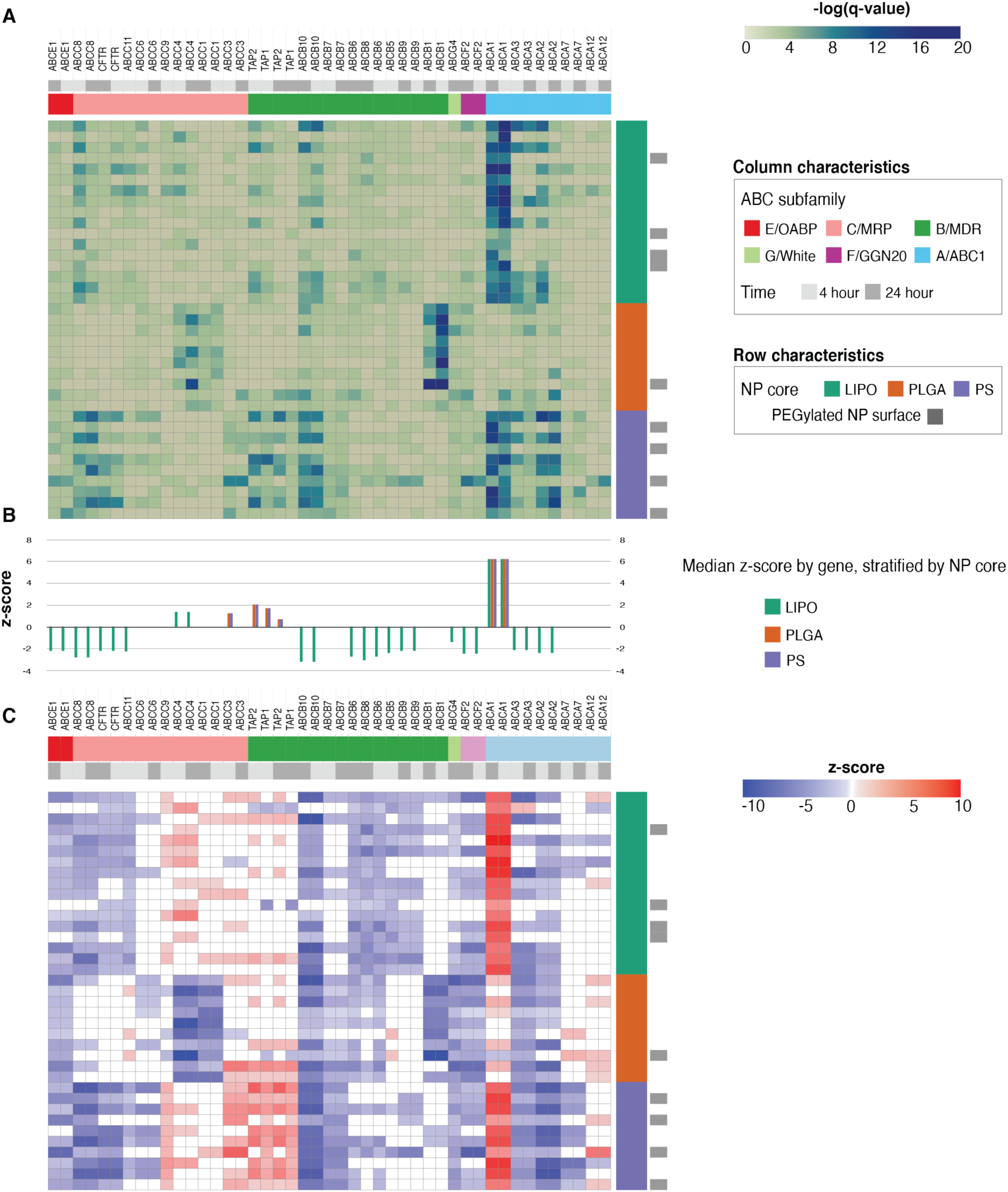
Investigation of ATP-Binding Cassette (ABC) transporter genes as candidate biomarkers for nanoparticle delivery. A) Heatmap showing genes in the ATP superfamily that were identified in this study in columns, annotated by subfamily and experimental time point. Rows correspond to nanoparticle formulations, annotated by core and presence or absence of surface PEGylation. The color indicates the significance of each gene/nanoformulation pair, with darker colors indicating higher statistical significance as quantifed by the -log(q-value). B-C) The directional relationship between gene expression and nanoparticle association is quantified by z-score with negative numbers indicating an inverse relationship between expression of the gene and nanoparticle association. In B z-scores are aggregated by nanoparticle core, and in C they are shown in heat map format.

**Figure S12.**
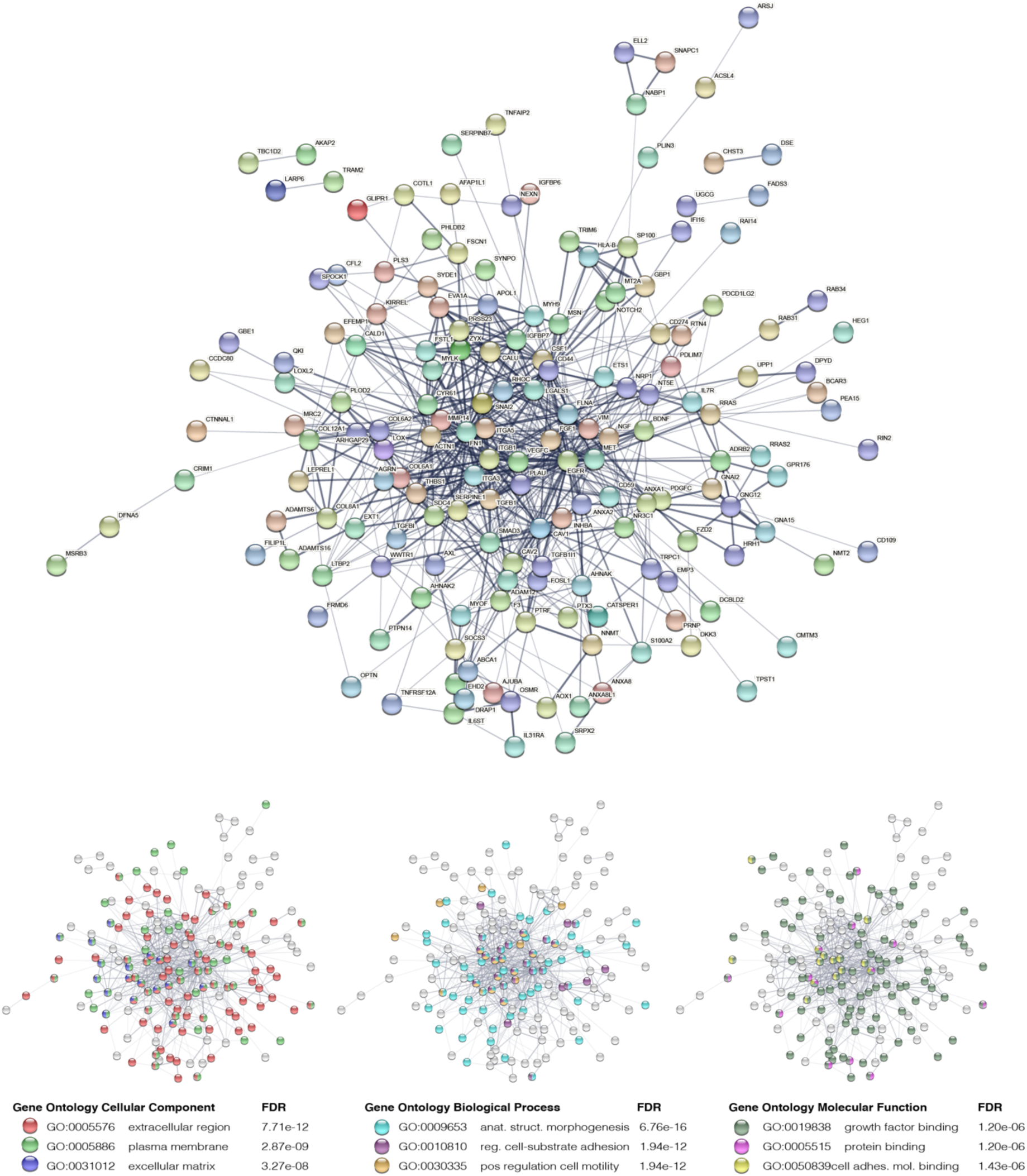
Visual representation of the STRING network represented in Figure 2E. The full network with protein names is represented in addition to gene ontology enrichments and false discovery rate (FDR) value.

**Figure S13.**
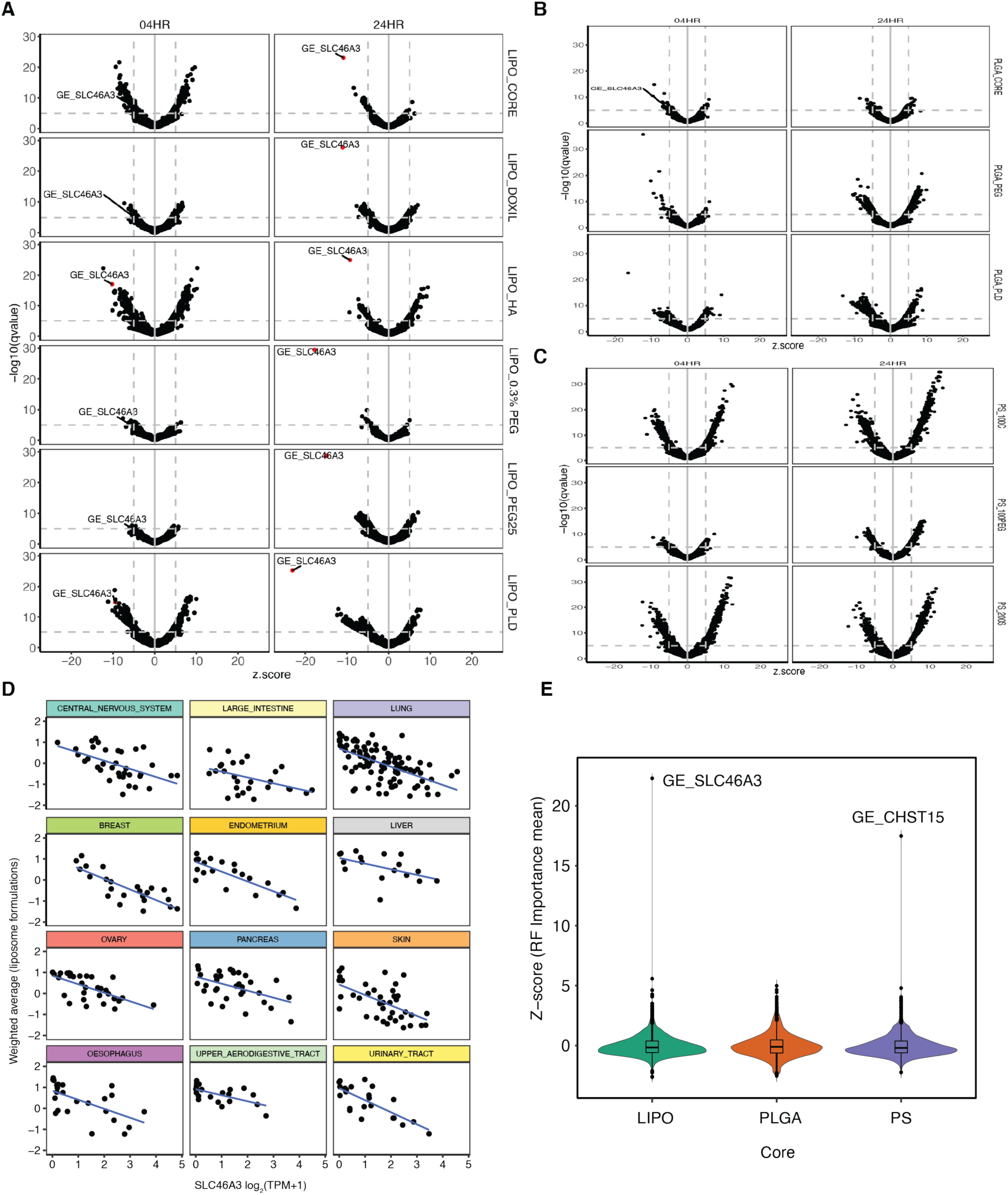
Expanded SLC46A3 univariate analysis and lineage analysis. (**A**) Gene expression of *SLC46A3* was a significant biomarker with negative z-score for liposome (LIPO) formulations with and without PEG and more significant at 24 h than at 4 h. Volcano plots for six liposomal NP formulations highlighting the position of SLC46A3 biomarker, which has higher significance and more negative z-score at 24 h compared to 4 h. SLC46A3 expression was not a top hit for (**B**) PLGA or (**C**) PS formulations at either time point. (**D**) The inverse relationship of *SLC46A3* expression was observed for tested LIPO formulations regardless of cancer cell lineage (24 h data shown). Lineages with representation of ≥ 15 cell lines were included. (**E**) Violin plot showing SLC46A3 as the most highly ranked Random Forest candidate biomarker in LIPO formulations. Top ranking for SLC46A3 is not observed in PLGA and PS formulations. NP formulations were collapsed by core and biomarkers Z-scores were generated using importance mean values from Random Forest. Biomarkers present in less than two formulations for a given core were excluded.

**Figure S14.**
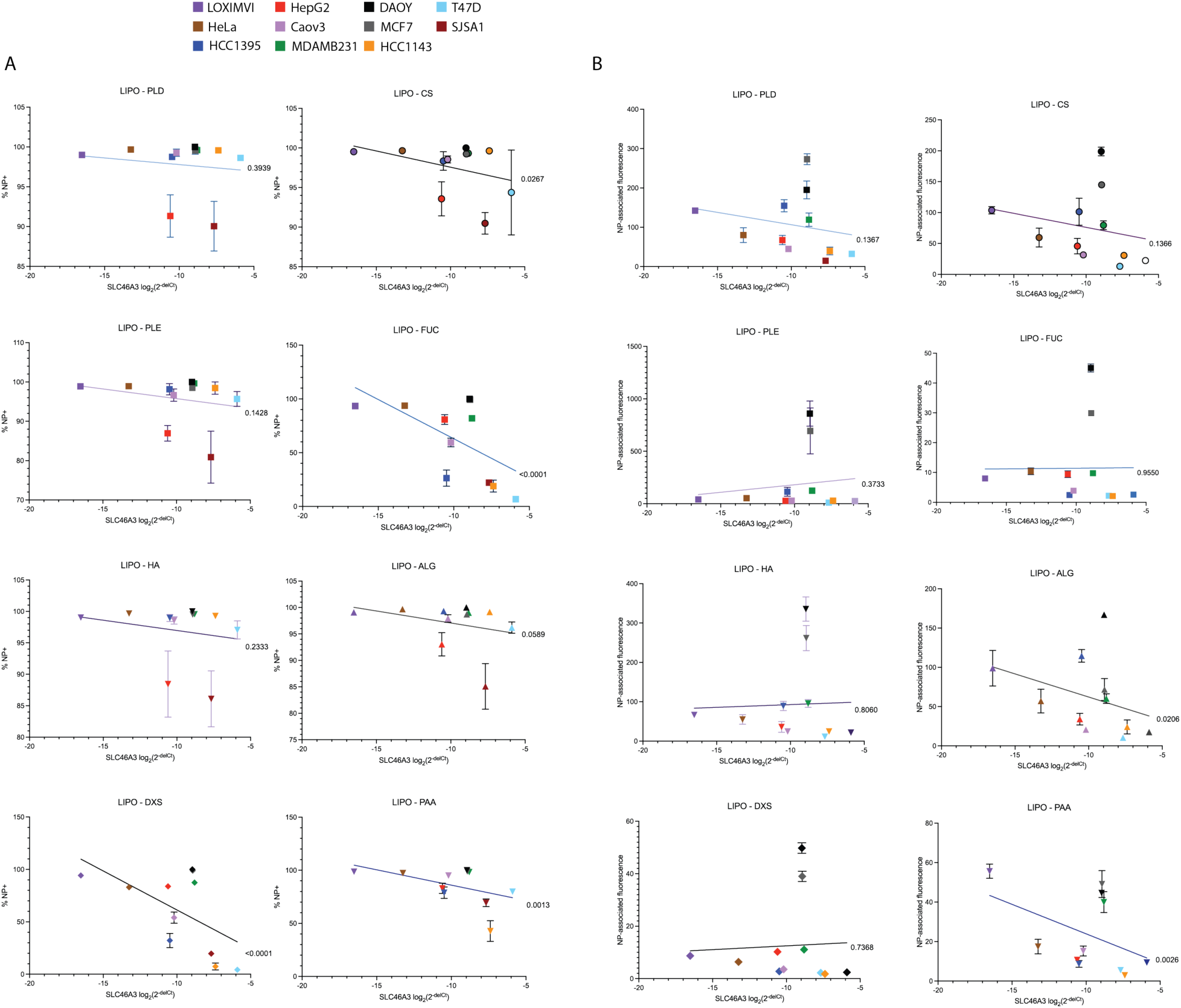
Expanded non-pooled screening via flow cytometry of cell interactions with Lipo NP formulations at 24 h reveals SLC46A3-dependent trends. Data is plotted as SLC46A3 expression data against (A) % NP cell population and (B) NP-associated fluorescence. Data is represented as the mean and standard deviation of four biological replicates. Error bars are not shown when smaller than data points. P values corresponding to the linear regression analysis for each data set are shown to the right of the respective fit lines. NP-associated fluorescence is defined as median fluorescence intensity normalized to untreated cells

**Figure S15.**
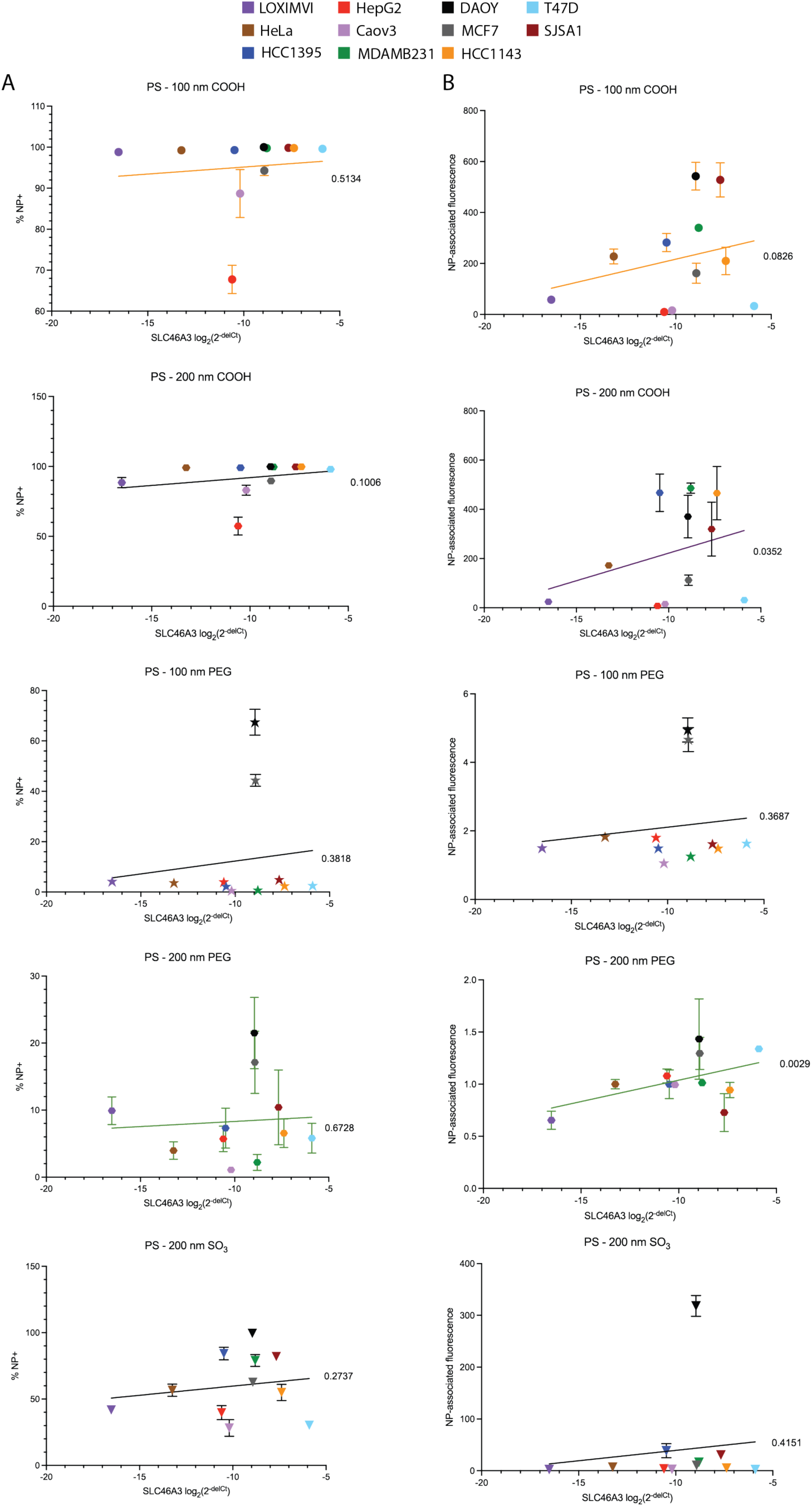
Expanded non-pooled screening via flow cytometry of cell interactions with PS NP formulations at 24 h reveals no consistent SLC46A3-dependent trends. Data is plotted as SLC46A3 expression data against (A) % NP cell population and (B) NP-associated fluorescence. Data is represented as the mean and standard deviation of four biological replicates. Error bars are not shown when smaller than data points. P values corresponding to the linear regression analysis for each data set are shown to the right of the respective fit lines. NP-associated fluorescence is defined as median fluorescence intensity normalized to untreated cells

**Figure S16.**
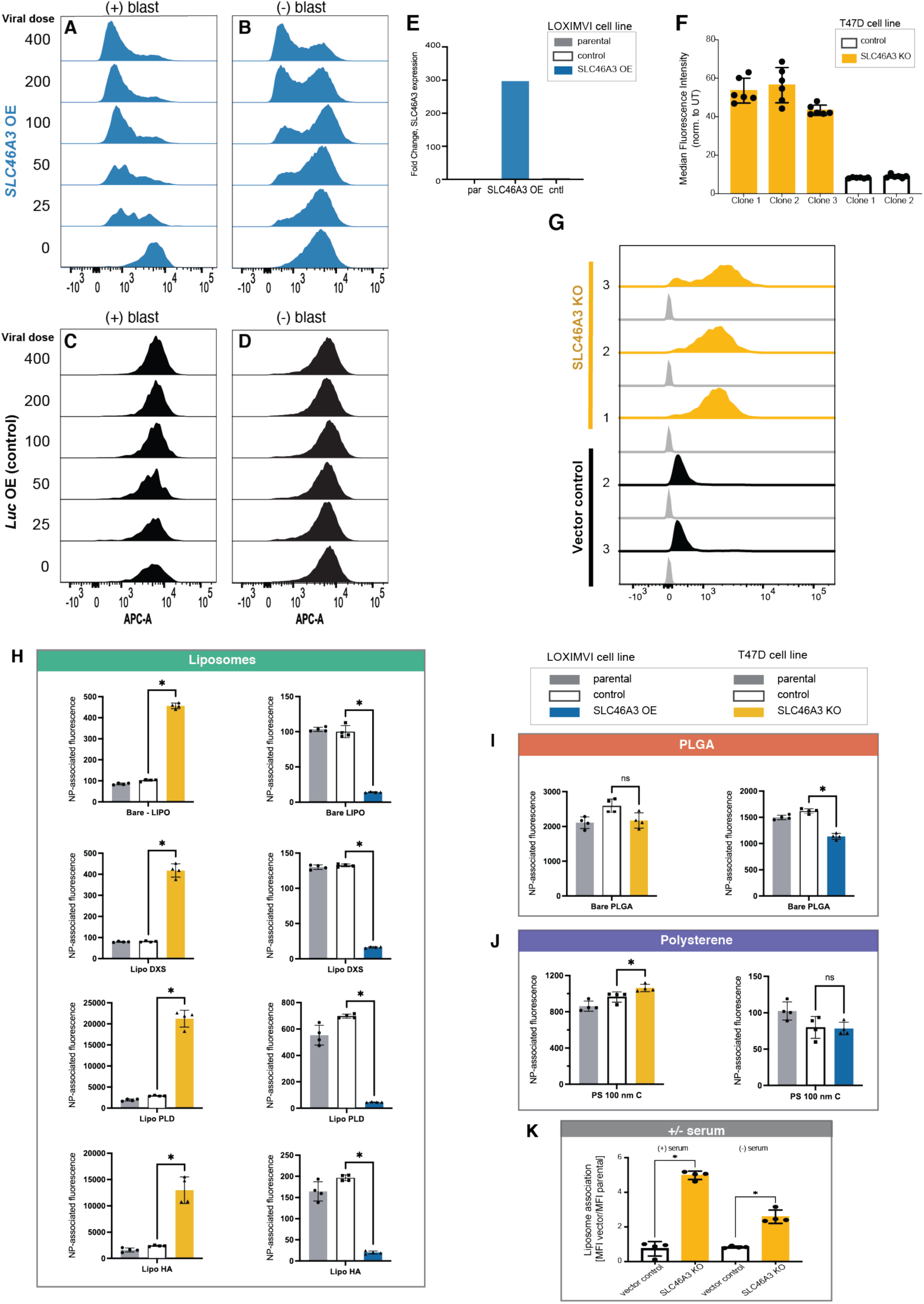
Flow cytometry profiling of engineered cell lines after *SLC46A3* modulation. After transduction (lentiviral dose, 0-400 μL), LOXIMVI were cultured (**A**, **C**) with and (**B**, **D**) without blasticidin (blast). Cells were then incubated with liposomal NPs (bare LIPO) for 24 h prior to flow cytometry analysis with NP-associated fluorescence detected in the APC-A channel. (**A**, **B**) A virus concentration-dependent decrease in liposome-cell association is observed but not in (**C**, **D**) the vector control cells. (**E**) Summary of PCR profiling of *SLC46A3* in LOXIMVI cells. (**F**) Quantification of liposomal association with clonal T47D-*SLC46A3* knockouts generated using the CRISPR/Cas9 system. Three clonal populations were tested for the *SLC46A3* knockout to ensure a consistent phenotype was observed. Data is represented as the mean and standard deviation of six biological replicates. (**G**) Representative flow histograms of clonal populations treated with liposomal NPs for 24 h; with NP-associated fluorescence detected in the APC-A channel. Quantification of (**H**) liposomal, (**I**) PLGA, and (**J**) PS NP flow data shown in Figure 4E. (*, *p* < 0.05, Mann-Whitney) (**K**) Flow cytometry analysis reveals *SLC46A3*-related trends in liposomal NP association with T47D cells are maintained with and without serum present in cell culture medium. T47D cells were cultured either with or without 10% FBS and treated with liposomal NPs for 24 h. Flow cytometry was used to quantify the liposome-cell association extent, and median fluorescence intensity (MFI) values for the vector control and *SLC46A3* KO cells were divided by the MFI of the liposome-treated parental T47D cells. Significant increases in liposome association (*, *p* < 0.05, Mann-Whitney) were observed in *SLC46A3* KO in both groups. Data is represented as the mean and standard deviation of four biological replicates.

**Figure S17.**
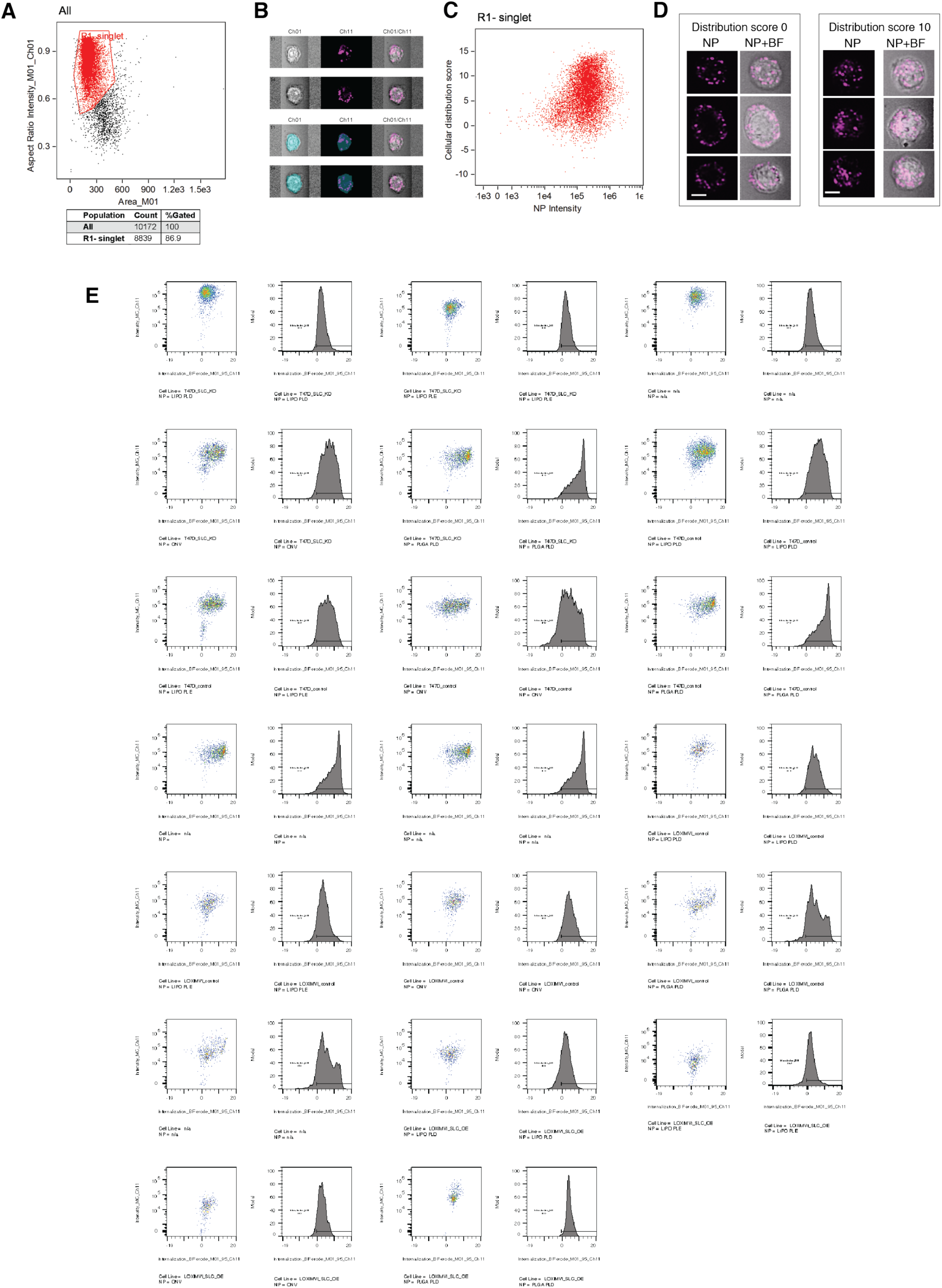
Imaging cytometry gating and raw data. (**A**) Gating for singlet cells (red) was based upon scatter plots using brightfield images (Ch01), with aspect ratio intensity on the y-axis and cell area on the x-axis. Gating data is shown for T47D-SLC46A3 knockout cells treated with LIPO-PLD; the same gate was used for all samples. (**B**) Example of singlet cells in the brightfield channel (Ch01) and Cy-5 NP channel (Ch11) with overlay. The bottom panel shows the same cells with the brightfield mask overlay in blue. (**C**) Using the brightfield mask, a cellular distribution score was generated using the built in ‘Internalization’ function in the IDEAS software. The relationship between the cellular distribution score (y-axis) and NP intensity (Cy5 signal, x-axis) is shown for T47D-SLC46A3 knockout cells treated with LIPO-PLD. (**D**) Representative cell images for a cellular distribution score = 0 (peripherally skewed) and =10 (homogeneous distribution) are shown from (C). (**E**) Raw data for all formulations (n=4) and cell lines (n=4) tested with dot plots of NP intensity on the x-axis and cellular distribution score on the y-axis and histograms of the cellular distribution score annotated with % of cells with internalized NPs.

**Figure S18.**
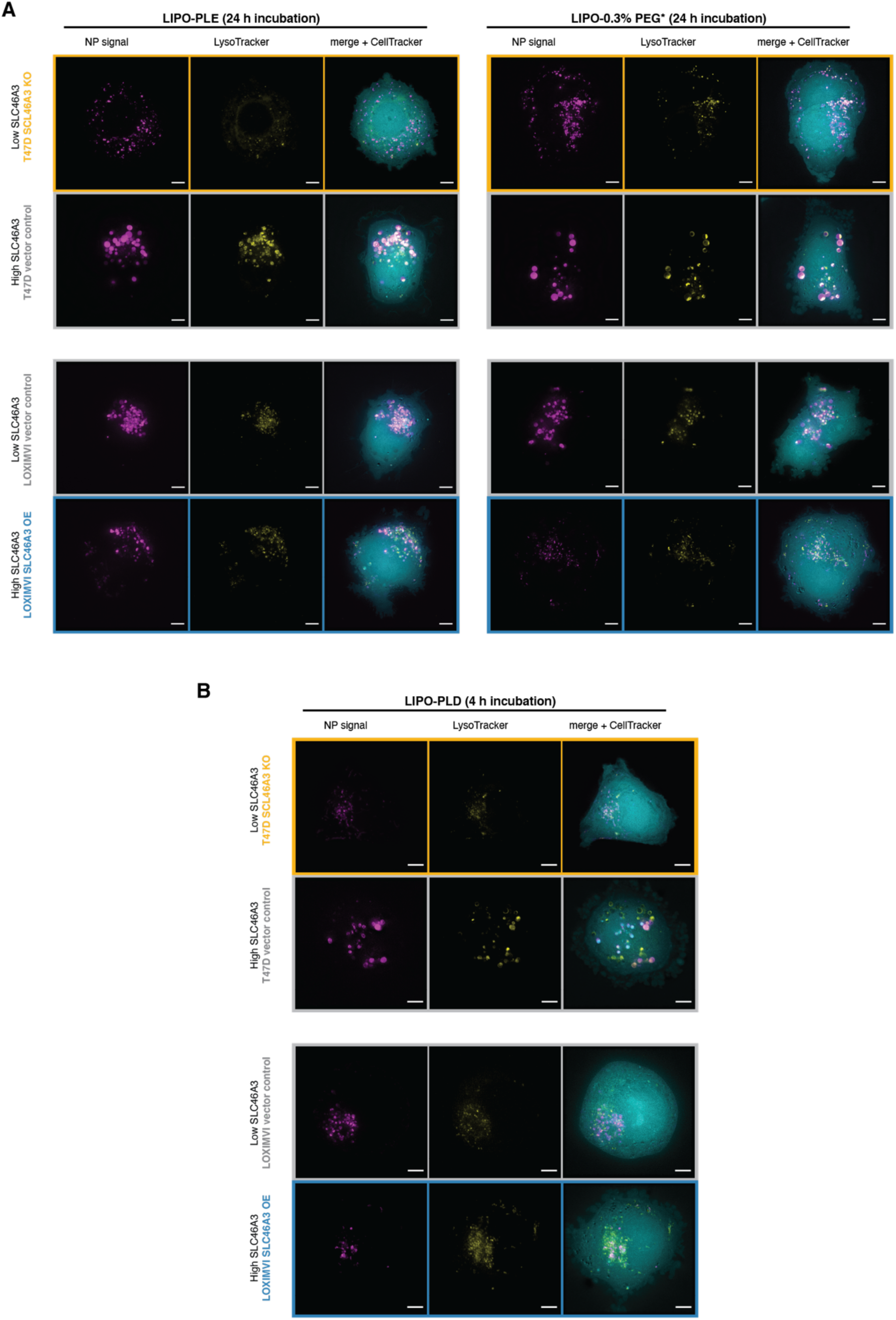
Liposomal nanoparticles exhibit SLC46A3-dependent changes in intracellular trafficking. Live cell micrographs of (**A**) LIPO-PLE and LIPO-0.3% PEG* NPs incubated with engineered T47D and LOXIMVI cells for 24 h. (**B**) Live cell micrographs of LIPO-PLD NPs incubated with T47D and LOXIMVI cells for 4 h. NP signal is pseudo-colored magenta, LysoTracker signal yellow, and CellTracker cyan. Scale bar = 5 μm.

**Figure S19.**
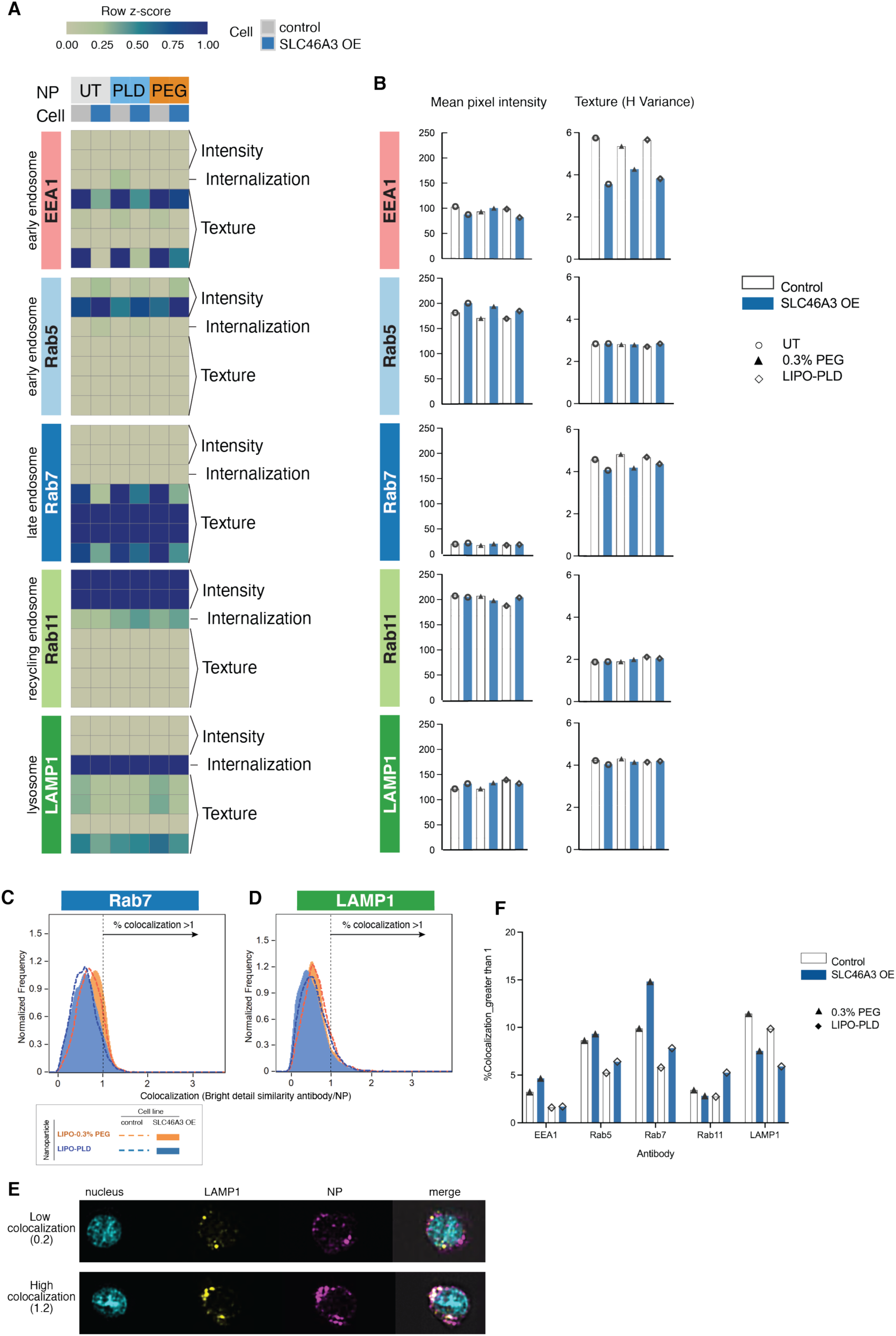
Imaging cytometry analysis of endolysomal markers and NP uptake as a function of SLC46A3 expression. (**A**) Heatmap showing metrics of endolysosomal marker intensity, internalization, and texture using imaging cytometry data for engineered LOXIMVI cells separately stained with five endolysosomal markers, both in the absence and presence of NPs. Cells were treated with NPs for 24 h with two separate formulations: LIPO-PLD (PLD) and LIPO-0.3% PEG (PEG). (**B**) Quantification of one measure of intensity (mean pixel intensity) and one measure of texture (H variance) for each column of the heatmap. (**C**) Colocalization scores of endolysomal marker and NP signals were generated by quantifying bright signal detail of these channels (bright detail similarity metric). Colocalization histograms of the two most variable endolysomal markers with LIPO-PLD and LIPO-0.3 mol%PEG are shown in (**C**) and (**D**) for Rab7 and LAMP1, respectively. (**E**) Representative images of LOXIMVI-control cells treated with LIPO-PLD and stained for LAMP1 at low and high colocalization. (**F**) Quantification of colocalization data for each marker-NP-cell combination. A higher colocalization score indicates a higher degree of colocalization between NP and marker signals.

**Figure S20.**
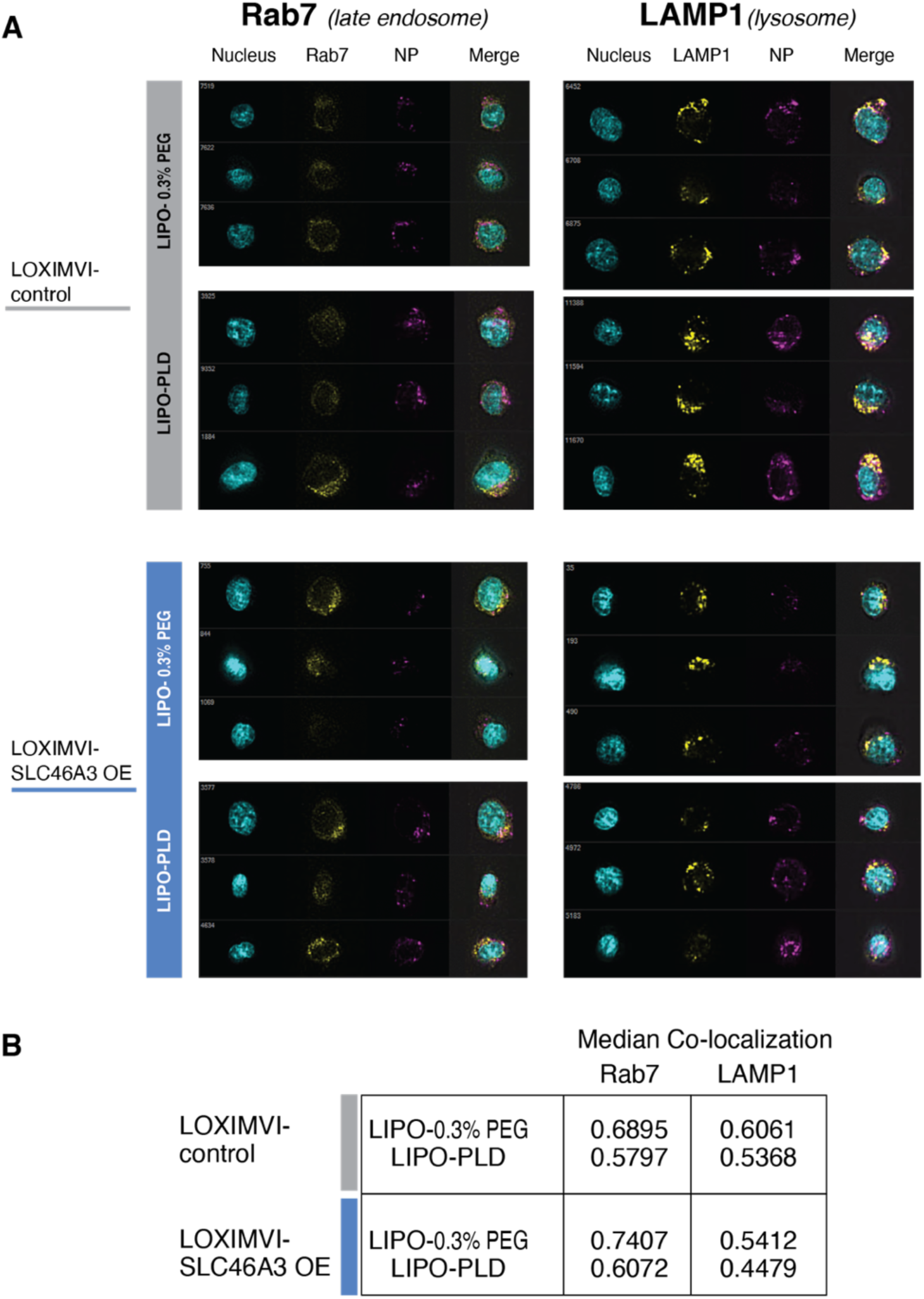
Representative cell images at median colocalization score. A) Images corresponding to the median bright field similarity between antibody and nanoparticle, raw histograms shown in S19, cells are shown with nuclear signal in cyan, antibody signal (Rab7, left; and LAMP1, right) in yellow, and nanoparticle signal in magenta). B) Quantified median co-localization scores for images shown in A. LOXIMVI-SLC46A3 OE cells have a higher median score for Rab7 co-localization and a lower score for LAMP1 co-localization with both liposomes tested (see also S19F).

**Figure S21.**
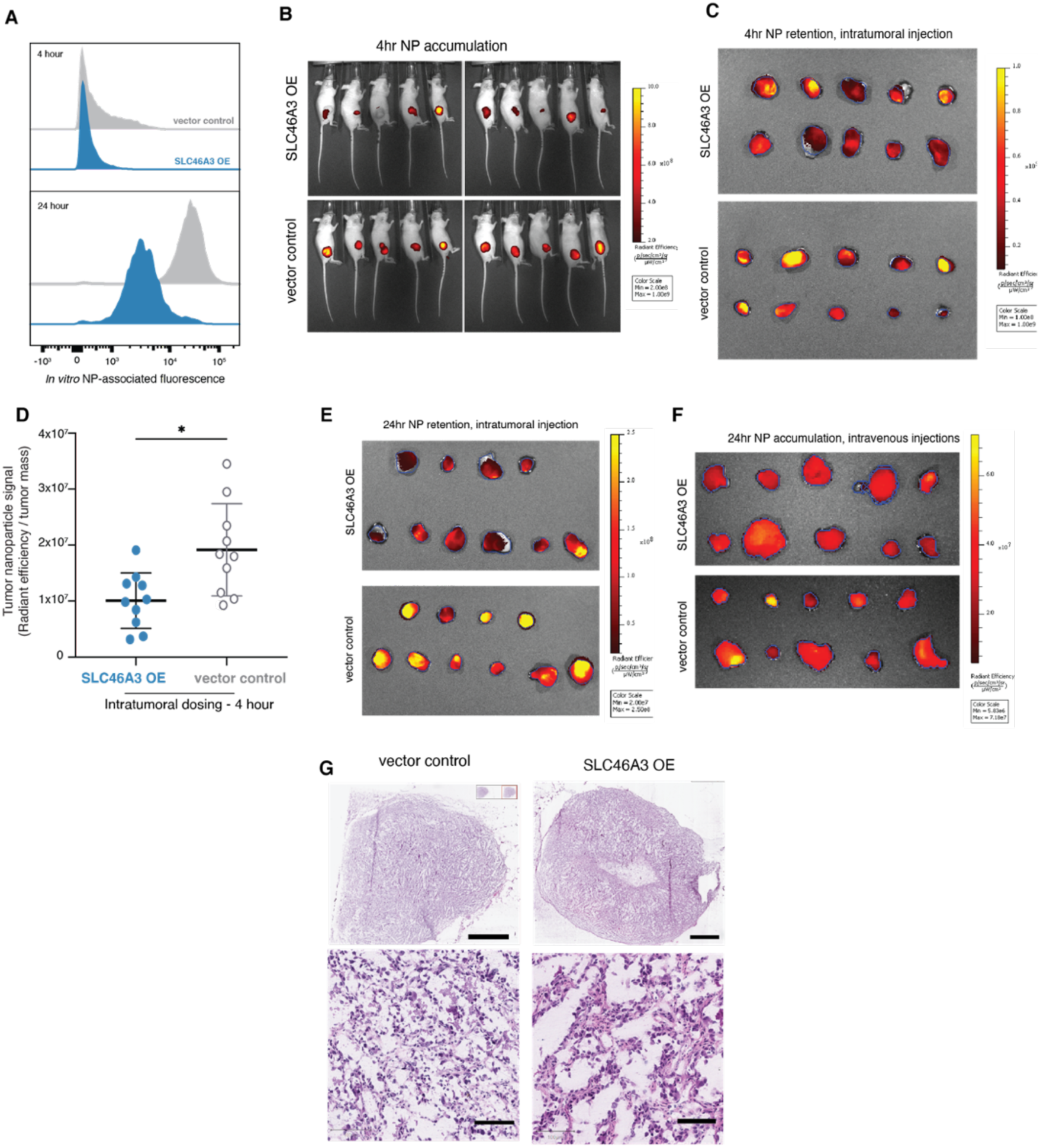
Raw data and histology images for *in vivo* studies. (**A**) Flow cytometry histograms for LIPO-0.3% PEG* (drug-free analog of liposomal irinotecan) in LOXIMVI cells overexpressing SLC46A3 (SLC46A3 OE) or luciferase (vector control) after 4 and 24 hours incubation wth NPs. (**B**) *In vivo* fluorescence images of mice bearing LOXIMVI flank tumors (n=10/group, 5 male and 5 female) 4 hours after intratumoral injection with LIPO-0.3% PEG* NPs. (**C**) Radiant efficiency from *ex vivo* tumors quantified in (**D**). (**E-F**) Radiant efficiency from *ex vivo* tumors used to generate plots in main manuscript Fig. 6C (**E**) and Fig 6D (**F**). (**G**) H&E staining of LOXIMVI tumor cross-sections show grossly similar morphology. Top two panels, scale bar = 2 mm; bottom two panels, scale bar = 0.1 mm.

**Figure S22.**
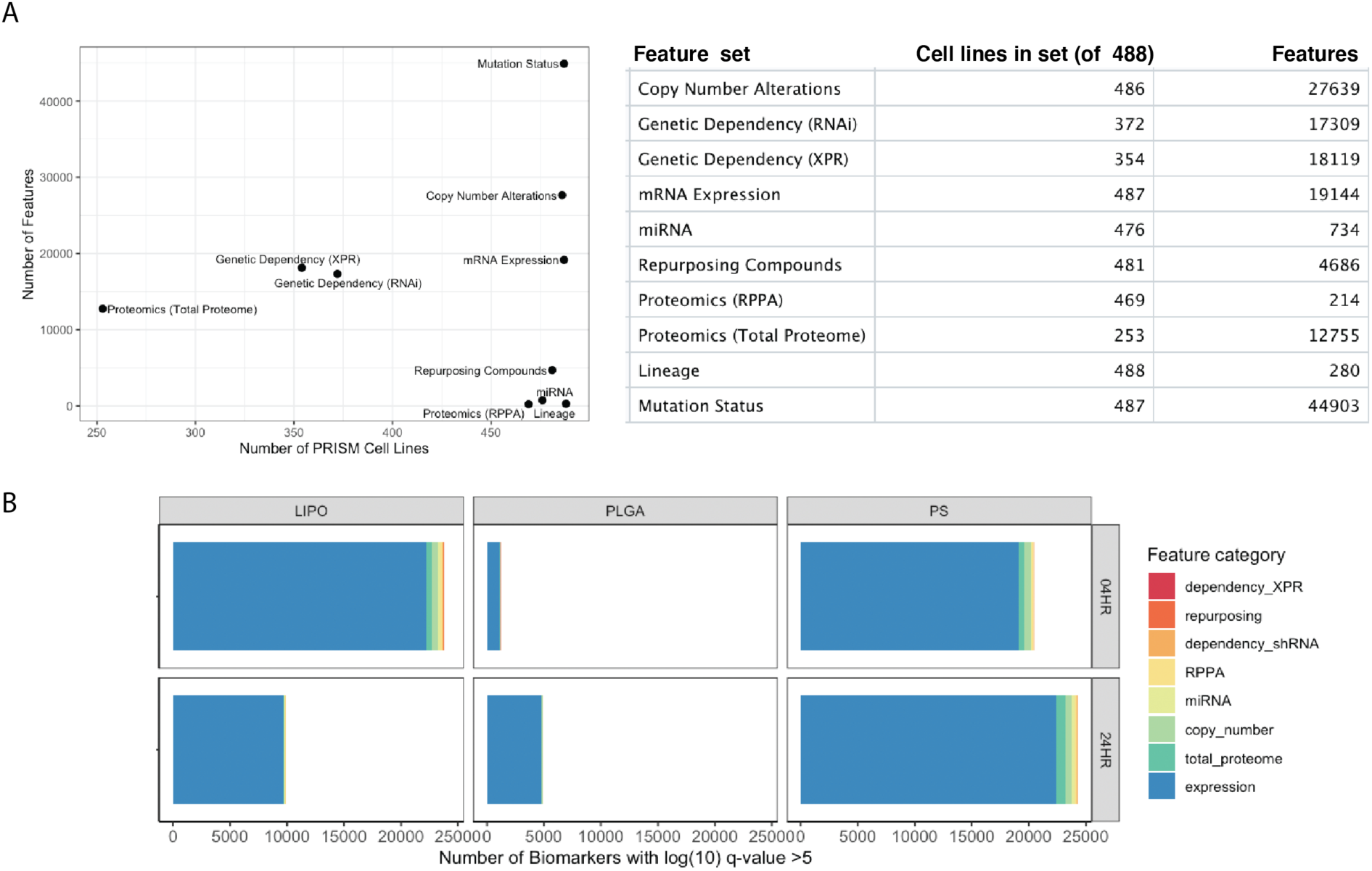
Ten feature sets were used in the analysis. (**A**) The cell lines and number of features per set are indicated. (**B**) Quantity of biomarkers with -log(q-value) > 5 per NP core are color coded by corresponding feature category. The majority of significantly associated features are from the expression data set.

**Table S1.**
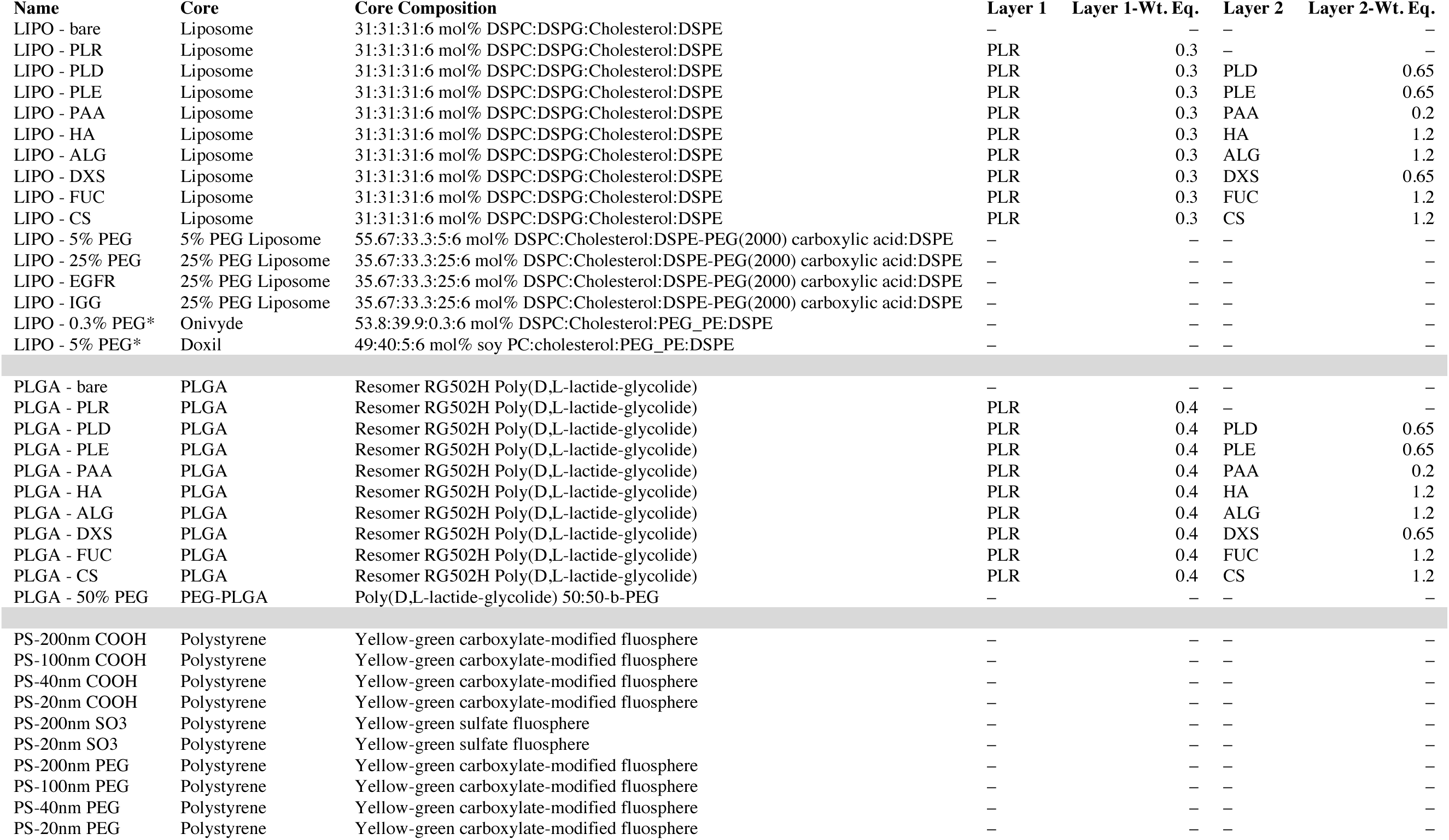
Nanoparticle formulation summary. Core compositions, polyelectrolyte identities and amounts used in the synthesis of the NP library are provided.

**Table S2.**
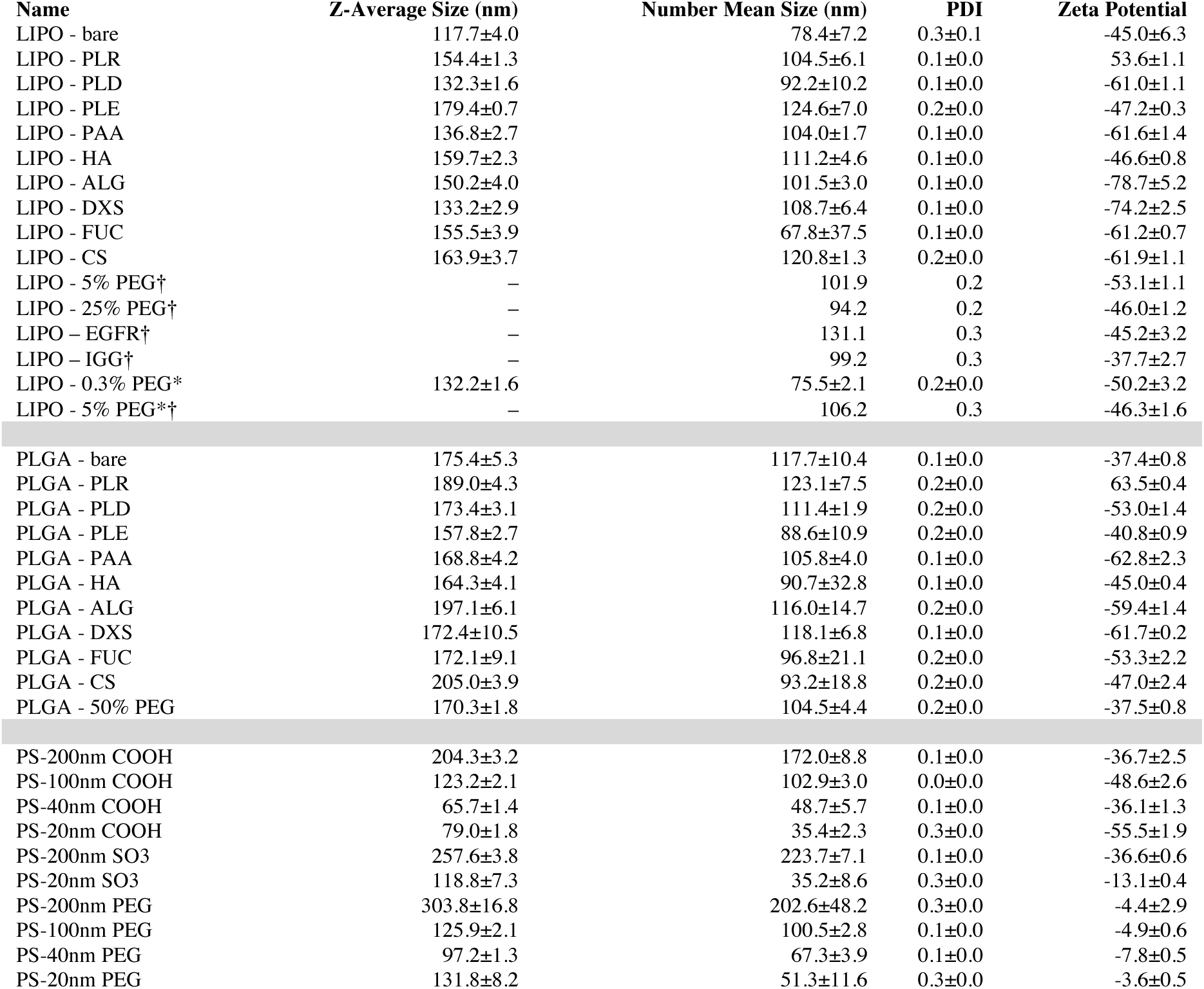
NP size, uniformity, and charge were measured using dynamic light scattering (DLS). Data is represented as the mean and standard deviation of three technical repeats, with the exception of formulations marked with a dagger (†). Size and PDI of these formulations were characterized using the Wyatt Dyna Pro Plate Reader, and only a single value for those measurements is presented.

**Table S3.** Fluorescence quantification of dye-labeled nanoparticles used in nanoPRISM screen and validation studies. Measurements were taken in flat black, 96 well plates (Nunclon). Nanoparticle solutions at the noted concentrations were diluted as follows: 10 μL into 90 μL dimethylsulfoxide (lipo and PLGA) or water (PS) prior to measuring at the noted settings. For LNP, sample was not diluted prior to fluorescence measurement.

| Name | Dye | Ex | Em | RFU | Gain ($\mu\text{m}$) | Z position | Est. Conc. (mg/mL) |
| --- | --- | --- | --- | --- | --- | --- | --- |
| Lipo - bare | sulfoCy5 | 640 | 690 | 6405 | 110 | 18350 | 0.125 |
| Lipo - PLD | sulfoCy5 | 640 | 690 | 4843 | 110 | 18350 | 0.1 |
| Lipo - PLE | sulfoCy5 | 640 | 690 | 4421 | 110 | 18350 | 0.1 |
| Lipo - HA | sulfoCy5 | 640 | 690 | 2445 | 110 | 18350 | 0.1 |
| Lipo - DXS | sulfoCy5 | 640 | 690 | 4973 | 110 | 18350 | 0.1 |
| Lipo - FUC | sulfoCy5 | 640 | 690 | 5730 | 110 | 18350 | 0.1 |
| Lipo - ALG | sulfoCy5 | 640 | 690 | 2355 | 110 | 18350 | 0.1 |
| Lipo - PAA | sulfoCy5 | 640 | 690 | 5634 | 110 | 18350 | 0.1 |
| Lipo - CS | sulfoCy5 | 640 | 690 | 5850 | 110 | 18350 | 0.1 |
| Lipo - 5% PEG | sulfoCy5 | 640 | 690 | 3528 | 110 | 18350 | 0.1 |
| Lipo - 25% PEG | sulfoCy5 | 640 | 690 | 1061 | 110 | 18350 | 0.1 |
| Lipo - EGFR | sulfoCy5 | 640 | 690 | 5693 | 110 | 18350 | 0.5 |
| Lipo - IGG | sulfoCy5 | 640 | 690 | 4734 | 110 | 18350 | 0.5 |
| Lipo - 0.3% PEG* | sulfoCy5 | 640 | 690 | 1352 | 110 | 18350 | 0.1 |
| Lipo - 5% PEG* | sulfoCy5 | 640 | 690 | 1443 | 110 | 18350 | 0.1 |
| PLGA - bare | Cy5 free acid | 640 | 690 | 2354 | 110 | 18350 | 0.1 |
| PLGA - PLD | Cy5 free acid | 640 | 690 | 3397 | 110 | 18350 | 0.1 |
| PLGA - PLE | Cy5 free acid | 640 | 690 | 3453 | 110 | 18350 | 0.1 |
| PLGA - PAA | Cy5 free acid | 640 | 690 | 3450 | 110 | 18350 | 0.1 |
| PLGA - HA | Cy5 free acid | 640 | 690 | 3276 | 110 | 18350 | 0.1 |
| PLGA - ALG | Cy5 free acid | 640 | 690 | 3405 | 110 | 18350 | 0.1 |
| PLGA - DXS | Cy5 free acid | 640 | 690 | 3351 | 110 | 18350 | 0.1 |
| PLGA - FUC | Cy5 free acid | 640 | 690 | 3300 | 110 | 18350 | 0.1 |
| PLGA - CS | Cy5 free acid | 640 | 690 | 3373 | 110 | 18350 | 0.1 |
| PLGA - 50% PEG | Cy5 free acid | 640 | 690 | 768 | 110 | 18350 | 0.1 |
| PS - 200 nm COOH | Yellow-green | 488 | 530 | 4168 | 72 | 18350 | 0.1 |
| PS - 100 nm COOH | Yellow-green | 488 | 530 | 4230 | 72 | 18350 | 0.1 |
| PS - 40 nm COOH | Yellow-green | 488 | 530 | 3538 | 72 | 18350 | 0.1 |
| PS - 20 nm COOH | Yellow-green | 488 | 530 | 3323 | 72 | 18350 | 0.1 |
| PS - 200 nm SO3 | Yellow-green | 488 | 530 | 5976 | 72 | 18350 | 0.1 |
| PS - 20 nm SO3 | Yellow-green | 488 | 530 | 3203 | 72 | 18350 | 0.1 |
| PS - 200 nm PEG | Yellow-green | 488 | 530 | 4053 | 72 | 18350 | 0.1 |
| PS - 100 nm PEG | Yellow-green | 488 | 530 | 4255 | 72 | 18350 | 0.1 |
| PS - 40 nm PEG | Yellow-green | 488 | 530 | 3457 | 72 | 18350 | 0.1 |
| PS - 20 nm PEG | Yellow-green | 488 | 530 | 3093 | 72 | 18350 | 0.1 |
| LNP1 | Cy5 | 640 | 690 | 1750 | 110 | 18350 | 0.22 |

**Table S4.** Information for antibodies used for endolysosomal staining.

| Antibody | Manufacturer | Item No | Lot | Purpose | Dilution |
| --- | --- | --- | --- | --- | --- |
| LAMP1 rabbit pAb | Abcam | Ab24170 | FR3409154 1 | Primary | 1:400 |
| Rab5 rabbit pAb | Thermo Fisher | PA5-29022 | XA3470699 | Primary | 1:100 |
| V5-tag rabbit mAb | Cell Signaling | D3H8Q | 6 | Primary | 1:400 |
| Rab11A |  | 71-5300 | WJ333110 | Primary | 1:100 |
| EEA1 rabbit mAb | Cell Signaling | C45B10 | 8 | Primary | 1:200 |
| Rab7 rabbit mAb | Cell Signaling | D95F2 | 3 | Primary | 1:50 |
| Anti-rabbit IgG Fab-AF555 conjugate | Cell Signaling | 4413S | 20 | Secondary | 1:1000 |

**Table S5.** Characterization of LNP formulations.

| Sample | Size (nm) | PDI | Zeta potential (mV) | Encapsulation efficiency |
| --- | --- | --- | --- | --- |
| LNP 1 | 94.8 ± 24.9 | 0.310 | 30.1 ± 1.4 | 82% |
| LNP 2 | 88.0 ± 3.0 | 0.190 | 35.4 ± 2.3 | 93% |
| LNP 3 | 95.1 ± 3.7 | 0.335 | 26.8 ± 3.8 | 94% |

**Data S1. String enrichment summary**

**Data S2. Cell line information for 488 cancer cell lines included in PRISM screen; raw numerical data for heat maps and univariate analyses.**

